# A conserved HSF:miR169:NF-YA loop involved in tomato and Arabidopsis heat stress tolerance

**DOI:** 10.1101/2021.01.01.425064

**Authors:** Sombir Rao, Sarita Jha, Chandni Bansal, Apoorva Gupta, Celine Sorin, Martin Crespi, Saloni Mathur

## Abstract

Regulatory feedbacks are at the basis of different stress and developmental networks in plants. Here, we report that tomato and Arabidopsis plants improve their heat stress tolerance through Heat stress transcription factor (HSF)-mediated transcriptional regulation of *MIR169* and post-transcriptional regulation of *NF-YA* transcription factors. We show that HSFs recognize tomato and Arabidopsis *MIR169* promoters using yeast-one-hybrid/ChIP-qPCR. Silencing tomato HSFs using virus-induced-gene-silencing (VIGS) reduce *Sly-MIR169* levels and enhance *Sly-NF-YA9/A10* target expression. Further, tomato transgenic plants overexpressing *Sly-MIR169* and *Sly-NF-YA9/A10-VIGS* knock-down tomato plants as well as Arabidopsis plants overexpressing *At-MIR169d* and *At-nf-ya2* mutants showed a link with increased heat tolerance. In contrast, Arabidopsis plants overexpressing *At-NF-YA2,* or those expressing a non-cleavable *At-NF-YA2* form (miR169-resistant *At-NF-YA2*) as well as plants inhibited for At-miRNA169d regulation (miR169d mimic plants) were more sensitive to heat stress, highlighting *NF-YA* as negative regulator of heat tolerance. Furthermore, post-transcriptional cleavage of *NF-YA* by elevated miR169 levels result in alleviating the repression of heat stress effectors HSFA7a/b in tomato and Arabidopsis revealing a retroactive control of HSFs by the miR169:NF-YA node. Hence, a regulatory feedback loop involving HSFs, miR169s and NF-YAs plays a critical role in the regulation of heat stress response in tomato and Arabidopsis plants.

## Introduction

Heat stress (HS) adversely affects the distribution and productivity of agriculturally important plants worldwide by affecting all aspects of plant processes like germination, growth, development, reproduction and yield (Mittler et al. 2010; Lobell et al. 2011; Hasanuzzaman et al. 2013). In nature, such HS conditions may be chronic or recurring, or both (Bäurle 2016); therefore, plants have evolved a variety of responses to elevated temperatures that minimize damage and ensure protection of cellular homeostasis.

HSFs are important regulators that play critical roles in signal transduction processes to mediate gene expression in response to multiple abiotic stresses, including cold, drought, salt, and heat (Collins et al. 1995; Nover et al. 2001; Guo et al. 2016; Jacob et al. 2017). HSFs recognize heat stress elements (HSE: 5′-GAAnnTTC-3′) that are conserved in promoters of heat stress-responsive (HSR) genes including HSP genes (Busch et al. 2005). During the initial stages of HS, HSFs are the primary molecules responsible for relaying signals of cellular stress to the transcriptional apparatus and reprogram gene expression to repair protein damage through elevated synthesis of molecular chaperones and proteases (Morimoto 2002; Scharf et al. 2012; Guo et al. 2016). In plants, there are multiple genes coding for HSFs for example, Arabidopsis has 21 HSF family members (von Koskull-D^ring et al. 2007) and tomato has 26 HSFs (Yang et al. 2016). Another class of regulatory molecules, the microRNAs (miRNAs) have emerged as a major class of small non-coding RNAs with roles in plant growth and development as well as under stress (Sunkar and Zhu, 2004; Lin et al. 2018; Ravichandran et al. 2019). HSF-mediated induction of MIR398 has been shown to trigger a regulatory loop that is critical for thermotolerance in Arabidopsis (Guan et al. 2013). Yan et al. (2012) have shown that stress-induced alternative splicing of MIR400 provides a mechanism for the regulation of miRNA processing in Arabidopsis during HS. Lin et al. (2018) have shown that miR160 modulates plant development and heat shock protein gene expression to mediate heat tolerance in Arabidopsis. Heat-responsive miRNAs in *Apium graveolens* (Li et al. 2014), switch grass (Hivrale et al. 2015), banana (Vidya et al. 2018) and apple (Niu et al. 2019) have also been reported.

The miR169 family members have significant roles in plant abiotic and biotic stress responses and are involved in various aspects of plant development including root architecture, nodule formation etc. by targeting *Nuclear factor-Y A* (NF-YA) gene family members (Zhao et al. 2011; Ni et al. 2013; Sorin et al. 2014; Zhang et al. 2015; Li et al. 2017). Tomato plants overexpressing miR169 show improved resistance to drought stress (Zhang et al. 2011) while in Arabidopsis enhanced miR169 levels promote leaf dehydration (Li et al. 2008; Ni et al. 2013). Over-accumulation of miR169 is correlated with a reduction of *NF-YA* target transcripts in Arabidopsis seedlings under cold stress (Zhou et al. 2008; Lee et al. 2010). Role of miR169 is also established in long-distance signalling and nitrogen/phosphorus starvation (Pant et al. 2009; Buhtz et al. 2010; Zhao et al. 2011). In a previous publication from our laboratory, we have reported 18 *MIR169* family members in tomato (Rao et al. 2020). Assessment of *Sly-MIR169* precursors in different heat stress (HS) regimes highlighted that fifteen of the sixteen precursors (two of the loci are bicistronic) are differentially regulated in response to HS. Heat-mediated up-regulation of *MIR169*s has also been reported in Arabidopsis (Li et al. 2010; Guan et al. 2013) however, a detailed mechanism was lacking. Here, we have dissected the transcriptional regulation of *MIR169* family by HSFs in tomato and Arabidopsis HSR. Our laboratory has previously established miR169-mediated transcript cleavage of five classical targets belonging to the *NF-YA* class of transcription factors (*Sly-NF-YA1, Sly-NF-YA3, Sly-NF-YA8, Sly-NF-YA9 and Sly-NF-YA10*) (Rao et al. 2020). Here, we show that miR169 mediated post-transcriptional down-regulation of *At-NF-YA2* in *Arabidopsis* and *Sly-NF-YA9* and *Sly-NF-YA10* in tomato enhances HS tolerance and impacts HSF expression. NF-YA transcriptionally regulates a suite of HSR genes including HSFs which exert a feedback regulation on *NF-YAs* through the control of miR169 expression. This is a new feedback loop involved in the regulation of plant tolerance to HS.

## Results

### *MIR169* members are transcriptionally regulated by HSFs in tomato

The transcriptional activation of heat stress genes is regulated by HSFs that recognize HSE cis-elements in promoters of HSR genes (Busch et al. 2005). All the 15 heat up-regulated *Sly-MIR169* promoters (Rao et al. 2020), contain one or more HSEs indicating possible involvement of HSFs in their heat inducibility (Supplementary figure S1, Supplementary table S1). This prompted us to investigate the transcriptional regulation of *MIR169* promoters by HSFs using yeast-one-hybrid (Y1H) assays. Out of 322 interactions studied between tomato HSFs and 14 heat-responsive *Sly-MIR169* promoters (one promoter could not be assessed as a minimal inhibitory Aureobasidin A antibiotic concentration could not be achieved), we found specific binding of seven HSFs on five *Sly-MIR169* promoters (Figure 1A, Supplementary table S2, Supplementary figure S2). Promoters of *Sly-MIR169a-1* and *Sly-MIR169d-1* are regulated by a single HSF viz., Sly-HSFA1e and Sly-HSFA1a, respectively (Figure 1A, Supplementary table S2). *Sly-MIR169b* promoter interacts with Sly-HSFA1a and Sly-HSFA7a while *Sly-MIR169c* promoter shows Sly-HSFA4a and Sly-HSFA6b binding. Transcription of *Sly-MIR169d* is regulated by Sly-HSFA8a and Sly-HSFB1a (Figure 1A).

**Figure 1:**
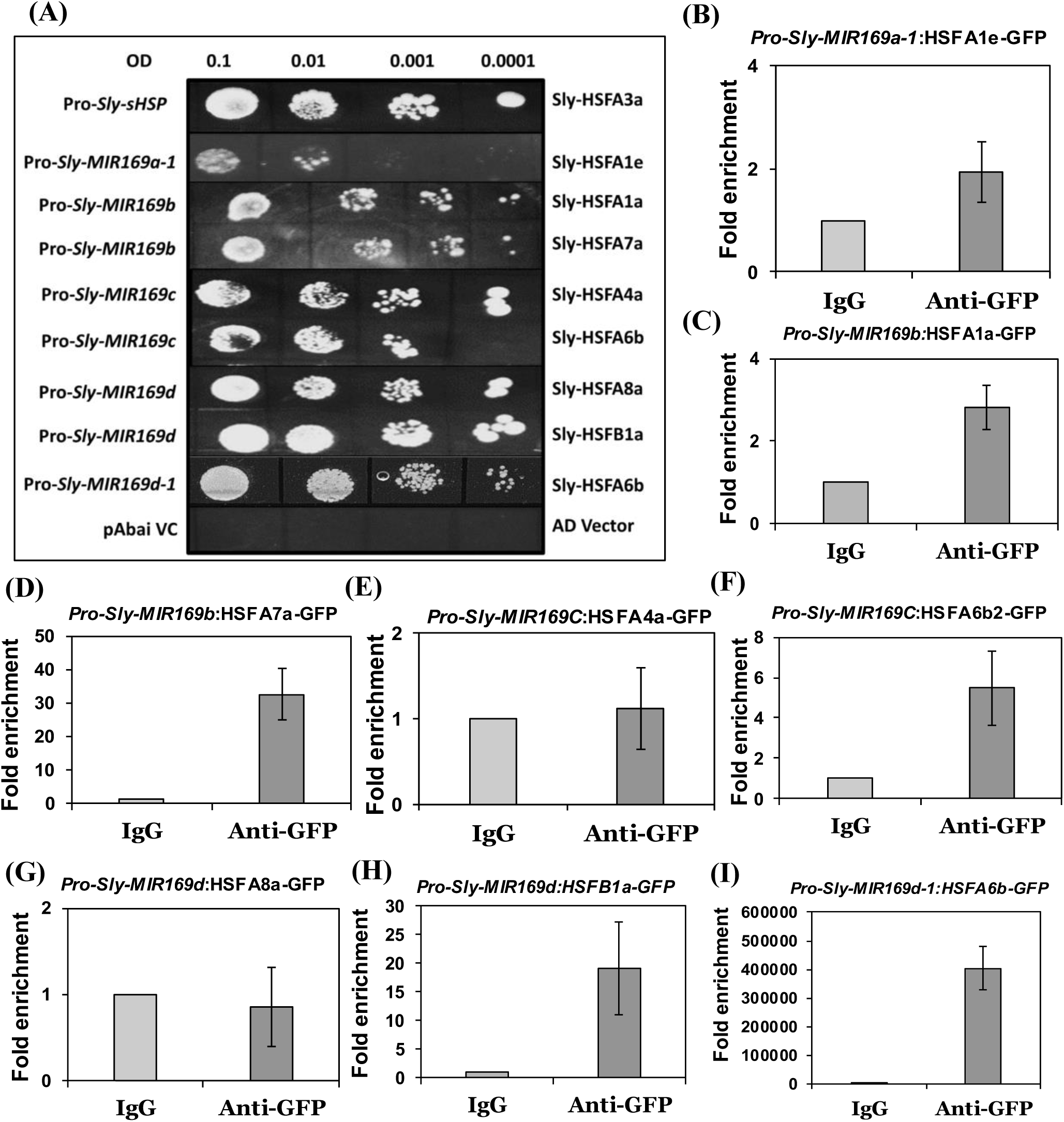
Yeast one hybrid and Chromatin immunoprecipitation (ChIP-qPCR) assay of Sly-MIR169 promoters and HSFs. (A) Yeast-one-hybrid assay to assess transcriptional regulation of heat-responsive *MIR169* promoters by HSFs. Only the positive Y1H interactions between different HSFs and *Sly-MIR169* promoters have been shown. The *Sly-MIR169* promoters were cloned in the reporter vector pABAi and the full-length cDNA of HSFs were cloned in the pGADT7 vector. After the co-transformation into Y1H gold yeast strain, serial dilutions (O.D. 0.1 to 0.0001) of yeast cultures were spotted onto plates lacking URA and LEU with specific Aureobasidin A concentration and incubated for 3 days. Promoter of *sHSP* has been used as a positive control to show binding of HSFA3a on *sHSP* promoter (Li et al. 2013) and empty vectors as negative controls. The Aureobasidin A concentrations for all *Sly-MIR169* promoters is presented in supplementary table S7. (B-I) Chromatin immunoprecipitation (ChIP-qPCR) assay of *Sly-MIR169* promoters and HSFs in *Nicotiana benthamiana*. Fold enrichment of *Sly-MIR169a-1* promoter in HSFA1e immunoprecipitated sample (B); Fold enrichment of *Sly-MIR169b* promoter in HSFA1a immunoprecipitated sample (C); in HSFA7a immunoprecipitated sample (D); Fold enrichment of *Sly-MIR169C* promoter in HSFA4a immunoprecipitated sample (E); Fold enrichment of *Sly-MIR169C* promoter in HSFA6b immunoprecipitated sample (F); Fold enrichment of *Sly-MIR169d* promoter in HSFA8a immunoprecipitated sample (G); in HSFB1a immunoprecipitated samples (H); Fold enrichment of *Sly-MIR169d-1* promoter in HSFA6b immunoprecipitated samples (I). These experiments were repeated 3-4 times; the error bars represent standard deviation between the replicates. The values for different promoters in IgG immunoprecipitated sample were set as one for normalization.

Further, to confirm the binding of these Sly-HSFs on *Sly-MIR169* promoters, we performed chromatin immunoprecipitation (ChIP) assays by co-expressing specific *Sly-MIR169* promoters and *P35S:HSF:GFP* constructs in *Nicotiana benthamiana* leaves. ChIP-qPCR was then performed on the HSE-specific regions in the *Sly-MIR169* promoters. Fold enrichment was calculated by comparing the Ct values of anti-GFP immunoprecipitated samples with IgG immunoprecipitation control (Figure 1B-I). ChIP-qPCR analysis of all HSFs with *Sly-MIR169* promoters matched the Y1H results except for HSFA4a and HSFA8a on *Sly-MIR169c* and *Sly-MIR169d* promoters (Figure 1E, G).

The HSF-mediated regulation of *MIR169* promoters was also assessed in *Nicotiana benthamiana* leaves using agro-infiltration (Supplementary Figure S3). For this, *Sly-MIR169* promoters were fused with *GUS* reporter gene and co-transformed with effector plasmids containing HSFs under CaMV35S promoter. All the five HSFs showing promoter enrichment in the ChIP assays were transformed individually or in combinations (as the case may be) to check their ability to drive *GUS* gene expression under respective *Sly-MIR169* promoters. qRT-PCR based estimation of *GUS* transcripts in presence of HSFs confirmed HSF-mediated activation of *Sly-MIR169* promoters for all the HSFs (Supplementary Figure S3) except Sly-HSFA1a (Supplementary Figure S3B). However, co-transformation of Sly-HSFA1a with Sly-HSFA7a (Supplementary Figure S3B) leads to enhanced levels of *GUS* transcripts as compared to individual HSFs suggesting a cooperative action in regulating *Sly-MIR169b* promoter.

### HSFs regulate the orchestration of *miR169:NF-YA* module during heat stress in tomato

To assess whether the HSF:*miR169:NF-YA* module is functional in tomato HS, we quantified the transcript abundance of all components of this module in wild type tomato plants upon HS. Expression profiles of HSFs reveal the heat-mediated up-regulation of *Sly-HSFA6b, Sly-HSFA7a* and *Sly-HSFB1a* while *Sly-HSFA1a* and *Sly-HSFA1e* are not HS responsive (Figure 2A, Supplementary figure S4A). HSFA1s, the master regulators of HSR response, are present in bound forms with HSPs in cytoplasm. Upon HS, HSFs dissociate from HSPs and move to nucleus to transcriptionally regulate HSR genes (Kotak et al. 2004; Chan-Schaminet et al. 2009) hence, HSFAs do not exhibit transcriptional up-regulation upon HS. However, we note that in non-stressed conditions the Sly-*HSFA1s* are more abundant in comparison to other HSFs (by comparing Ct values of these HSFs), suggesting their sufficient levels to orchestrate gene transcription upon HS (Supplementary table S3A). In agreement with our previous results (Rao et al. 2020), four miR169 mature forms (Figure 2B, Supplementary figure S4B) exhibit heat-mediated induction in comparison to non-stressed WT tomato plants. Comparison of their Ct values in control conditions revealed the abundant presence of miR169c in non-stressed plants, while imposition of heat stress led to strong induction of miR169a (Supplementary table S3B). Furthermore, expression analysis of miR169s targets (NF-YAs) shows reduction of only *Sly-NF-YA9* and *Sly-NF-YA10* transcripts in HS in WT plants (Figure 2C, Supplementary figure S4C).

**Figure 2:**
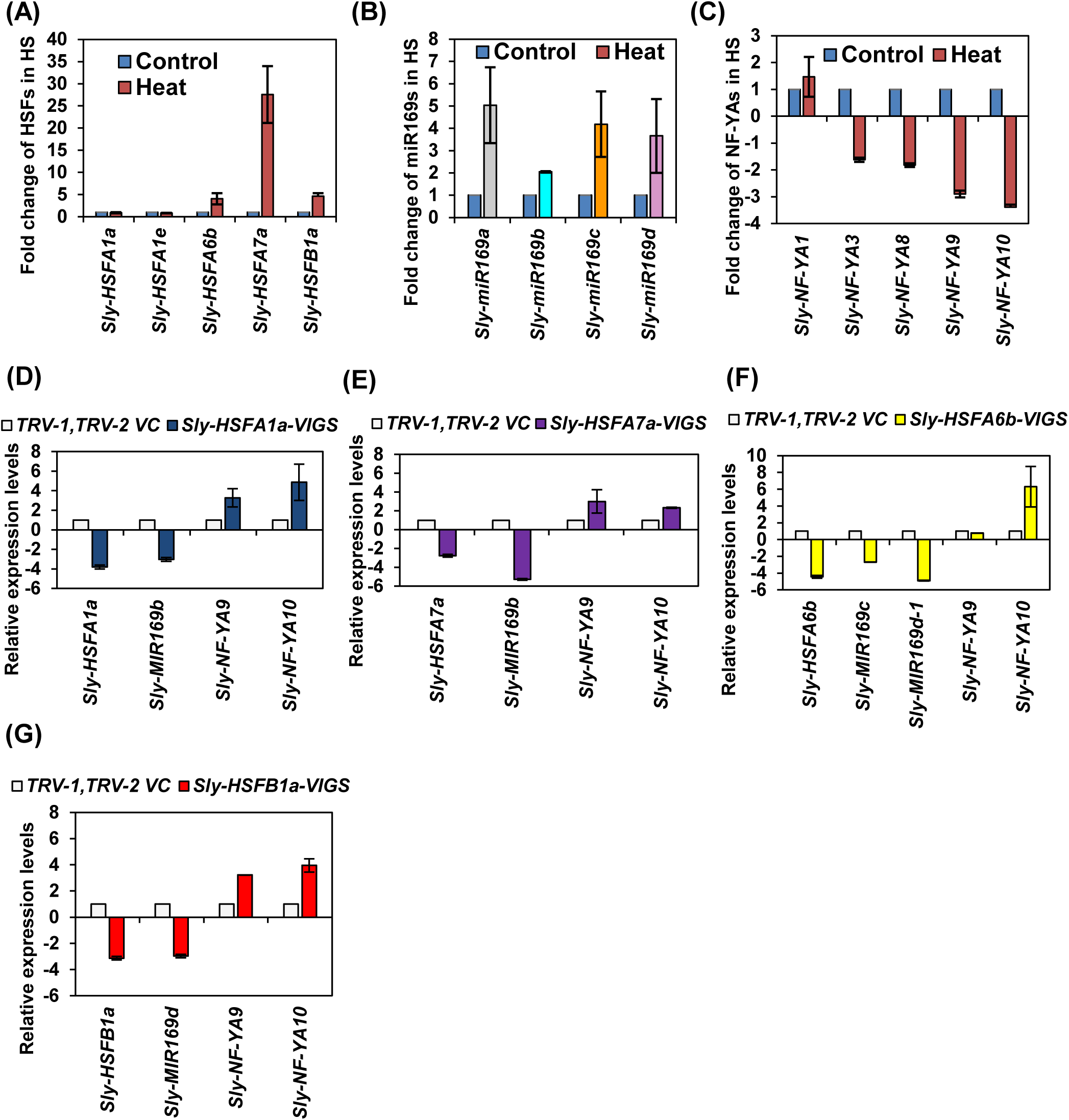
Validation of HSF:miR169:*NF-YA* node components in tomato upon heat stress. (A) Expression profiles of *HSF* genes that regulate *Sly-MIR169* transcription in WT tomato leaves during control and heat stress condition by qRT-PCR using the 2−ΔΔCt method. (B) Expression of mature miR169s in response to heat stress as determined by taqman based qRT-PCR in control and heat challenged WT tomato leaves. (C) qRT-PCR based expression profiles of target *NF-YA* transcripts in WT control and heat stressed tomato leaves using the 2−ΔΔCt method. (D-G) The expression analysis of *HSF* genes, *Sly-MIR169* precursors and *NF-YA* transcripts in VIGS plants silenced for *Sly-HSFA1a* (D), *Sly-HSFA7a* (E), *Sly-HSFA6b* (F) and *Sly-HSFB1a* (G). In (D-G) 15-days-old tomato plants were agro-infiltrated with empty vector (TRV-1, TRV-2 VC) or TRV-Sly-HSF VIGS constructs. The VIGS established plants were subjected to heat stress after 3-weeks-of agro-infiltration, and used for expression studies of different genes. Graphical data represents mean values of expression of three to four biological sets. Error bars show standard deviation. To represent the negative fold-change on y axis in (C-G), the relative expression values of down-regulated genes were transformed using the formula [-(1 / RQ ≤0.5)]. *Actin* was used as endogenous control in (A), (C) and (D-G), while *5SrRNA* was used as reference control in (B). The fold change normalization was done by setting the control value as one for all qRT-PCR expression analysis.

To investigate the direct role of HSF-mediated regulation of the *MIR169:NF-YA* module in tomato, we utilized TRV-based virus induced gene silencing (VIGS) of HSFs. The VIGS silencing assays were performed 3-weeks-post agro-infiltration. Setting up of efficient silencing for TRV-HSFs was assessed by the reduction in the endogenous HSFs transcripts in silenced plants (Figure 2D-G). To ensure the specific downregulation of selected HSFs, we have examined the expression profiles of putative off targets HSFs predicted by SGN-VIGS tool (Supplementary table S4) by qRT-PCR. The unaltered expression of putative off targets in different HSFs-VIGS silenced plants confirms the specificity of the silencing (Supplementary figure S5A-C). Subsequently, downstream components in the pathway like *MIR169* precursor levels and their heat responsive *NF-YA* targets (*Sly-NF-YA9* and *Sly-NF-YA10*) were quantified by qRT-PCR after these plants were given a HS (Figure 2D-G, Supplementary figure S6A-D). Silencing of *Sly-HSFA1a* and *Sly-HSFA7a* reduces the transcription of *Sly-MIR169b* by 3 and 5 fold respectively, and leads to more than 2 fold enhanced expression of *Sly-NF-YA9* and *Sly-NF-YA10* transcripts upon HS (Figure 2D, E). Transcription of *Sly-MIR169c* is reduced by 2.6 folds in *Sly-HSFA6b* silenced plants, this in turn leads to up-regulation of *Sly-NF-YA10* upon HS (Figure 2F). Silencing of *Sly-HSFB1a* (Figure 2G) reduces the expression of *Sly-MIR169d* by 2.9 folds, resulting in enhanced accumulation of *Sly-NF-YA9* and *Sly-NF-YA10* transcripts upon HS. *Sly-HSFA6b* silencing also reduces the levels of *Sly-MIR169d-1* by 4.9 folds that in-turn enhances the transcript levels of *Sly-NF-YA10* upon HS (Figure 2F). Depressed expression of *Sly-MIR169s* with concomitant enhanced expression of *Sly-NF-YA9/10* in *Sly-HSF*-silenced backgrounds upon HS suggests the HSF-mediated regulation of *Sly-MIR169:Sly-NF-YA9/10* module in HS.

### The miR169:NF-YA module is critical for high temperature tolerance in tomato

To establish the functions of Sly-*MIR169:Sly-NF-YA9* /*Sly-NF-YA10* module in heat tolerance, we raised *Sly-MIR169* overexpression (*Sly-MIR169*-OE) transgenic tomato lines and used the VIGS-silencing approach to *knock*-down *Sly-NF-YA9*/*A10* expression in tomato plants. qRT-PCR based expression analysis exhibited higher accumulation of *Sly-MIR169* precursor and significant reduction for *Sly-NF-YA9* and *Sly-NF-YA10* target transcripts in *Sly-MIR169*-OE lines as compared to wild-type control plants (Figure 3A). Furthermore, promoter reporter transgenic line analysis for *pro-Sly-MIR169* and *pro-Sly-NF-YA10* revealed the complementary expression patterns of the miRNA and target confirming the validity of this module in tomato leaves and roots (Figure 3B). Seedling growth measurement and thermotolerance assays of 4-days-old *Sly-MIR169*-OE transgenic seedlings exhibited enhanced hypocotyl length and survival rate as compared to WT seedlings in non-stressed as well as upon experiencing heat stress (Figure 3C, D). The survival rate of *Sly-MIR169-OE* overexpressing seedlings was twice that of the WT plants (Figure 3E) Also, the imposition of HS impaired the growth of WT seedlings as is evident from ∼50% reduction in hypocotyl length during HS as compared to the WT seedlings kept in the non-stressed conditions (Figure 3F). On the contrary, the *Sly-MIR169-OE* transgenic seedlings did not exhibit any significant arrest in hypocotyl length during HS in comparison to the non-stressed *Sly-MIR169-OE* seedlings (Figure 3F), confirming the positive regulatory roles of *Sly-MIR169* in tomato thermotolerance. Taken together, the data suggests that *Sly-MIR169* act as positive regulator of heat tolerance. To further explore the biochemical and physiological mechanisms involved in governing the higher thermotolerance of *Sly-MIR169-OE* lines, we quantified the relative water content (RWC), electrolyte leakage (EL) and proline content in the leaves of the 30-days-old *Sly-MIR169-OE* and WT tomato. RWC measurement of WT and *Sly-MIR169-OE* plants during non-stressed and HS conditions revealed a strong reduction in RWC of WT during HS conditions (Supplementary figure S7), while very slight reduction was observed for *Sly-MIR169-OE* lines during HS as compared to the control *Sly-MIR169-OE* lines (Supplementary figure S7). Furthermore, the estimation of EL in WT and *Sly-MIR169-OE* plants revealed the considerable higher leakage of electrolyte in WT plants as compared to the *Sly-MIR169-OE* plants in HS conditions (Supplementary figure S7). Since, there is comparable EL levels between the WT and *Sly-MIR169-OE* control non-stressed plants, overexpression of *Sly-MIR169* imparts robust physiological status to the transgenic lines upon exposure to HS (Supplementary figure S7). This was further confirmed by DAB and Trypan blue staining of WT and *Sly-MIR169-OE* plants; a high degree of damage and cell death was observed in WT plants in comparison to *Sly-MIR169-OE* transgenic lines in HS (Supplementary figure S7). Moreover, the qRT-PCR based expression analysis of HSR genes in WT and *Sly-MIR169-OE* plants revealed the involvement of *Sly-sHSP17*.6-CII and *Sly-HSFA7* in maintaining the higher survival percentage of *Sly-MIR169-OE* plants (Supplementary figure S7).

**Figure 3:**
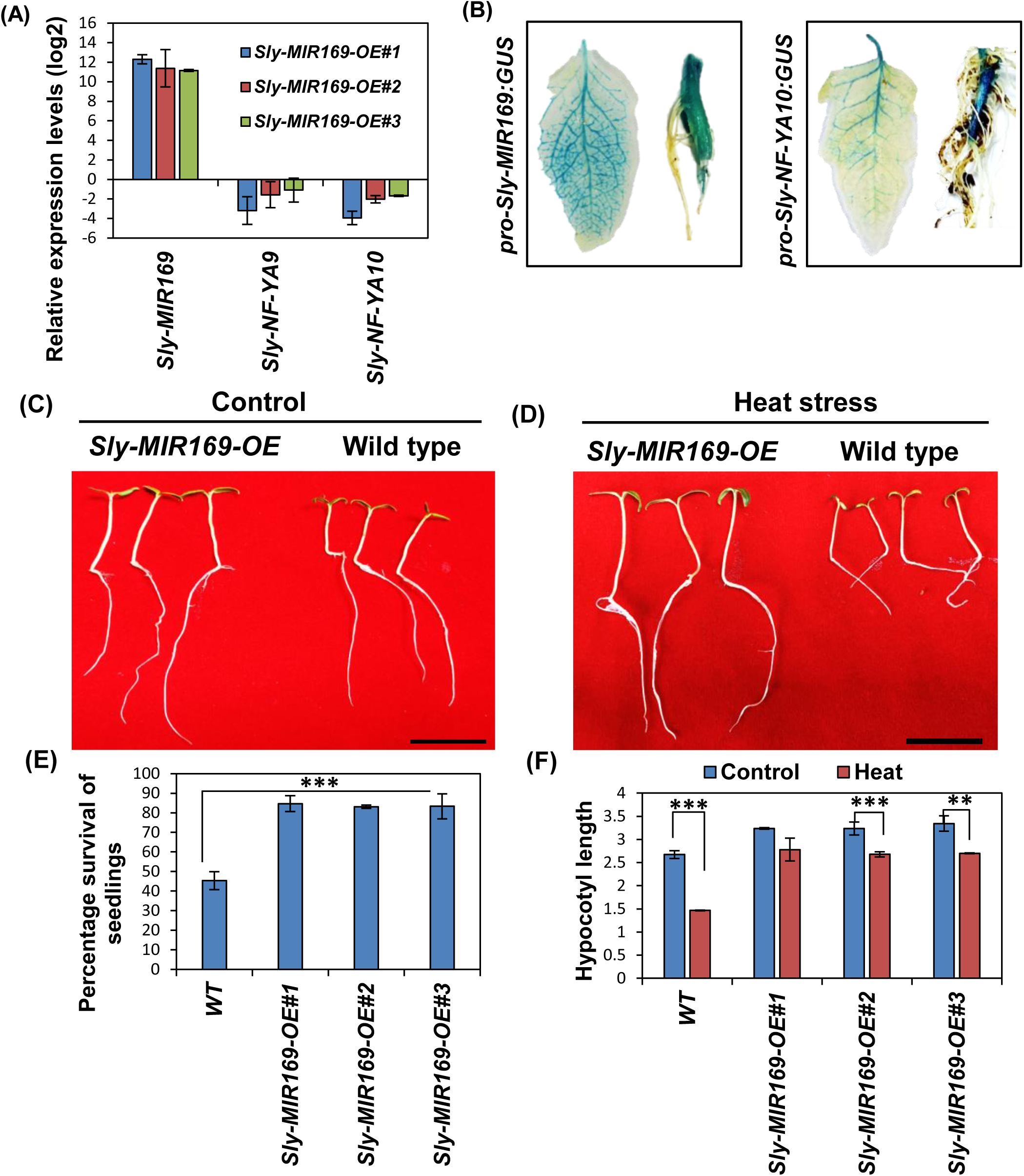
Functional evaluation of *Sly-MIR169* overexpression transgenic lines in tomato. (A) qRT-PCR based expression analysis of *Sly-MIR169* and *Sly-NF-YA9/A10* targets in transgenic *Sly-MIR169-OE* lines. The expression levels of genes were calculated using the 2−ΔΔCt method and presented using fold-change values. The fold change normalization was done by setting the WT value as one. Log2 transformation was applied on fold change values to obtained negative fold changes on y-axis. *Actin* was used as endogenous control. Graphical data represents mean values of expression of three biological sets Error bars represent the standard deviation of three independent biological replicates. (B) GUS signals showing temporal expression of *Sly-MIR169* and *Sly-NF-YA10.* The experiments were repeated at least three independent transgenic lines, and data from one representative line are shown. (C-D) Phenotypes of control (C) and heat challenged (D) WT and *Sly-MIR169-OE* seedlings. For survival assay 4 days old seedlings were exposed to 45^ᵒ^ C for 4.5 hours. The seedlings were photographed after 4 days of recovery post heat stress. Black line at the base of the picture represents 2cm scale. (E) Percentage survival estimation of WT and *Sly-MIR169-OE* lines after 4 days of heat stress at 45^ᵒ^ C for 4.5 hours. (F) Measurement of hypocotyl length of WT and *Sly-MIR169-OE* lines in control and heat stressed conditions. A sample size of 50 seedlings was used per replicate and the experiment was conducted with three independent *Sly-MIR169-OE* transgenic lines for Figure (E) and (F). Graphical data represents mean values of multiple sets. Error bars show standard deviation. *p<0.05, **p<0.01 and ***p<0.001 vs. wild type, by two-tailed Student’s t-test.

Furthermore, the heat mediated downregulation of *Sly-NF-YA9/A10* (Figure 2C) and their significantly reduced expression in *Sly-MIR169-OE* lines (Figure 3A), indicates the involvement of *Sly-NF-YA9/A10* as downstream component in regulating the heat specific phenotype of *Sly-MIR169-OE* lines. Therefore, to explore the involvement of *Sly-NF-YA9/A10* in regulating tomato thermotolerance, we performed virus induced silencing of *Sly-NF-YA9/A10.* A silencing efficiency of ∼80% was achieved for both *Sly-NF-YA9* and *Sly-NF-YA10* transcripts as compared to the TRV-vector negative control plants after three weeks of the agro-infiltration of the VIGS constructs (Supplementary Figure S8A). Expression analysis of Sly-*NF-YA9* or Sly-*NF-YA10* in WT control (non-stressed) and *TRV-EV* infiltrated plants revealed no significant fluctuation in their expression levels (Supplementary figure S8B), suggesting the specificity of the VIGS silencing constructs. The *TRV-Sly-NF-YA9* and *TRV-Sly-NF-YA10* plants were given a HS and scored for heat tolerance six-days-after HS. qRT-PCR based expression analysis revealed that a strong downregulation of *Sly-NF-YA9* and *Sly-NF-YA10* is maintained in VIGS silenced plants post HS (Supplementary figure S8C-D), ruling out any time point and treatment biased alteration in their expression levels. Phenotypic evaluation post-HS showed that the leaves in vector control plants are highly damaged in these conditions, in contrast to the *Sly-NF-YA9* and *Sly-NF-YA10* silenced plants which are robust with green, healthy, erect leaves and also show emergence of new leaves (Figure 4A). The *Sly-NF-YA9* and *Sly-NF-YA10* knock-down plants show ∼55% and 80% survival rate (judged by phenotype of 7^th^ to 9^th^ leaves and emergence of new leaves) in comparison to ∼33% for vector-control plants (Figure 4B). Further, we gauged the physiological performance of *TRV-Sly-NF-YA9* and *TRV-Sly-NF-YA10* silenced plants, six days after HS using a Li-Cor 6400 photosynthesis measuring system. Both the *NF-YA* silenced plants show significant enhancement in water use efficiency and net photosynthetic rates as compared to TRV-control plants under HS (Figure 4C). To rule out that the post HS phenotyping and survival scoring is not because of an effect of silencing of off targets, we predicted any possible off targets of *Sly-NF-YA9* and *Sly-NF-YA10* genes. Unaltered expression patterns of these putative off target genes in *Sly-NF-YA9* and *Sly-NF-YA10* VIGS silenced plants confirmed that the phenotype observed is specific to the knock-down of *Sly-NF-YA9* and *Sly-NF-YA10* genes (Supplementary table S4, Supplementary figure S9A-B).

**Figure 4.**
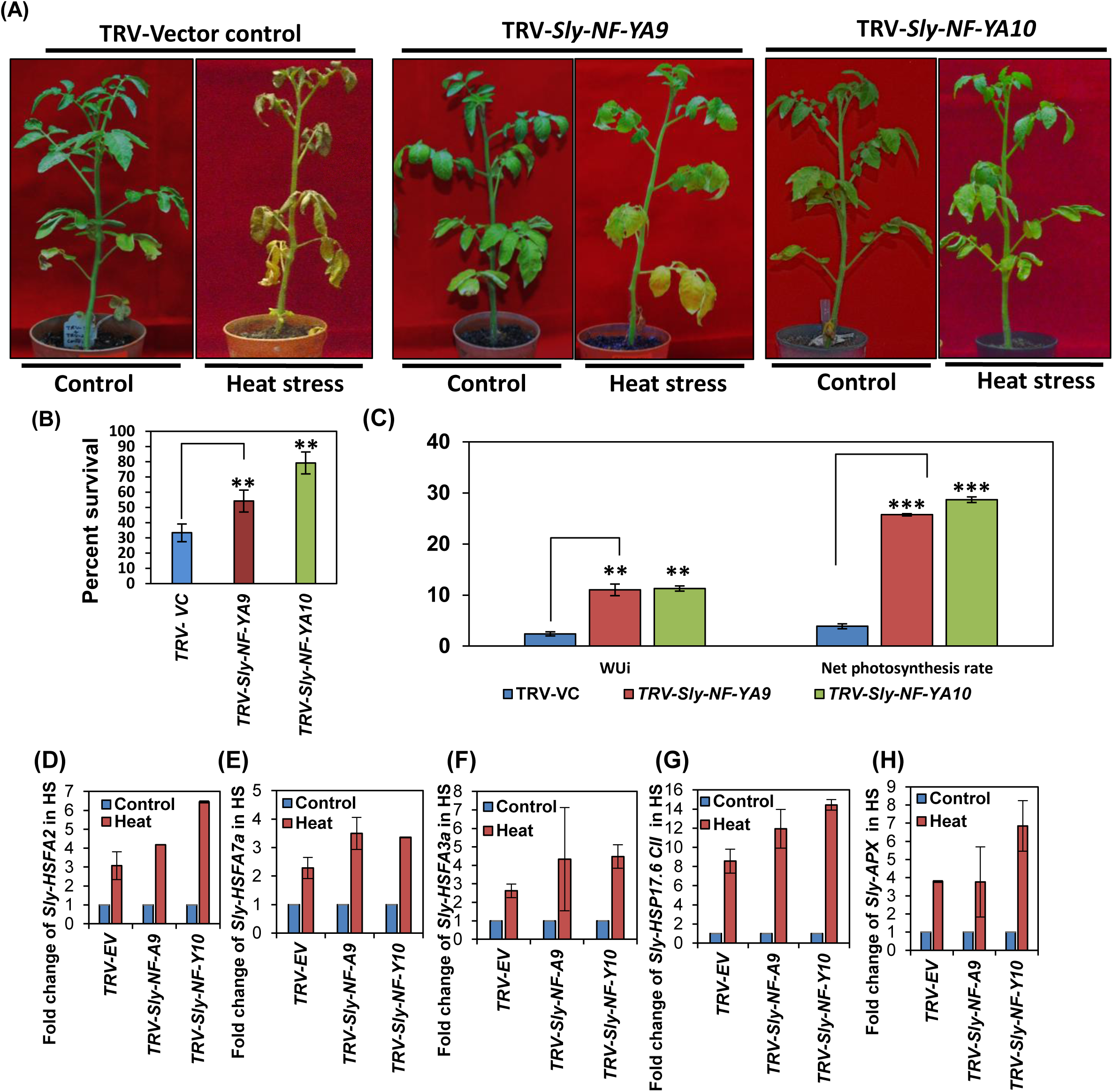
Functional validation of the role of *Sly-NF-A9* and *Sly-NF-A10* in HS response in tomato. (A) Phenotype of TRV-VC (Vector control), *TRV-Sly-NF-YA9* and *TRV-Sly-NF-YA10* silenced plants after heat stress. Fifteen-days-old tomato plants were agro-infiltrated with empty vector (TRV-VC), *TRV-Sly-NF-YA9* or *TRV-Sly-NF-YA10* silencing constructs. Plants were given HS 3-weeks-after infiltration. Phenotype of HS-treated silenced plants was scored six-days-post HS. Experiments were repeated 3-4 times with 8 plants per replicate. (B) Percent survival of TRV-EV, *TRV-Sly-NF-YA9* and *TRV-Sly-NFYA10* silenced plants afters HS. (C) Estimation of water use efficiency (mmol mol^−1^), net photosynthesis rate (μmol m^−2^s^−1^), and transpiration rate of TRV-EV, *TRV-Sly-NF-YA9* and *TRV-Sly-NFYA10* silenced plants afters HS. (D-E) Expression profiles of HSR genes in control and heat treated *TRV-EV, TRV-Sly-NF-YA9* and *TRV-Sly-NF-YA10* silenced plants. The expression levels of genes were calculated using the 2−ΔΔCt method and presented using fold-change values. The fold change normalization was done by setting the control value as one in (D-H). *Actin* was used as endogenous control. Error bars represent the standard deviation of two independent biological replicates. *p<0.05, **p<0.01 and ***p<0.001 vs. wild type, by two-tailed Student’s t-test.

To further delineate the molecular events driving the improved thermotolerance of *TRV-Sly-NF-YA9* and *TRV-Sly-NF-YA10* knock-down plants, we checked whether the expression of conserved HSR genes like *HSFs (Sly-HSFA3a* and *Sly-HSFA7a), Sly-sHSP* and *Sly-APX (ascorbate peroxidase)* is significantly modulated upon heat stress. First, to ascertain that these genes are reliable HSR genes in the VIGS conditions, we show that they all are upregulated in TRV-EV plants under heat stress in comparison to non-stressed control TRV-EV plants (Figure 4D-H, Supplementary figure S10A-E). Furthermore, these HSR genes are inducible in *TRV-Sly-NF-YA9* and *TRV-Sly-NF-YA10* knock-down plants during heat stress as compared to non-stressed conditions (Figure 4D-H, Supplementary figure S10A-E). The strong upregulation of HSR genes in heat stressed *TRV-Sly-NF-YA9* and *TRV-Sly-NF-YA10* knock-down plants in comparison to TRV-EV HS plants confirms the direct involvement of these HSR genes in regulating the thermotolerance of tomato plants (Supplementary figure S10F). Further, since all these HSR genes (except *Sly-APX*) exhibit upregulation in *TRV-Sly-NF-YA9* and *TRV-Sly-NF-YA10* knock-down plants in comparison to TRV-EV plants in non-stressed conditions (Supplementary figure S10G), this indicates that these HSR genes (except *Sly-APX*) are downstream targets of *Sly-NF-YA9* and *Sly-NF-YA10.* Furthermore, promoter analysis (1kb) of these HSR genes revealed the presence of CCAAT boxes in *Sly-HSFA3a* and *Sly-HSFA7a* (Supplementary table S5), suggesting a direct Sly-NF-YA9/10 mediated regulation on these genes and an indirect regulation on other HSR genes. Since the HSR genes are upregulated in *TRV-Sly-NF-YA9* and *TRV-Sly-NF-YA10* knock-down plants, these results suggest that *Sly-NF-YA9* and *Sly-NF-YA10* act as negative regulators of thermotolerance.

### *MIR169s* are regulated by HSFs during heat stress in Arabidopsis

To assess whether the HSF:miR169:NF-YA module under HS is active in other plants too, we decided to evaluate this module in the model plant Arabidopsis. Guan et al. (2013) have reported heat-mediated induction of miR169 and miR398 by Northern hybridization in Arabidopsis. However, the regulation of miR169 in the heat responses of Arabidopsis remains unknown. Using taqman based qRT-PCR analysis of WT control and heat stress challenged Arabidopsis plants, we show that all four mature miR169 forms are strongly upregulated (≥5 fold) under HS (Figure 5A, Supplementary figure S11). Further, to support a probable mechanism of their heat inducibility, promoters of all 14 *At-MIR169* genes (Li et al. 2010) were screened for the presence of HSEs and we found two to multiple HSEs in all *MIR169* promoters (Supplementary table S6). To delineate this speculated interplay among Arabidopsis HSFs and *At-MIR169* promoters, Y1H assays were performed (Supplementary table S7) with 21 At-HSFs and 11 *At-MIR169* promoters (as six precursors are transcribed as three bi-cistronic units viz., Pro-*At-MIR169i/j*, Pro-*At-MIR169k/l*, Pro-*At-MIR169m/n*; Li et al. 2010). This analysis identified 5 HSFs interacting with 3 *At-MIR169* promoters (Figure 5B). Three HSFs (At-HSFB3a, At-HSFA7b, At-HSFB2b) regulate *At-MIR169b*; At-HSFA2a controls *At-MIR169d* transcription and transcription of *At-MIR169h* is regulated by At-HSFA1d. At*-*HSFA2 is considered as the key regulator of Arabidopsis HS-response and is required for the maintenance of acquired thermotolerance (Charng et al. 2007). Therefore, we decided to dissect the At-HSFA2 mediated *At-MIR169d* module for characterization of HS-response and thermotolerance in Arabidopsis. In wild type Arabidopsis plants, we find strong induction of *At-MIR169d* precursor (≥15 fold) upon HS (Figure 5C, Supplementary figure S12). Transgenic promoter-*At-MIR169d:GUS* plants support this heat-dependent transcriptional acceleration (Figure 5D, E). Moreover, *GUS* aided reporter analysis of promoter-*At-MIR169d:GUS* in WT and in the *At-hsfa2* knockout mutant (Supplementary figure S13A, B, C) background in HS highlights minimal *GUS* expression in the *At-hsfa2* mutant as compared to WT plants (Figure 5F, G). Further, there is strong reduction of *At-MIR169d* precursor levels in *At-hsfa2* mutant as compared to WT plants (Figure 5G) confirming At-HSFA2-mediated regulation of *At-MIR169d* in Arabidopsis HS and suggesting a general regulatory loop in plants.

**Figure 5.**
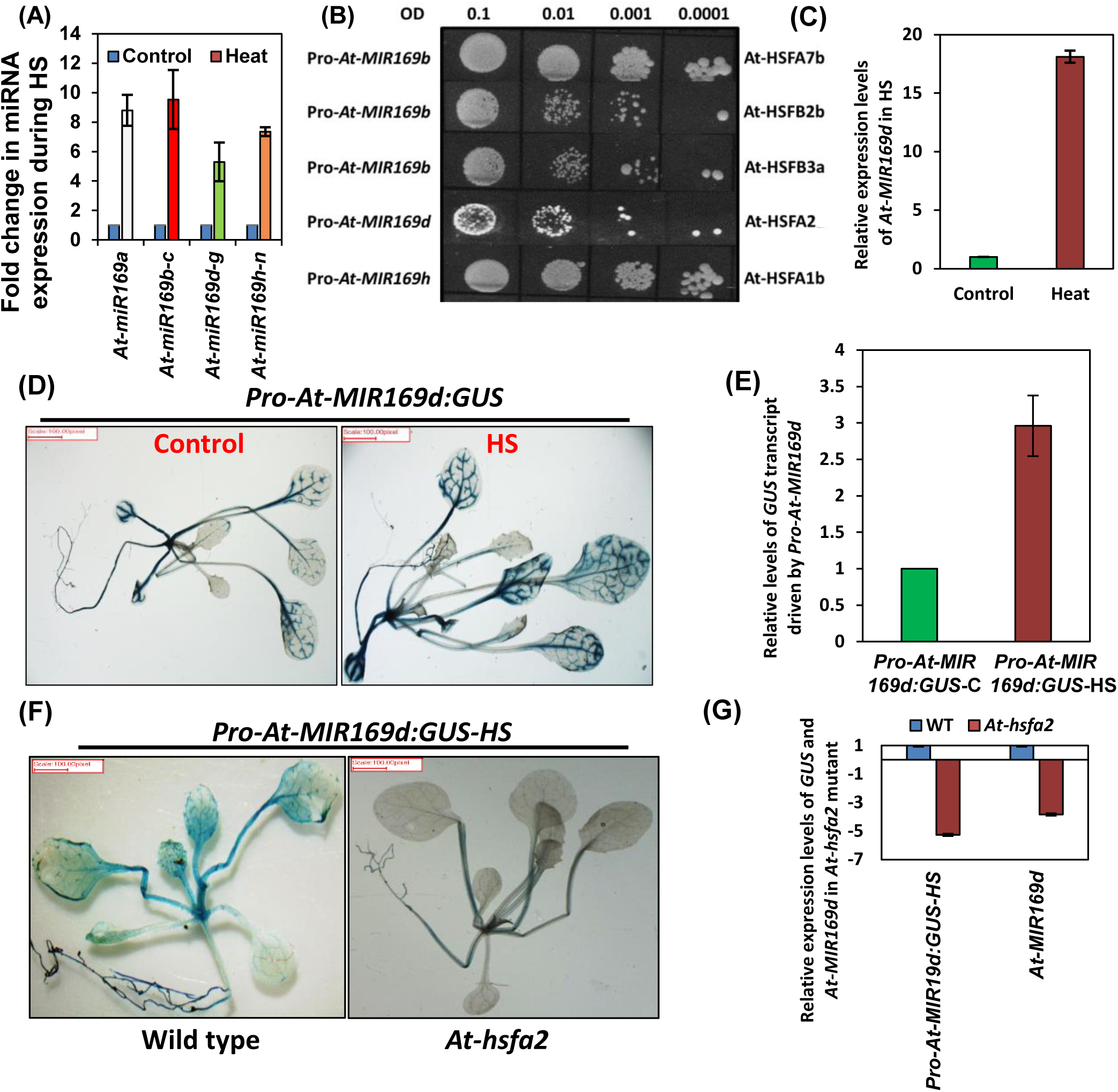
Transcriptional regulation of *MIR169* promoters by HSFs in Arabidopsis. (A) miR169s expression in response to heat stress as determined by Taqman-based qRT-PCR of control and heat challenged plants. 5S rRNA was used as the normalization control. Experiment was repeated three times and average values are plotted as bars, error bars depict standard deviation between the replicates. (B) Y1H assay showing the positive interaction of HSFs with the promoters of different *At-MIR169s*, Y1H assay shows binding of At-HSFA7b, At-HSFB2b, At-HSFB3a, At-HSFA2 and At-HSFA1b on *At-MIR169b*, d and h promoters. This experiment was repeated at least three times with similar results, and data from one representative experiment is shown. Serial dilutions (O.D. 0.1 to 0.0001) of yeast cultures were spotted onto plates lacking URA and LEU with specific Aureobasidin A concentration and incubated for 3 days. The Aureobasidin A concentrations for all *At-MIR169* promoters is presented in supplementary table S3. (C) Expression levels of *At-MIR169d* in wild-type plants subjected to 0 (control) and 2 h heat stress at 42 °C. Data are shown as means ± SD of three biological replicates and normalized to actin reference gene using (2−ΔΔCT) method. (D) Expression patterns of *Pro-At-MIR169d:GUS* lines of transgenic Arabidopsis plants subjected to 0 (control) or 2 h of heat stress at 42°C. (E) Levels of GUS transcripts in *Pro-At-MIR169d:GUS* lines, measured by qRT-PCR after 0 or 2 h of heat stress at 42°C. (F) Transcriptional activity of *Pro-At-MIR169d:GUS* reporter construct in WT and *At-hsfa2* mutant background during 2 h of heat stress at 42°C. (G) Expression of *GUS* transcript driven by *Pro-At-MIR169d* and *At-MIR169d* precursor in soil-grown wild-type plants of Arabidopsis and *At-hsfa2* mutant. To represent the negative fold-change on y axis in figure (G), the relative expression values of down-regulated genes were transformed using the formula [-(1 divided by RQ ≤0.5)]. The fold change normalization was done by setting the control value as one in Figure (A, C, E and G). Error bars represent the standard deviation. These experiments were repeated at least three times with similar results, and data from one representative experiment are shown in D and F.

### The miR169d:NF-YA2 regulatory node is heat responsive and spatially co-expressed in Arabidopsis

To explore the functional significance of heat-mediated regulation of *MIR169s*, we assessed the expression of the eight canonical *NF-YA* targets (Sorin et al. 2014) in HS in Arabidopsis. qRT-PCR based profiling shows significant reduction of *At-NF-YA2* (−3.5 fold), *At-NF-YA3* (−2.2 fold), *At-NF-YA6* (−2.1 fold) and *At-NF-YA10* (−2.1 fold) transcripts in WT plants when exposed to HS (Supplementary figure S14A, B). Further, to assign the functional relevance of the selected *At-MIR169d* isoform in particular, we analyzed target mimicry plants generated by overexpressing MIM169defg constructs under CaMV35S, using constructs from Todesco et al. (2010). The expression analysis of mature form At-miR169defg in these MIM169defg lines revealed the significant reduction of this form (Supplementary figure S15A). The target mimic lines sequester the specific mature miRNAs by acting as non-cleavable targets; thereby they protect the target genes from miRNA cleavage (Todesco et al. 2010). The expression of *At-NF-YA2* is significantly increased in *MIM169defg* lines as compared to WT plants (Supplementary figure S15B). Moreover, overexpressing *At-MIR169d* (At-*MIR169d*-OE) precursor under constitutive CaMV35S promoter in Arabidopsis strongly represses (−4 to −11-fold in three different *At-MIR169d-OE* lines) *At-NF-YA2* transcripts, establishing it as the major miR169d target (Supplementary figure S15C). Moreover, promoter-*At-NF-YA2* lines do not show HS-mediated up-regulation specifying a pure post-transcriptional regulation (Supplementary figure S16A, B).

To delineate the existence and localization of miR169d:*At-NF-YA2* functional module in plant tissues and development stages, promoter reporter (*GUS)* lines were generated by transforming WT Arabidopsis plants with constructs that contained the 2 kb genomic sequences upstream of the *At-MIR169d* precursor. Parallelly, 2 kb region ahead of the translation start site of *At-NF-YA2* was similarly assessed. The *At-MIR169d* and *At-NF-YA2* promoters show prominent GUS localization in the vasculature of roots and aerial parts of seedling, mature rosette/cauline leaves and sepals in flowers of Arabidopsis, suggesting functional co-existence of the miR169d:*NF-YA2* module (Supplementary figure S17).

### Arabidopsis plants with enhanced miR169d or reduced *NF-YA2* expression are thermotolerant

To further investigate the miR169d:*NF-YA2* module in HS-response in Arabidopsis, we assayed 2-weeks-old plants of *At-MIR169d*-OE, *At-NF-YA2*-OE (*NF-YA2* overexpressed under 35SCaMV promoter, Supplementary figure S18), *At-nf-ya2* knockout mutant (Supplementary figure S19), an miR169d resistant version of *At-NF-YA2* under native promoter (*pNF-YA2:NF-YA2*-r) and MIM169defg lines for their ability to survive HS. The survival rate of plants was gauged 6-days-after exposure to 2 h of HS at 42 ᵒC. Plants overexpressing *At-MIR169d* (*At-MIR169d*-OE) that have reduced target *At-NF-YA2* levels and plants that lack *At-NF-YA2* transcripts (*At-nf-ya2* mutant) exhibit twice the survival rate than WT plants, in contrast, plants with high abundance of *At-NF-YA2* transcripts viz. *At-NF-YA2*-OE, *MIM169defg* and the *pNF-YA2:NF-YA2*-r (a variant *At-NF-YA2* transcript that allows miRNA169 binding but escapes miRNA-mediated cleavage) are nearly 3-folds more HS sensitive (Figure 6A, B). While only ∼10% plants survive in transgenic plants overexpressing *At-NF-YA2* mRNAs, as much as 80-90% plants survive the HS in *At-MIR169d-OE* and *At-nf-ya2* mutant plants. This establishes that HS-tolerance in Arabidopsis by At-miR169d is mediated by down-regulation of *At-NF-YA2* functions wherein, *At-NF-YA2* acts as negative regulator of heat stress-response.

**Figure 6:**
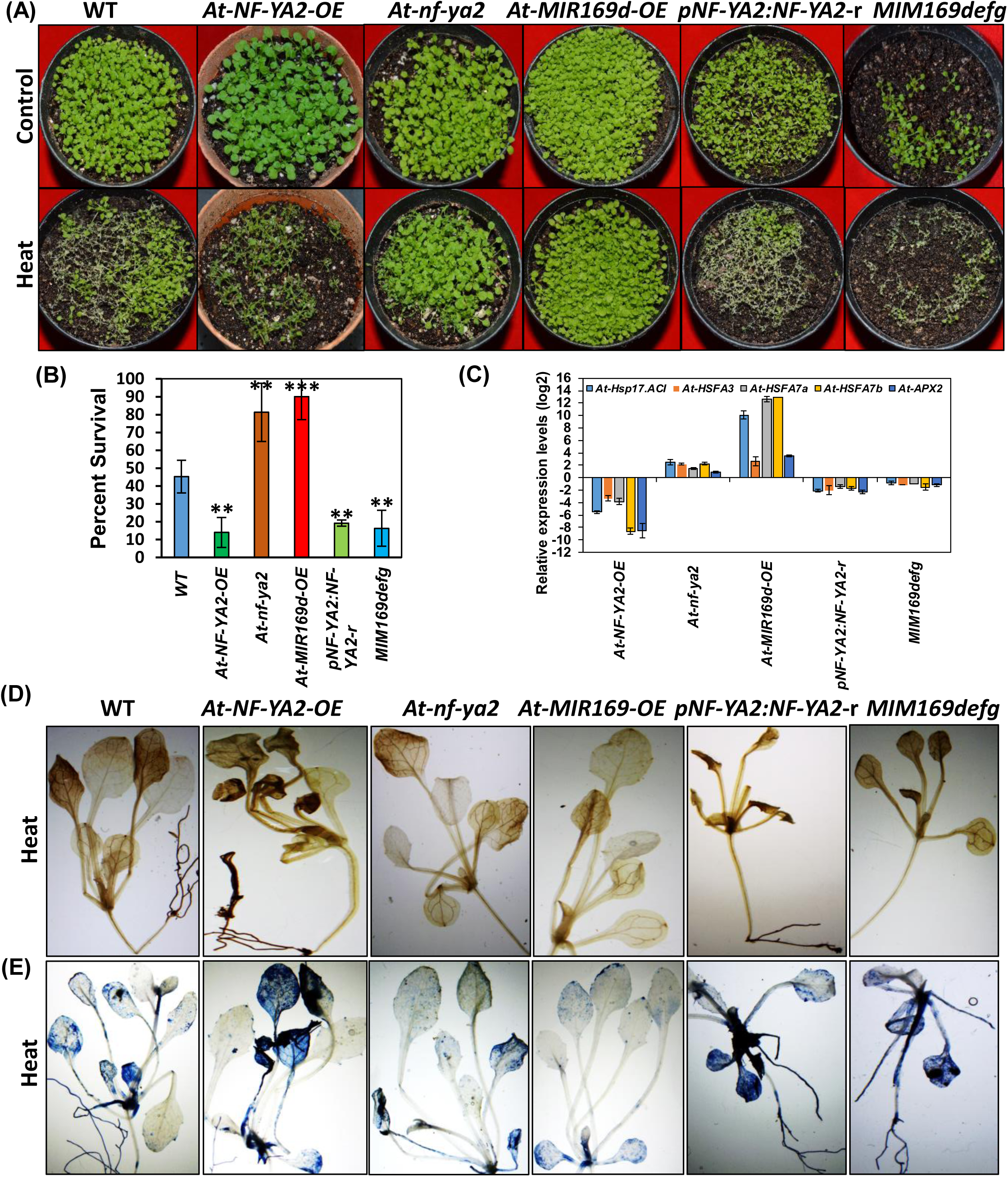
Assaying the miR169d:At-NF-YA2 node in heat stress in Arabidopsis. (A) Thermotolerance was gauged in *At-MIR169d-OE*, *At-NF-YA2-OE*, *At-nf-ya2, pNF-YA2:NF-YA2*-r and MIM169defg transgenic plants in comparison to wild-type (WT) plants. Two-weeks-old soil-grown plants were subjected to 0 (control) or 2h heat stress at 42°C, and damage was recorded 6 days later. (B) Estimation of survival (percentage) of HS treated *At-MIR169d-OE*, *At-NF-YA2-OE*, *At-nf-ya2, pNF-YA2:NF-YA2*-r and MIM169defg plants. Plants were assayed for heat stress tolerance after 1 week of HS. These experiments were repeated at least five times with similar results, and average data of all experiments are shown. Error bars, ± SD of five repeats. (C) Expression patterns of heat stress-responsive genes in WT, *At-MIR169d-OE*, *At-NF-YA2-OE*, *At-nf-ya2, pNF-YA2:NF-YA2*-r and MIM169defg plants. Data are shown as means ± SD of biological replicates and normalized to *Actin* using (2^−ΔΔCT^) method. Log2 transformation was applied to the RQ data to obtained negative fold change. (D-E) DAB and Trypan blue staining of HS treated *At-MIR169d-OE*, *At-NF-YA2-OE*, *At-nf-ya2*, *pNF-YA2:NF-YA2*-r and MIM169defg plants to estimate the production of ROS and cell death, respectively. *p<0.05, **p<0.01 and ***p<0.001 vs. wild type, by two-tailed Student’s t-test. Three independent lines of *MIR169d-OE* were used and similar results were obtained for all studied parameters, a single representative picture was shown in the figure A, B, D and E. The fold change normalization was done by setting the control value as one for figure (C).

To further explore the mechanism underlying the better survival of *At-MIR169d*-OE and *At-nf-ya2* lines as well as the sensitivity of *At-NF-YA2*-OE, MIM169defg and *pNF-YA2*:*NF-YA2*-r lines, we analyzed the expression of several heat responsive genes reported in literature (Larkindale and Vierling 2008) in all these lines. Expression profiles of 11 HSR genes in response to HS (Supplementary table S8) revealed that the improved thermotolerance in *At-MIR169d*-OE and *At-nf-ya2* mutant plants is correlated with enhanced expression of *At-Ascorbate peroxidase2* (*At-APX2*), *At-HSFA3a*, *At-HSFA7a*, *At-HSFA7b* and *At-HSP17.ACI* genes (Figure 6C, Supplementary figure S20). In contrast, transgenic plants *At-NF-YA2*-OE, *pNF-YA2:NF-YA2*-r, MIM169defg that are defective in heat-responsive gene regulation, exhibit reduced *At-APX2, At-HSFA3a, At-HSFA7a, At-HSFA7b, and At-HSP17.ACI* transcripts (Figure 6C, Supplementary figure S20). These results ascertain At-NF-YA2 as negative regulator of many HSR genes and thermotolerance in Arabidopsis.

It is well established that imposition of HS can give rise to excess levels of reactive oxygen species (ROS), resulting in cellular oxidative damage. *APX* plays crucial role as H_2_O_2_-scavenging enzyme in plant cells. Therefore, a higher level of *At-APX2* transcripts in *At-MIR169d*-OE and *At-nf-ya2* mutant plants should effectively maintain the antioxidants that protect plants. Indeed, these plants have low H_2_O_2_ levels as judged by DAB staining (Figure 6D). In contrast, the overexpression of *At-NF-YA2* gene (*At-NF-YA2-OE*) and chelation of At-miR169d (MIM169defg lines) and *pNF-YA2:NF-YA2*-r show enhanced DAB staining due to higher H_2_O_2_ levels (Figure 6D). Further, cellular death estimation as a consequence of this in *At-MIR169d*-OE and *At-nf-ya2* lines is negligible while it is much pronounced in the *At-NF-YA2*-OE, *MIM169defg* and *pNF-YA2:NF-YA2*-r lines, as assessed by trypan blue staining (Figure 6E).

### Arabidopsis At-NF-YA2 negatively regulates *HSR* genes leading to HS sensitivity

Repressed expression of *HSR* genes in *At-NF-YA2* abundant lines and enhanced expression of *HSR* genes in *At-NF-YA2* down-regulated lines confirms that *At-NF-YA2* is a negative regulator of these HSR genes (Figure 6C, Supplementary figure S20). *At-HSP17.ACI*, *At-HSFA3a*, *At-HSFA7a, At-HSFA7b* and *At-APX2* promoters have NF-YA binding sites (CCAAT-box), making them probable downstream targets of At-NF-YA2 transcription factor (Figure 7A). Indeed, ChIP-qPCR and Y1H assay shows that the At-NF-YA2 protein interacts with *At-HSFA3a* and *At-HSFA7b* promoters, but not with *At-HSP17.ACI*, *At-HSFA7a* and *At-APX2* promoter (Figure 7B, Supplementary figure S21).

**Figure 7:**
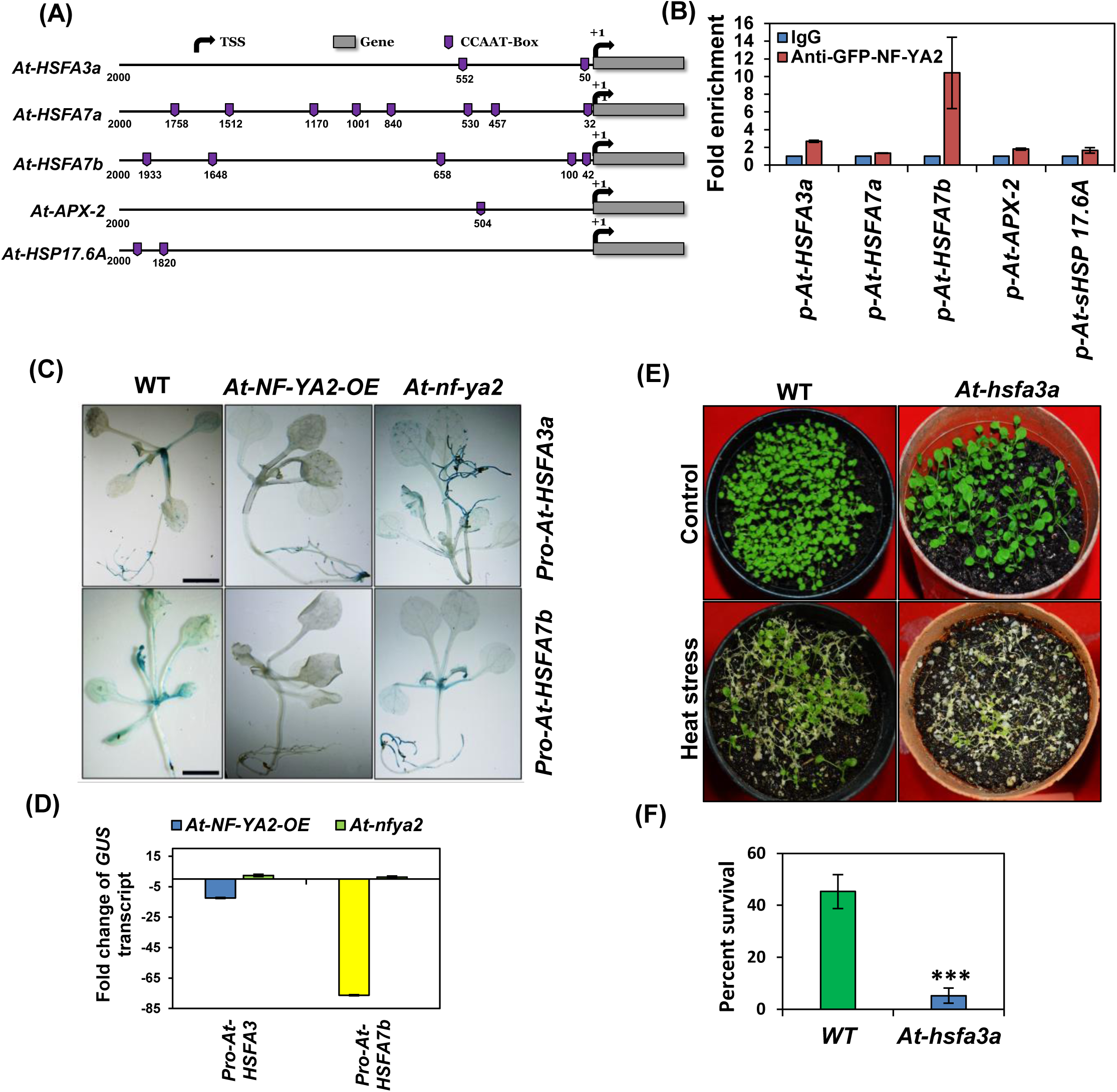
Transcriptional regulation of *At-HSFA3a* and *At-HSFA7b* by At-NF-YA2 during heat stress in Arabidopsis. (A) Graphical representation of CCAAT cis-elements in At-HSR gene promoters HSFA3a, HSFA7a, HSFA7b, APX and sHSP17.6A. The 2 kb upstream sequences of start codon were analysed by plant care and Plant PAN promoter analysis tools. The positions of CCAAT boxes are depicted by violet coloured tabs. (B) Chromatin immunoprecipitation (ChIP)-qPCR assay of At-NF-YA2 on At-HSR gene promoters. (C-D) *In-planta* estimation of transcriptional regulation of *At-HSFA3a* and *At-HSFA7b* transcription by At-NF-YA2 during heat stress. (C) *Pro-At-HSFA3a:GUS* reporter construct was agro-infiltrated in WT, *At-NF-YA2-OE* and *At-nf-ya2* backgrounds. *GUS* was visualized 2-days post-infiltration by histochemical staining. (D). qRT-PCR based quantification of *GUS* transcripts in promoter reporter constructs infiltrated plants, 2-days-post infiltration by using *GUS* specific primers. The *NPTII* gene present in the same vector backbone was used as reference control. To represent the negative fold-change on y axis in, the relative expression values of down-regulated genes were transformed using the formula [-(1 divided by RQ ≤0.5)]. The fold change normalization was done by setting the control value as one. (E) Thermotolerance assay of WT and *At-hsfa3a* mutant plants. Two-weeks-old soil-grown plants were subjected to 0 (control) or 2h heat stress at 42°C, and damage was recorded 6 days post-stress. (F) Estimation of survival (percentage) of WT and *At-hsfa3a* plants 6 days post heat stress. The experiments were repeated at least three times with similar results, and average data of all experiments are shown. Error bars, ± SD of three repeats. *p<0.05, **p<0.01 and ***p<0.001 vs. wild type, by two-tailed Student’s t-test.

To further confirm the repression of *At-HSFA3a* and *At-HSFA7b* transcription by At-NF*-*YA2 *in planta,* GUS promoter reporter constructs Pro-*At-HSFA3a:GUS* and Pro-*At-HSFA7b:GUS* were introduced in WT, *At-NF-YA2*-OE and *At-nfy-a2* mutant backgrounds. Expression analysis and quantification of *GUS* transcription using these lines confirm At-NF-YA2-mediated repression of *At-HSFA3a* and *At-HSFA7b* as *GUS* accumulation is reduced in *At-NF-YA2*-OE line and strongly expressed in *At-nf-ya2* mutant background as compared to WT (Figure 7C, D). Thus, At-NF-YA2 regulates the transcription of *At-HSFA3a* and *At-HSFA7b* directly. Thermotolerance assay using *At-hsfa3a* and *At-hsfa7a* mutants show high sensitivity to HS (Figure 7E, Supplementary figure S22A, B), only 5% *At-hsfa3a* and 1.7% *At-hsfa7a* plants are able to recover as compared to 45% WT plants six-days-after HS imposition (Figure 7F, Supplementary figure S22B). This confirms the positive role of *At-HSFA3a* and *At-HSFA7a* genes in maintaining thermotolerance. Furthermore, role of *At-HSFA7b* gene in salt and heat stress tolerance via an E-box-like motif was recently reported by Zang et al. (2019). Thus, in Arabidopsis, HS induces At-HSFA2 that increases the expression of At-miR169d which in turn post-transcriptionally down-regulates *At-NF-YA2* transcriptional repressor. At-NF-YA2 in turn enhances transcription of HSR genes like *At-HSFA3a* and *At-HSFA7b*. This regulatory feedback loop plays a critical role in thermotolerance.

## Discussion

Previously we have shown heat-responsiveness of *MIR169* members in tomato (Rao et al. 2020). Moreover, heat-mediated induction of miR169 in Arabidopsis (Guan et al. 2013; Li et al. 2010) and in rice (Zhao et al. 2007; Zhao et al. 2009) is also reported. This study focuses on defining the underlying upstream regulatory mechanism viz., the HSF-mediated transcriptional control of *MIR169* genes as well as the post-transcriptional regulation of downstream targets of miR169s i.e., the *NF-YA* transcription factors, which in turn transcriptionally regulate HSR genes in both tomato and Arabidopsis.

### Cooperative HSFs binding regulate *MIR169* transcription in HS

At-HSFA1b and At-HSFA7b mediated induction of *At-MIR398b* during heat stress has been reported earlier in Arabidopsis (Guan et al. 2013). Presence of multiple HSEs in promoters foster cooperative interactions between multiple HSF trimers (Topol et al. 1985; Xiao et al. 1991; Bonner et al. 1994). The promoters of HS-responsive *MIR169s* exhibit multiple HSFs binding making these promoters suitable targets for cooperative binding of HSFs as evidenced by Y1H/ChIP-qPCR assays in both tomato and Arabidopsis (Figures 1A-I, 4B and 6B). The binding ability of Sly-HSFs on *Sly-MIR169* promoters in tomato was further confirmed by transient assays in *N. benthamiana*, where effector plasmids containing different Sly-HSFs were co-transformed with specific *Sly-MIR169* promoters hooked with *GUS* reporter gene. Sly-HSF-mediated activation of *Sly-MIR169a-1*, *Sly-MIR169b*, *Sly-MIR169c*, *Sly-MIR169d* and *Sly-MIR169d-1* promoters was confirmed for all Sly-HSFs except Sly-HSFA1a. Individual co-transfection of Sly-HSFA1a with Pro-*Sly-MIR169b* was unable to induce *GUS* transcription. Co-infiltration of Sly-HSFA1a with Sly-HSFA7a (the other HSF binding on Pro-*Sly-MIR169b*) leads to strong induction of *GUS* transcripts, suggesting the cooperative interactions between Sly-HSFA1a and Sly-HSFA7a. The *Sly-HSFA1a*-VIGS study suggests that threshold level Sly-HSFA1a is sufficient to regulate transcription of its target genes, as portrayed by strong reduction of *Sly-MIR169b* transcription (Figure 2D). Some studies have also reported the stress-mediated hetero-oligomeric complex formation of HSFA1a with other A-class HSFs to synergistically regulate the expression of a number of small HSPs, DnaJ/Hsp40, and HSP70 (Chan-Schaminet et al. 2009; Liu et al. 2011; Hahn et al. 2011; Li et al. 2014).

The HSF ‘B-class’ is known primarily as a repressor of transcription (Czarnecka-verner et al. 2000; Ikeda et al. 2009). We find Sly-HSFB1a-mediated transactivation of Pro-*Sly-MIR169d* (Supplementary figure S3D). Sly-HSFB1a has been shown to activate transcription of Myc-HSP17.6, alone and synergistically with Sly-HSFA1a as well as with acidic activators like HAC1/CBP (Bharti et al. 2004). The class-A1 HSFs are considered as master regulators of HS response (Liu et al. 2011; Yoshida et al. 2011). In our study we find ‘Class-A’ HSFs regulate expression of all *MIR169* either singularly or in combination with other HSFs belonging to class-A/-B/-C in both tomato and Arabidopsis. The ‘Class-A1’ HSFs regulate 2 of 5 and 1 of 3 *MIR169* promoters in tomato and Arabidopsis, respectively reiterating the significance of this class in HS tolerance.

### Alleviation of *Sly-NF-YA9*/*Sly-NF-YA10* and *At-NF-YA2* enhances thermotolerance

Further, analysis of thermotolerance of plants with reduced *NF-YA* levels (*Sly-MIR169-OE*,TRV-*Sly-NF-YA9* and TRV-*Sly-NF-YA10* in tomato and *At-MIR169d*-OE and *At-nf-ya2* knockout mutant in Arabidopsis) and those with higher *NF-YA* expression (At-*NF-YA2*-OX, p*NF-YA2*:*NF-YA2*-r and *MIM169defg*) revealed that reduced *NF-YA* levels increase thermotolerance (higher survival rate) and there is elevated expression of HSR genes including *HSF* genes, APX and several HSP genes (Figure 4A-F, 4D-H, Supplementary figure S7, S10 and Figure 7C, Supplementary figure S20). Thus, establishing *Sly-NF-YA9 and Sly-NF-Y10* as negative regulators of HSR gene expression and thermotolerance in tomato and *At-NF-YA2* in Arabidopsis. Other reports by Ceribelli et al. (2008) and Leyva-González et al. (2012) have also shown NF-YAs as repressors. Further, we have demonstrated in both tomato and Arabidopsis that the diminished expression of *Sly-NF-YA10* (Rao et al. 2020) and *At-NF-YA2* (Figure 2C, Supplementary figure S14A, B and supplementary figure S16A, B) upon HS is a post-transcriptional regulation which is a result of miR169-mediated cleavage and not a transcriptional regulation.

### At-NF-YA2 binds to HSR gene promoters

Our study establishes direct At-NF-YA2 mediated transcriptional regulation of *At-HSFA3a* and *At-HSFA7b* by Y1H/ChIP-qPCR assays. Further, loss of Pro-*HSFA3a*:*GUS*, Pro-*HSFA7a*:*GUS* and Pro-*HSFA7b:GUS* activity in *At-NF-YA2*-OE lines and high *GUS* expression in *At-nf-ya2* mutant background confirms regulation of these promoters by At-NF-YA2 as a transcriptional repressor (Figure 7C and D). Studies in literature support the involvement of *HSFA3a* in governing thermotolerance in Arabidopsis and tomato (Li et al. 2013; Wu et al. 2018). Larkindale and Vierling (2008) have reported *At-HSFA3a* as a universal candidate gene commonly induced in heat acclimated plants in different regimes of heat stress treatment. We also observe high thermo-sensitivity in *At-hsfa3a* mutant plants (Figure 7E). It is known that the NF-YA subunit forms functional trimer with NF-YB and NF-YC subunits to regulate transcription of a large number of genes (Warpeha et al. 2007; Kumimoto et al. 2010; Sato et al. 2014). Sato et al. (2014) have shown synergistic interaction of At-DREB2A with trimer of NF-YA2, NF-YB3 and NF-YC10 to form a transcriptional complex that activates *At-HSFA3* expression in Arabidopsis protoplasts. *DREB2A* is also a HS inducible gene and its overexpression significantly increases HS tolerance (Sakuma et al. 2006). Moreover, Sato et al. (2014) have found heat-inducible expression of *NF-YB3* and *NF-YC10* but relatively stable expression of *At-NF-YA2* transcripts upon HS. Here, we conclusively show that there is heat-mediated down-regulation of *At-NF-YA2* (Supplementary figure S14A, B) that is a consequence of *miR169*-mediated cleavage and also that At-NF-YA2 acts as a repressor of *At-HSFA3a* transcription (Figures 5C, 6C and 6D). Thus, we speculate that the reduced availability of NF-YA2 subunits during HS (due to degradation by miR169d) may favor functional DREB2A:NF-YC-10:NF-YB3 complex formation, which in turn up-regulates *At-HSFA3a* expression during HS (Figure 8).

**Figure 8:**
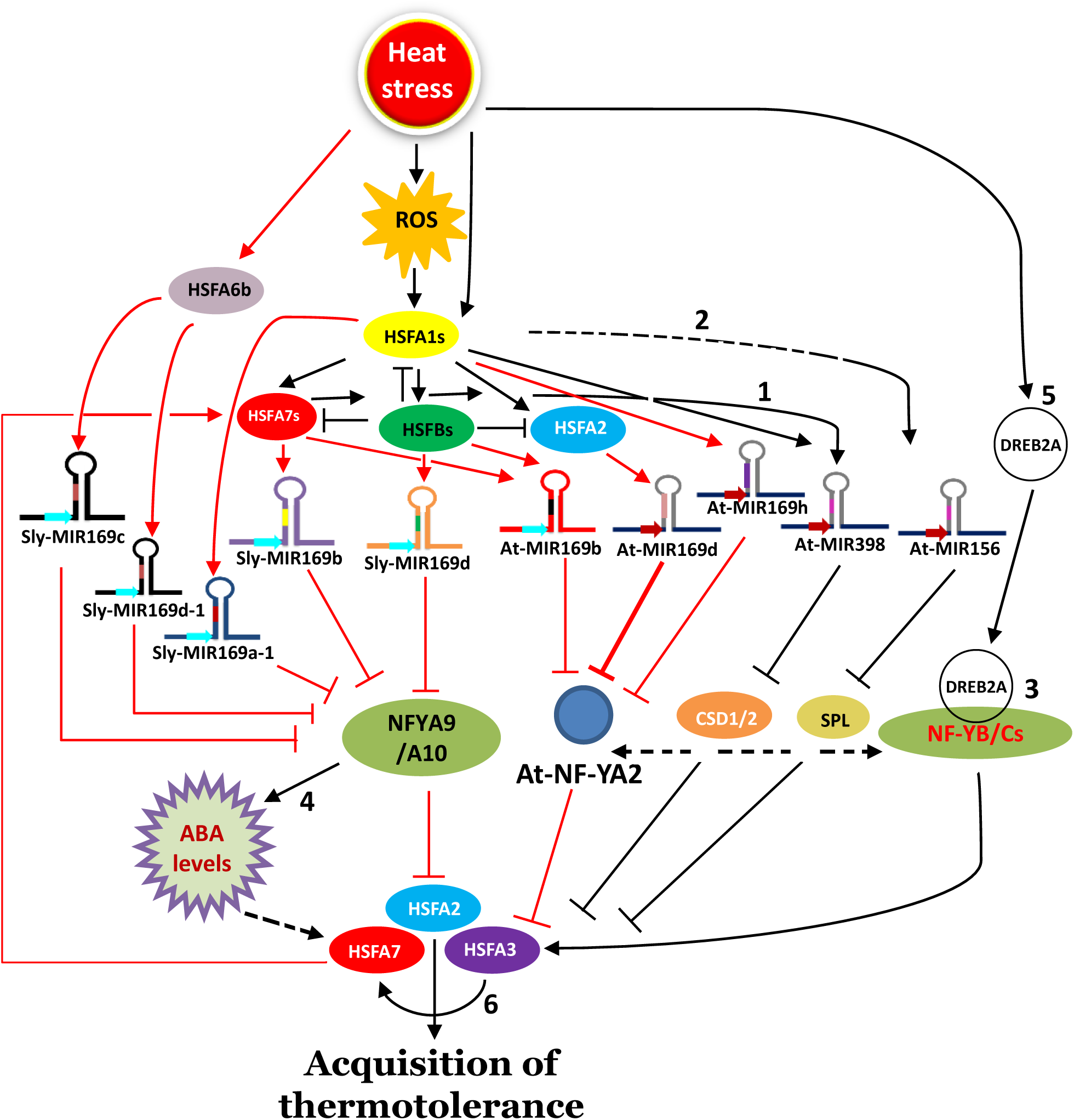
Model for HSF-induced miR169:NF-YA module in mediating thermotolerance in tomato and Arabidopsis. Heat stress induces the expression of HSFs by altering the cellular ROS levels as well as other cues independent of ROS signaling. HSFA1a, HSFA2 and HSFA7a are the key player of plant HSR response. These HSFs bind to the promoters of *MIR169* leading to transcriptional enhancement of miR169s in both tomato and Arabidopsis. Enhanced accumulation of miR169s reduces the levels of *Sly-NF-YA9/A10* in tomato and *At-NF-YA2* in Arabidopsis that leads to enhancement of HSR genes expression like *HSFA2, HSFA3* and *HSFA7s*, possibly via altering ABA levels. In Arabidopsis, At-miR169-mediated down-regulation of *At-NF-YA2* during HS may favor functional DREB2A:NF-YC-10:NF-YB3 complex formation to up-regulate *At-HSFA3a* expression during HS. Enhanced levels of HSFAs in turn feed the transcription of *MIR169* genes. The heat-mediated orchestration of parallel miRNA pathways (miR169:NF-YA, miR398:CSD and miR156:SPL) alters the levels of HSR genes and regulate plant thermotolerance. Solid lines denote a confirmed interaction and dotted lines are predictive. Red and black lines denote findings of this study and existing literature, respectively. Black colored numerals refer to the studies that established the particular connection between different components of the model. Reference key-1: Guan et al. 2013; 2: Stief et al. 2014; 3: Sato et al. 2014; 4: Zhao et al. 2009, Luan et al. 2015, Ding et al. 2016; 5: Sakuma et al. 2006. CSD: Copper superoxide dismutase; SPL: SQUAMOSA-promoter binding like; ABA: Abscisic acid; ROS: Reactive oxygen species; DREB: Dehydration responsive element binding protein.

### An HSF-mediated feedback loop regulates *MIR169* in heat stress

Furthermore, under HS, *Sly-HSFA7a is* up-regulated in *Sly-NF-YA9/A10* silenced tomato plants and there is enhanced expression of *At-HSFA7b* in *At-nf-ya2* mutant plants (Figures 4D-H, 6C, and supplementary figure S20 and table S8). We have shown that these HSFs interact with *MIR169* promoters. Moreover, the enhanced expression of *Sly-HSFA3* in *Sly-NF-YA9/A10* silenced tomato plants and *At-nf-ya2* mutant plants (Figure 4F and 6C), further adds another level of loop mediated regulation as Sly-HSFA3 was known to regulate *Sly-HSFA7* transcription (Yoshida et al. 2008), which is established as upstream regulator of *Sly-MIR169b* in tomato and Arabidopsis in our analysis (Figure 1A, D, 5B and 8). This suggests the existence of an HSF-mediated feedback module regulating HS-mediated transcriptional regulation of *MIR169s* directly that may be crucial for thermotolerance in both tomato and Arabidopsis (Figures 1A, 4D-H, 4B, 6C and 8). While our study demonstrates a direct role of NF-YA class to regulate HSFs, one cannot rule out the possibility that NF-YAs regulate HSFs indirectly. It is known that *NF-YA* genes regulate abiotic/biotic stress response mainly via ABA pathway (Zhao et al. 2009; Luan et al. 2015; Ding et al. 2016). Similarly, HSFs are also induced in response to ABA in Arabidopsis (Huang et al. 2016). Thus, alteration in HSFs levels in *Sly-NF-YA9/10* down-regulated tomato plants or *At-nf-ya2* knockdown Arabidopsis plants could also be a result of the fluctuations in the ABA levels of the silenced plants. Moreover, the HSF perturbation during HS could also be a result of ROS-mediated HSF induction (Volkov et al. 2006; Guan et al. 2013), thus feeding to the HSF:*MIR169*:*NF-YA* loop. We find that heat stress induces At-HSFA2a that in turn transcriptionally activates *At-MIR169d*. At-HSFA2a has also been shown to be positively regulated by NF-YC10 together with HSFA1s (Yoshida et al. 2011; Sato et al. 2014). Overexpression of miR156 and the resultant down-regulation of target SPL genes also induce HSR genes including *At-HSFA2a* (Stief et al. 2014) (Figure 8). Moreover, our data shows other heat responsive miR169s may also act parallely to regulate NF-YAs and contribute towards heat tolerance of Arabidopsis (Figure 8). Thus, multiple pathways work in tandem, interacting among themselves to regulate cellular homeostasis during HS.

### A conserved thermotolerance mechanism in tomato and Arabidopsis

It is also noteworthy, that tomato Sly-NF-YA9/A10 are the best orthologues of At-NF-YA2 (Supplementary figure S23). Also, we show that Sly-HSFA7a and At-HSFA7b regulate the transcription of *MIR169* in both tomato and Arabidopsis, respectively. Interestingly, the heat-responsive *At-MIR398* is also shown to be regulated by an HSFA7 class member (Guan et al. 2013). This suggests that the Sly-NF-YA9/10 and At-NF-YA2 orthologues and Sly-HSFA7-class members have conserved roles in heat tolerance in both plant species.

Our study also highlights the existence of HSF-independent pathways in regulating *MIR169s* transcription during HSR as HSF-mediated regulation could be established for only 5 out of 15 and 3 out of 11 *MIR169* promoters of tomato and Arabidopsis, respectively (Figures 1A, 4B). Though these *MIR169* loci may be indirectly controlled by other cascades regulated by HSFs like the DREB2-mediated *HSFA3* regulation or the NF-YA-mediated regulation of HSFs as we have shown. Moreover, in tomato miR169c is the most abundant miR169 in control as well as heat stress conditions (Supplementary table S3b), which is encoded by a single locus and governed by HSFA6b (Figure 1A, F). The present study establishes a conserved strategy that involves HSF-mediated induction of *miR169* in both tomato and Arabidopsis that leads to reduction of target *NF-YA* genes that in-turn regulate HSF transcription which feeds back to the miR169:*NF-YA* pathway. Down-regulation of *At-NF-YA2* homologues may be a viable strategy for improving the thermotolerance and yield stability of crop plants.

## Materials and Methods

### Plant materials and growth conditions

Tomato cultivar CLN seeds (Balyan et al. 2020) were germinated on filter paper soaked with deionized water at 26 °C. Three-days-old seedlings of uniform growth were transplanted in plastic pots, filled with soilrite and placed in a plant growth chamber (CMP6050, Conviron, Canada) maintained at 26 °C/21 °C (day/night: 16/8 h) relative humidity 60%, light intensity 300 µM per m^2^ per sec.

*Arabidopsis thaliana* (ecotype Columbia) was used as the wild-type in this study. Seeds of the *MIR169d*-OE, *NF-YA2*-OE, Pro-*NF-YA2:NF-YA2*-r, Pro-*At-MIR169d: GUS*, Pro-*At-NF-YA2: GUS* were from Sorin et al. (2014) study. Seeds of *At-nf-ya2* (SALK_021228) were obtained from the Arabidopsis Biological Resource Center (Columbus, OH). Homozygous plants were identified by diagnostic PCR analysis using the primers listed in Table S6. Arabidopsis plants were grown in soilrite containing plastic pots and kept under cool white light of 100 μmol m−2 sec−1 intensity, with a 16 h light/8 h dark photoperiod at 22 °C/20 °C (day/night) in a plant growth chamber (CMP6050, Conviron, Canada).

One-month old tomato plants were initially exposed for 4 h at a gradual increasing range of temperature from 26 °C to 45 °C then exposed additionally for 4.5 h at 45 °C for HSFs and miR169s expression profiling study. VIGS silenced tomato plants were exposed to heat stress for 4.5 h at 45 °C while 2-weeks old *Arabidopsis* plants were exposed to 42 ᵒC for 2 h heat stress. For phenotype scoring and survival assays, HS treated VIGS silenced plants and *Arabidopsis* plants were allowed to recover for six days at 26 °C/21 °C (day/night: 16/8 h) and at 22 °C/20 °C (day/night) in a plant growth chamber (CMP6050, Conviron, Canada). Leaves from these experimental plants were harvested immediately after HS for expression profiling of HSR genes.

### Real-time quantitative RT–PCR analysis

Four (expression studies of *HSFs* and miR169s) and five weeks (VIGS silenced) old tomato plants and two-weeks-old seedlings of Arabidopsis were used for total RNA extraction with Trizol reagent (Invitrogen, USA). Total RNA was treated with DNase I (Ambion, USA) to remove genomic DNA contamination. For real-time RT–PCR (qRT–PCR) analysis, 2 μg of total RNA was used for cDNA synthesis by using high capacity cDNA reverse transcription kit (Applied Biosystem, USA) following the manufacturer’s protocol. The cDNA reaction mixture was diluted 2-fold, a 0.5 μl aliquot was used as a template in a 10 μl PCR reaction. PCR reactions included a pre-incubation at 95°C for 2 min, followed by 40 cycles of denaturation at 95°C for 15 sec, annealing at 60°C for 40 sec, and extension at 72°C for 45 sec. All the reactions were performed in a CFX-Connect real-time PCR detection system (Bio–Rad, www.bio-rad.com/) using GO Taq qPCR master mix (Promega, USA). Each experiment was repeated at least three times. The relative fold change was calculated by following the 2^−ΔΔCT^ method using *Actin* and *Tubulin* gene as endogenous control (Paul et al. 2016). The stability and similarity in the Ct range of *Actin* and *Tubulin* were assessed across the experimental conditions (Supplementary table S9). For calculation of fold change in the gene expression the control value was set as one for all qRT-PCR expression analysis. The primers used in this study are listed in Supplementary table S10 and all the qRT-PCR primers having a good amplification efficiency ranging between 90-110% were used (Supplementary table S11).

For miRNA expression analysis 2 µg of the small RNA was polyadenylated using the Poly(A) Tailing kit (epicenter-An Illumina company) following the manufacturer’s protocol. The above poly-A tailed small RNA reaction mixture was then reverse transcribed with a special 100 bp primer i.e. miR_oligodT_RTQ (Mutum et al. 2013) which is composed of two variable bases at its end followed by the poly(T)25 and then with a random adapter sequence using the Superscript III Reverse Transcriptase (Invitrogen, USA). The taqman based q-RT-PCR was performed using the TaqMan Fast Universal PCR Master Mix (2X, Applied Biosystems) and ‘Fam (Fluorescein) dye’ and BHQ labeled taqman universal probe specific to the adapter region of the miR_oligodT_RTQ (Supplementary table S8) which enable the use of a common reverse primer (Mutum et al. 2013) and miRNA specific forward primer in the expression analysis. For each well a total reaction volume of 7 µl was prepared and for each miRNA two technical with 3 biological replicates were analyzed, *5S rRNA* and *U6snRNA* (Supplementary table S8) were used as the endogenous control for data normalization. For calculation of fold change in the gene expression the control value was set as one for all qRT-PCR expression analysis.

### Promoter sequences analysis

The 2 kb sequences, upstream of the precursor start site of *MIR169* precursors and translational start codon (ATG) of Arabidopsis HSF genes were downloaded from Sol Genomic Network database (http://sgn.cornell.edu) and TAIR, respectively and analyzed for the presence of putative *cis*-regulatory elements by PlantPAN 3.0 (Chow et al. 2019). Positions of heat stress elements (HSE) were also searched manually in miRNA169 promoters. We searched for both perfect and imperfect HSE types manually, as these imperfect variants also have been validated for HSFs binding by different in vitro and in vivo assays (Santoro et al. 1998; Mittal et al. 2011; Guo et al. 2008).

### Yeast one-hybrid assays

The coding sequences for 23 tomato HSFs and 21 Arabidopsis HSFs were cloned into the pGADT7 vector (Clontech) as a prey (Supplementary table S8). DNA fragments corresponding to the promoters (250 to 1500 bp) of heat responsive *MIR169* genes from tomato and Arabidopsis (Supplemental tables S7, S3) were cloned into the pABAI plasmid (Clontech) as baits. For positive control, sHSP promoter previously reported to have HSFA3a binding (Li et al. 2013) was cloned. Promoter-specific bait yeast strains were generated by transforming Y1HGold yeast strain with 1 µg of linearized bait plasmids and plated on SD-ura medium. Positive bait strains having insertion at the URA3 locus were selected by performing colony PCR using primers specific to the bait sequences (Supplementary table S10). To negate this bait specific background *AbAr* expression, minimal inhibitory concentrations for all 27 baits (miRNA promoters) were determined by spotting yeast cultures at OD_600_ ranging from 0.1 to 0.0001on SD/-Ura medium supplemented with different concentration of aureobasidin A (100-1000 ng/ml; Supplementary tables S6, S12). For one-on-one bait/ prey interaction, bait strains were transformed with pGADT7 vector containing CDS of tomato HSFs. Positive bait/prey containing yeast colonies were selected by plating the transformed yeast cells on SD/-Leu medium. To study the interaction between different tomato HSFs and miRNA promoters, specific bait/prey containing yeast strains were spotted on SD/-Leu/AbA medium having bait specific aureobasidin A concentration.

### Chromatin immunoprecipitation (ChIP)-qPCR assays

Promoters for tomato *Sly-MIR169a-1, Sly-MIR169b, Siy-MIR169c, Sly-MIR169d* and *SlyMIR169d-1* were amplified from the genomic DNA and cloned into pbi101 vector upstream of the *GUS* reporter gene. All transcription factors (*Sly-HSFA1a*, *Sly-HSFA7a*, *Sly-HSFA8a*, *Sly-HSFB1a*, *Sly-HSFA4a*, *Sly-HSFA6b* and *Sly-HSFA1e* CDS) were cloned in pCAMBIA1302 in fusion with the *GFP* reporter. The binary constructs were co-infiltrated as per the combinations obtained from Y1H assay into 4-weeks-old *N*. *benthamiana* leaves via *Agrobacterium tumefaciens* (strain EHA105). These leaves were harvested 2 days after the infiltration and ChIP assays was performed using four infiltrated leaves from five independent plants for each co-infiltration. To identify the binding of At-NF-YA2 on HSR gene promoters, *NF-YA2:GFP* overexpressing transgenic lines were used. Leaves from *N*. *benthamiana* plants (for tomato ChIP) and 3-week-old *NF-YA2:GFP* overexpressing Arabidopsis plants were cross-linked with 1% formaldehyde. Chromatin was sheared to an average length of 500bp by sonication, and immunoprecipitated with GFP tag-specific monoclonal antibody (abcam; catalogue no. ab290). The fold enrichment was calculated for different promoters by performing real-time quantitative PCR analysis on immunoprecipitated DNA using a Bio–Rad CFX96 real-time PCR detection system. For each promoter pair used for ChIP-qPCR the primer amplification was first tested on input DNA and the fold enrichment was checked by keeping no-antibody, IgG and input DNA controls. Data for enrichment of Sly-HSFA6b on *Sly-MIR169d-1* promoter is presented as an example in Supplementary figure S24. All the primers used for ChIP-qPCR are listed in Supplementary Table S10.

### Transient promoter reporter assay to confirm HSFs-mediated regulation of *Sly-MIR169* promoters in *N. benthamiana*

Agro-infiltration was performed in 4-weeks-old *N*. *benthamiana* leaves. For promoter reporter assays Pro-*Sly-MIR169s*:GUS and p35S:*Sly-HSFs* were transformed in *Agrobacterium tumefaciens*. For co-infiltrations, overnight cultures of individual constructs were harvested and suspended at OD 1 in infiltration buffer (10 mM MgCl2, 10 mM MES, pH 5.8, 0.5% glucose and 150 μM acetosyringone). After 3 h of incubation at room temperature, various combinations were mixed at a 1:1:1 ratio; these mixes were used to infiltrate leaves. These leaves were harvested 2 days after the infiltration and used for RNA extraction and qRT-PCR as described above. The expression of GUS transcript is measured by qRT-PCR and presented as fold-change values calculated using the 2−ΔΔCt method. The values were normalized with *NPTII* that is co-expressed in the promoter: GUS construct. The GUS expression from the reporter (*pro-Sly-MIR169:GUS)* infiltrated samples was set to one to normalize the background signal.

### Virus induced gene silencing

Transient silencing assays following VIGS was performed in tomato plants as described by Senthil and Mysore (2014). For silencing of selected genes, 300 to 400 bp cDNA fragments designed using VIGS tool (Sol Genomics Network) were amplified and cloned into the *Eco* RI–*Bam* HI site of the TRV2 vector. After confirmation by Sanger sequencing, the TRV2-gene vectors were transformed into *A. tumefaciens* strain GV3101. About 15-days-old tomato plants were inoculated with a mixture of *A*. *tumefaciens* strains containing the TRV2 vector constructs and TRV1 (a helper plasmid, contains RNA-dependent RNA polymerase (RdRp), movement protein (MP), and a 16 kDa cysteine-rich protein) into the first and second true leaf. Empty TRV2 vector was used as a control. Infected plants were kept in dark and moist conditions for 3 days and then transferred to light; HS was imposed after 3 weeks (the time required for appearance of chlorophyll bleaching phenotype in the leaves of TRV-*Phytoene desaturase*-infiltrated plants used as silencing control) (Supplementary figure S25A, B) at 45 °C for 4.5 h. Leaves were frozen for qRT-PCR analysis. Off targets of the selected VIGS constructs were also predicted at VIGS tool (Supplementary table S4) and validated by qRT-PCR in the silenced plants. To confirm the maintenance of silencing throughout the phenotype scoring and biochemical assays time period, the expression of *Sly-NF-YA9* and *Sly-NF-YA10* was assessed further during the post heat stress period. Survival and cell death was assayed after 6 days’ recovery. The experiment was performed in four biological replicates with 8 plants per replicate for each gene.

### Gas-exchange parameters measurements

Gas-exchange parameters, like water use efficiency (WUEi), net photosynthetic rate (A), and transpiration rate (E) were measured simultaneously by using a portable Licor 6400 photosynthesis system (LI-6400, Li-Cor Inc., Lincoln NE, USA). WUEi was calculated using the formula A/E. For all measurements, sixth and seventh fully expanded tomato leaves of 6 plants for each VIGS construct were used and the experiment was repeated thrice keeping similar parameters. The measurement conditions were as follows: leaf temperature 27°C, leaf-air vapour pressure deficit 1.5±0.5 kPa, photosynthetic photon flux 300 μmol m^−2^ s^−1^, relative air humidity 60% and ambient CO2 concentration 400±5 μmolmol^−1^.

### Histochemical detection of H2O2 accumulation and cell death in HS

Hydrogen peroxide (H_2_O_2_) levels and cell death were estimated by using 3,3’-diaminobenzidine (DAB) and trypan blue stain, respectively on the 6th and 7th leaves of 6-week-old TRV-Vector and TRV-gene plants immediately after heat stress (45 °C for 6 h). DAB staining was performed as previously described (Daudi et al. 2012). The viability of leaf cells was measured using the trypan blue exclusion method (Koch and Slusarenko 1990). Experiment was repeated twice with similar parameters having a sample size of 8 plants per replicate.

### GUS assays

To analyze the histochemical localization of GUS reporter protein, tissue samples were incubated for 12 h at 37 °C with the substrate solution (1 mM 5-bromo-4-chloro-3-indolyl-β-D-glucuronide, pH 7.0, 100 mM sodium phosphate buffer, 10 mM Na2EDTA, 0.5 mM potassium ferricyanide, 0.5 mM potassium ferrocyanide, and 0.1% Triton X-100). Subsequently, seedlings and leaves were washed with de-staining solution containing ethanol:acetone:glycerol (3:1:1) to eliminate chlorophyll. They were photographed with a Nikon SMZ1000 Stereomicroscope (Tokyo, Japan) and OLYMPUS SZX16 Stereomicroscope (India). *GUS* transcripts were quantified using qRT-PCR. The assays were performed with three biological replicates.

### Constructs generation

For yeast one hybrid assay, all the promoters and CDS of transcription factors (HSFs and NF-YA) were cloned in pABAI and PGADT7 vector, respectively. For promoter reporter assays, all promoters were cloned in pbi101 vector and all transcription factors (*Sly-HSFA1a*, *Sly-HSFA7a*, *Sly-HSFA8a*, *Sly-HSFB1a*, *Sly-HSFA4a*, *Sly-HSFA6b* and *Sly-HSFA1e* CDS) were cloned in pCAMBIA1302. Promoters were amplified as approximately 2000 bp region upstream of the precursor start site for *MIR169s* (Pro-*Sly-MIR169a-1*:*GUS*, Pro-*Sly-MIR169b:GUS*, Pro-*Sly-MIR169c:GUS*, Pro-*Sly-MIR169d:GUS*, Pro-*Sly-MIR169e:GUS*, Pro-*Sly-MIR169d-1:GUS*, Pro-*At-MIR169d:GUS)* and translation start site for HSFs (Pro-*At-HSFA3a:GUS* and Pro-*At-HSFA7b:GUS),* Pro-*NF-YA2:GUS* and Pro-*Sly-NF-YA10:GUS* from genomic DNA. For overexpressing miR169d (*p35S:At-MIR169d-OE)* and sly-miR169d-5p (*p35S:Sly-MIR169d-OE),* 300 bp and 426bp fragment sequence surrounding the miRNA sequence was amplified from genomic DNA, cloned in pbi121 and transformed into Arabidopsis and tomato WT plants, respectively by floral dip method (Arabidopsis) and *Agrobacterium* mediated tomato transformation (Sun et al. 2015). For VIGS, 300-400 bp fragment of *Sly-HSFA1a*, *Sly-HSFA7a*, *Sly-HSFB1a*, *Sly-HSFA6b* and *Sly-NF-YA9*, *Sly-NF-YA10* were cloned in pTRV-2 vector.

### Measurement of physiological and biochemical parameters

For Physiological parameters, *Solanum lycopersicum* cv. Pusa Ruby (tomato) was used as wild type control. Thirty-days-old plants of transgenic lines and wild type plants were initially acclimated to heat stress with gradual increase in temperature from 26 °C to 45 °C for 2.5h and then exposed for 5h at 45 °C. Leaves from these experimental plants were harvested immediately after stress for measuring physiological parameters. For RWC measurement, fully expanded leaves were excised from control and heat stressed plants of respective overexpression lines and wild type plants were collected for their fresh weight was measured immediately while for EL measurement, leaves were cut into small pieces (1 cm^2^), immersed in 20 mL deionized water and incubated at 25°C and were further preceded as described by Houimli et al. (2010).Proline estimation was determined as Bates et al. (1973) in leaves of control and heat stressed respective overexpression lines and Pusa Ruby.

### Thermotolerance assay

To estimate the percentage survival rate of the tomato seedlings under heat stress, 4-days-old germination paper-grown seedlings of WT Pusa Ruby and *Sly-MIR169-OE* transgenic seedlings were subjected to 45 °C for 4.5 hours, the number of surviving seedlings was recorded and the percentage survival was calculated by dividing it with the total number of seedlings used for the treatment. Hypocotyl length was measured after 4days of recovery at 26^ᵒ^C post heat stress. The treated seedlings were photographed after 4 days of heat stress.

## Acknowledgments

This work is supported by grants from DBT-NIPGR, India and grants from DST-SERB, India (Project No.: EMR/2016/006229). The authors acknowledge phytotron facility, CIF and field area provided by NIPGR. The authors are thankful to DBT-eLibrary Consortium (DeLCON) for providing access to e-resources. SR acknowledges Department of Biotechnology (DBT) Govt. of India, SJ, CB and AG acknowledge University Grants Commission (UGC) and DST-INSPIRE for the award of research fellowships.

## Author contributions

SM designed and supervised the study; SR designed and performed all the experiments and analyzed the data; SJ generated Sly-MIR169-OE and promoter lines; CB did mutant genotyping for hsfa2 and nf-ya2; AG helped in conducting experiments with tomato transgenic lines; CS generated MIR169, NF-YA2 and NF-YA2-r transgenic lines of Arabidopsis. SR and SM wrote the article; CS and MC complemented the writing and critically reviewed the manuscript. SM agrees to serve as the author responsible for contact and ensures communication.

## Supplementary Figure legends

**Supplementary figure S1: Promoter analysis of *MIR169* genes in tomato.** Distribution of perfect and imperfect heat stress elements (HSE) in *MIR169* promoters of tomato. HSEs were identified manually by curating a list from published literature and marked as pink (perfect HSE) and turquoise blue (imperfect HSE) colored shapes on black colored lines representing *MIR169* promoters. The red right-handed arrow represents precursor start site. The HSE sequence variants for perfect and imperfect HSEs is provided in supplementary table S1.

**Supplementary figure S2: Yeast-one-hybrid assay of 24 tomato HSFs on *Sly-MIR169d-1* promoter.** Interaction of only Sly-HSFA6b on *Sly-MIR169d-1* promoter highlights the specificity of the Y1H assay. The co-transformed Y1H gold yeast strain cultures were spotted (O.D. 0.1 to 0.0001) onto plates lacking URA and LEU with and without specific Aureobasidin A concentration and incubated for 3 days. Promoter of *sHSP* has been used as a positive control to show binding of HSFA3a on *sHSP* promoter (Li et al. 2013) and empty vectors as negative controls. The positive binding of HSFA6b on *Sly-MIR169d-1* promoter was assessed at 200 ng/ml Aureobasidin A concentration.

**Supplementary figure S3: HSF-mediated transcriptional activation of tomato *MIR169* promoters.** *In-planta* transient assays showing HSF-mediated transcriptional activation of *Sly-MIR169:GUS* promoters in *Nicotiana benthamiana* as assessed by qRT-PCR. The fold-change of *GUS* transcripts were calculated using the 2−ΔΔCt method in (A) to (E). (A) Sly-HSFA1e with *pro-Sly-MIR169a-1:GUS*. (B) Sly-HSFA7a and Sly-HSFA1a with *pro-Sly-MIR169b:GUS*. (C) Sly-HSFA6b with *pro-Sly-MIR169c:GUS*. (D) Sly-HSFB1a with *pro-Sly-MIR169d:GUS*. (E) Sly-HSFA6b with *pro-Sly-MIR169d-1:GUS*. The *NPTII* gene co-expressed in the *pro-Sly-MIR169:GUS* construct was taken as the reference gene for normalization. The *GUS* expression from the reporter (*pro-Sly-MIR169:GUS*) infiltrated samples was set to one for normalization (marked as green bars). These experiments were repeated at least six times; the error bars represent standard deviation between the biological replicates.

**Supplementary figure S4: Expression profiling of HSFs:miRNAs:NF-YAs during heat stress in tomato leaves.** (A) Expression profiles of *HSF* genes that regulate *Sly-MIR169* transcription in WT tomato leaves during control and heat stress condition by qRT-PCR using the 2−ΔΔCt method. (B) Expression of mature miR169s in response to heat stress as determined by taqman based qRT-PCR in control and heat challenged WT tomato leaves. (C) qRT-PCR based expression profiles of target *NF-YA* transcripts in WT control and heat stressed tomato leaves using the 2−ΔΔCt method. Data represents mean values and standard deviation of biological replicates. *Tubulin* was used as endogenous control in (A) and (C), while *Sly-U6snRNA* was used as normalization control in (B). Error bars in (A), (B) and (C) represent the standard deviation of biological replicates. The fold change normalization was done by setting the control value as one for all qRT-PCR expression analysis.

**Supplementary figure S5: Determining the specificity of HSFs silencing in VIGS silenced tomato plants.** (A-C) qRT-PCR based expression profiles of putative off target genes using the 2−ΔΔCt method. (A) Relative expression of Sly-*HSFA1b* and Sly-*HSFA1c* off targets in TRV-*Sly-HSFA1a* silenced plants. (B) Relative expression of Sly-*HSFA6a* off target in *TRV-Sly-HSFA6b* silenced plants. (C) Relative expression of Sly-*HSFA4c* and Sly-*HSFB4a* off targets in *TRV-Sly-HSFA7a* silenced plants. Error bars in (A-C) represent the standard deviation of biological replicates. *ACTIN* was used as endogenous control in A-C. Data normalization and fold change values with *TUBULIN* were presented in Supplementary table S3. The fold change normalization was done by setting the TRV-EV expression as one for all qRT-PCR expression analysis.

**Supplementary figure S6: In planta validation of heat governed HSFs mediated transcriptional regulation of MIR169s and NF-YAs in tomato leaves.**

The expression analysis of *HSF* genes, *Sly-MIR169* precursors and *NF-YA* transcripts in VIGS plants silenced for *Sly-HSFA1a* (A), *Sly-HSFA7a* (B), *Sly-HSFA6b* (C) and *Sly-HSFB1a* (D). 15-days-old tomato plants were agro-infiltrated with empty vector (TRV-EV) or TRV-Sly-HSF VIGS constructs. The VIGS established plants were subjected to heat stress after 3-weeks-of agro-infiltration, and used for expression studies of different genes. Graphical data represents mean values of expression of three to four biological sets. Error bars show standard deviation. To represent the negative fold-change on y axis, the relative expression values of down-regulated genes were transformed using the formula [-(1 / RQ ≤0.5)]. *Tubulin* was used as endogenous reference control. The fold change normalization was done by setting the control value as one for all qRT-PCR expression analysis.

**Supplementary figure S7: Physiological, Biochemical and Molecular characterization of *Sly-MIR169-OE* lines.** (A) Estimation of relative water content in WT and *Sly-MIR169-OE* (three independent lines #1,2,3) transgenic plants during non-stressed and heat stressed conditions. (B) Quantification of electrolyte leakage in WT and *Sly-MIR169-OE* (three independent lines #1,2,3) transgenic plants during non-stressed and heat stressed conditions. (C) Estimation of proline content in WT and *Sly-MIR169-OE* (three independent lines #1,2,3) transgenic plants during non-stressed and heat stressed conditions. (D) DAB and trypan blue staining of WT and *Sly-MIR169-OE* transgenic plants during heat stressed conditions. (E) qRT-PCR based expression profiles of HSR genes in WT and *Sly-MIR169-OE* transgenic plants. Error bars represent the standard deviation of biological replicates in (A, B, C and E). The experiments were repeated at least three times with similar results, and data from one representative experiment are shown in (D). Error bars show standard deviation. *p<0.05, **p<0.01 and ***p<0.001 vs. wild type, by two-tailed Student’s t-test.

**Supplementary figure S8: Virus induced silencing of Sly-*NF-YA9* and Sly-*NF-YA10*.** (A) qRT-PCR analysis of Sly-*NF-YA9* and Sly-*NF-YA10* gene in TRV-EV, *TRV-Sly-NF-YA9* and *TRV-Sly-NF-YA10* plants in control (non-stressed) conditions confirming their silencing. The fold change normalization was done by setting the TRV-EV expression as one for all qRT-PCR expression analysis. (B) Expression of Sly-*NF-YA9* and Sly-*NF-YA10* in WT tomato and TRV-EV infiltrated plants in control (non-stressed) conditions. The fold change normalization was done by setting the WT control value as one for all qRT-PCR expression analysis (C-D) Expression profiles of *Sly-NF-YA9* (C) and *Sly-NFY-A10* (D) in *TRV-Sly-NF-YA9/A10* VIGS silenced plants in heat stress (HS) in comparison to control non-stressed conditions. The fold change normalization was done by setting the control value as one for all qRT-PCR expression analysis. The expression levels of genes were calculated using the 2−ΔΔCt method and presented using fold-change values. *Actin* was used for normalising expression in (A-D). Fold change values for (A-D) with *TUBULIN* normalisation were presented in Supplementary table S3. EV: vector control. Error bars represent the standard deviation of biological replicates.

**Supplementary figure S9: Determining the specificity of NF-YA9/A10 silencing in VIGS silenced tomato plants.** (A-B) qRT-PCR based expression profiles of putative off target genes using the 2−ΔΔCt method. (A) Relative expression of Sly-*NF-YA1, Sly-NF-YA3* and Sly-*NF-YA10* off targets in TRV-*Sly-NF-YA9* silenced plants. (B) Relative expression of *Sly-NF-YA9 (*no off targets were predicted for TRV-Sly-NF-YA10, we used *Sly-NF-YA9* to show specificity of silencing) as in *TRV-Sly-NF-YA10-VIGS* silenced plants. Error bars in (A-B) represent the standard deviation of biological replicates. *ACTIN* was used as endogenous control in A-B. Fold change values for (A-B) with *Tubulin* normalisation were presented in Supplementary table S3. The fold change normalization was done by setting the TRV-EV expression as one for all qRT-PCR expression analysis.

**Supplementary figure S10: Expression analysis of heat stress responsive genes in control and heat treated TRV-Sly-*NF-YA9* and TRV-Sly-*NF-YA10* silenced tomato plants.**

(A-E) Expression profiles of HSR genes in control and heat treated TRV-EV, *TRV-Sly-NF-YA9* and *TRV-Sly-NF-YA10* silenced plants. qRT-PCR based fold change in expression in control and heat stressed plants of *Sly-HSFA2* (A); *Sly-HSFA7a* (B); *Sly-HSFA3a* (C); *Sly-APX* (D) and *Sly-HSP17.6 CII*. (E). Relative expression of HSR genes in *TRV-Sly-NF-YA9* and *TRV-Sly-NF-YA10* silenced plants in control conditions as compared to *TRV-EV* plants. (G) Expression profiles of HSR genes in control and heat treated *TRV-Sly-NF-YA9* and *TRV-Sly-NF-YA10* silenced plants. The expression levels of genes were calculated using the 2−ΔΔCt method and presented using fold-change values. *Tubulin* was used as reference gene for A-E and *Actin* was used as endogenous control for F-G. Fold change values for (F-G) with *TUBULIN* normalisation were presented in Supplementary table S3 The fold change normalization was done by setting the expression in control non-stressed as one for all qRT-PCR expression analysis in (A-E). The fold change normalization was done by setting the TRV-EV heat stressed expression as one for all qRT-PCR expression analysis in (F). The fold change normalization was done by setting the expression of TRV-EV in control non-stressed as one for all qRT-PCR expression analysis in (G). Error bars represent the standard deviation of independent biological replicates.

**Supplementary figure S11: Expression profiling of At-miR169s in heat stress.** Expression of mature At-miR169s in response to heat stress as determined by Taqman-based qRT-PCR. *U6snRNA* was used as the endogenous normalization control. The average values of multiple biological replicates are plotted as bars, error bars depict standard deviation between three replicates. The fold change normalization was done by setting the control value as one for all qRT-PCR expression analysis.

**Supplementary figure S12: Expression analysis of At-MIR169d in heat stress in Arabidopsis**. Expression levels of *At-MIR169d* in wild-type plants subjected to 0 (control) and 2 h heat stress at 42 °C. Data are shown as means ± SE of three biological replicates and normalized to *At-EF-α* reference gene using (2−ΔΔCT) method. The fold change normalization was done by setting the control value as one for the qRT-PCR expression analysis.

**Supplementary figure S13: Schematic structure of *At-*HSFA2 gene carrying the T-DNA insertion and lack of *At-*HSFA2 expression in the *At-hsfA2* mutant (Salk_008978).** (A) The two exons of *At-*HSFA2 genomic DNA are shown according to the gene features annotated at TAIR database. T-DNA insertion site is shown as inverted triangle in the second exon of *At-*HSFA2. The exon sequences are depicted by red and the UTRs are represented by green color. (B) Genotyping PCR of *At-hsfa2* mutants using genomic DNA, presence of border specific amplification in *At-hsfa2* confirms their homozygosity, while presence of only gene specific amplification in control Columbia plants confirms their wild type nature. (C) Expression of *At-HSFA2* was determined by RT-PCR, using full length cDNA from WT and mutant *At-hsfa2* plants. Absence of *At-HSFA2* specific band in mutant plants confirms the knockout nature of *At-hsfa2* homozygous mutants. RNA was isolated from detached mature leaves of the wild-type (wt) or *At-hsfa2* plants. Expression of *actin* gene is shown as a loading control.

**Supplementary figure S14: Expression pattern of miRNA169d:NF-YA in Arabidopsis.** (A) Expression levels of mature miR169d and eight miR169 target genes in wild-type plants subjected to 0 (control) and 2 h heat stress at 42°C using qRT-PCR. Data are shown as means ± SD of biological replicates and normalized to *actin* and *5SrRNA* reference genes, for mRNA and miRNA respectively, using (2^−ΔΔCT^) method for A and with *At-EF-α* reference gene and *U6snRNA* for mRNA and miRNA respectively, using (2^−ΔΔCT^) method for B. The fold change normalization was done by setting the control value as one for all qRT-PCR expression analysis.

**Supplementary figure S15: Establishing miRNA169:target modules operating in Arabidopsis.** (A) Taqmann based qRT-qPCR analysis of the accumulation of mature miR169defg and miR171bc (used as negative control) in Arabidopsis MIM169defg lines. Average values of expression for different miR169s in three independent lines of MIM169defg transgenic plants were plotted as bars and error bars deoicts the standard deviation between the replicates. (B) qRT-PCR analysis of the accumulation of the *At-NF-YA2* transcripts in MIM169defg Arabidopsis lines. Data are presented as average fold induction relative to control where induction is the ratio of transcript levels in two independent MIM169defg transgenic lines to those in wild-type plants (WT). Error bars, ± SD of three repeats. (C) qRT-PCR analysis of the reduction of *At-NF-YA2* transcripts in three independent miR169d over expressing (OE) lines. Data are presented as average fold in two independent *MIM169defg* lines relative to WT control, that was set as one.

**Supplementary figure S16: Heat stress mediated transcriptional regulation of *At-NF-YA2*.** (A) Histochemical GUS expression of Pro-*At-NF-YA2:GUS* transgenic Arabidopsis plants during control and heat stress. (B) qRT-PCR based determination of *GUS* transcripts of Pro-*At-NF-YA2:GUS* transgenic Arabidopsis plants during control and heat stress. These experiments were repeated 3 times with 20 seedling per replicate. Error bars depict standard deviation between three replicates. The fold change normalization was done by setting the control value as one for qRT-PCR expression analysis.

**Supplementary figure S17: Delineating the existence and localization of miR169d:At-NF-YA2 functional module in Arabidopsis.** Histochemical GUS expression patterns of *Pro-At-MIR169d:GUS and Pro-At-NF-YA2:GUS* promoter reporter transgenic Arabidopsis plants in: seedling (2 weeks old), complete rosette (4 weeks), mature rosette leaf, cauline leaf and flower

**Supplementary figure S18: Expression levels of *At-NF-YA2* in *At-NF-YA2* overexpressing transgenic plants.** Relative expression of *At-NF-YA2* in transgenic Arabidopsis plants overexpressing *At-NF-YA2* under constitutive CaMV35S promoter. Bar represents the average data of three biological replicates and error bar represents standard deviation of the replicates.

**Supplementary figure S19: Schematic structure of *At-NF-YA2* gene carrying the T-DNA insertion and lack of *At-NF-YA2* expression in the *At-nf-ya2* mutant (SALK_021228).** (A) The four exons of *At-NF-YA2* genomic DNA are shown according to the gene features annotated at TAIR database. T-DNA insertion site is shown by inverted triangle in the fourth exon of *At-NF-A2*. The exon sequences are depicted by red and the UTR are represented by green color. (B) Genotyping PCR of *At-nf-ya2* mutants, presence of border specific amplification in *At-nf-ya2* plants genomic DNA confirms their homozygosity, while presence of only gene specific amplification in control Columbia plants confirms their wild type nature. (C) Expression of *At-NF-YA2* was determined by RT-PCR, using full length cDNA from WT and mutant *At-nf-ya2* plants. Absence of *At-NF-YA2* specific band in mutant plants confirms the knockout nature of *At-nf-ya2* homozygous mutants. RNA was isolated from detached mature leaves of the wild-type (WT) or *At-nf-ya2* plants. Expression of actin is shown as a loading control.

**Supplementary figure S20: Expression profiling of HSR genes in different transgenic lines of Arabidopsis.** Expression patterns of five heat stress-responsive genes in WT, *At-MIR169d-OE*, *At-NF-YA2-OE*, *At-nf-ya2, pNF-YA2:NF-YA2*-r and MIM169defg plants. Data are shown as means ± SD of biological replicates and normalized to *At-EF-α* reference gene using (2^−ΔΔCT^) method. The fold change normalization was done by setting the control value as one for qRT-PCR expression analysis.

**Supplementary figure S21: Yeast-one-hybrid assay based transcriptional regulation of *At-HSFA3a, At-HSFA7a* and *At-HSFA7b* promoters by At-NF-YA2**. Positive interaction of At-NF-YA2 with the promoters of *At-HSFA3a* and *At-HSFA7b* were obtained. Serial dilutions (O.D. 0.1 to 0.0001) of yeast cultures were spotted onto plates lacking LEU with specific Aureobasidin A concentration and incubated for 3 days.

**Supplementary figure S22: Thermotolerance assay of WT and *At-hsfa7a* plants**. (A) Two-weeks-old soil-grown WT and *At-hsfa7a* mutant plants were subjected to 0 (control) or 2h heat stress at 42°C, and damage was recorded 6 days later. (B) Estimation of survival (percentage) of heat stress treated WT and *At-hsfa7a* plants. Plants were assayed for heat stress tolerance after 6 days of HS. These experiments were repeated at least three times with similar results, and average data of all experiments are shown. Error bars, ± SD of three repeats. ***p<0.001 vs. wild type, by two-tailed Student’s t-test.

**Supplementary figure S23: At-NF-YA2, Arabidopsis orthologue of tomato Sly-NF-YA9 and Sly-NF-YA10.** (A-B) Protein sequence alignment of tomato Sly-NF-YA9 (A) and Sly-NF-YA10 with At-NF-YA2. Protein sequences were aligned with Arabidopsis proteome at TAIR database by using Blastp tool.

**Supplementary figure S24:** Binding of *Sly-HSFA6b* to Sly-MIR169d-1 promoter. Absence of any amplification in no-Ab (no antibody) and slight amplification in IgG control confirms the specificity of immunoprecipitation. Presence of strong amplification on input DNA confirms the good amplification efficiency of the primer used for analysis. Presence of amplification in Anti-GFP lane confirms the enrichment of Sly-MIR169d-1 promoter in *Sly-HSFA6b:GFP* CHIP. L denotes the 100 bp ladder.

**Supplementary figure 25: Virus induced silencing of tomato *Phytoene desaturase* (*PDS*) gene.** *PDS* silencing was used as a control for assessing establishment of gene silencing by VIGS. (A) Silencing of the PDS control gene causing photobleaching in tomato plants 3 weeks after agroinfection. (B) Enlarged individual leaves of TRV-PDS and control WT leaves.

## Supplementary Table legends

**Supplementary table S1:** Distribution of perfect and imperfect heat stress elements (HSE) in*Sly-MIR169* promoters.

**Supplementary table S2:** Interaction summary of *Sly-MIR169* promoters and Sly-HSFs. Positive HSF interaction is marked in red colored boxes.

**Supplementary table S3:** All Qrt-PCR raw data, calculations and fold change normalizations with different endogenous controls.

**Supplementary table S4:** Off targets of candidate VIGS constructs used in the study

**Supplementary table S5:** Promoter analysis of HSR genes in tomato.

**Supplementary table S6:** Number of HSE elements and minimal inhibitory concentration of Aureobasidin (Aba) for all At-MIR169 promoters

**Supplementary table S7:** Interaction summary of At-MIR169 promoters and At-HSFs. Positive HSF interaction is marked in red colored boxes.

**Supplementary table S8:** qRT-PCR based expression profiles of HSR genes in *At-MIR169d OE, At-NF-YA2 OE, At-nf-ya2, At-pNF-YA2:NF-YA2-r* and *MIM169defg* transgenic lines.

**Supplementary table S9:** Analysis of Ct similarities across different conditions for the reference genes used in the study.

**Supplementary table S10:** List of Primers used in the study.

**Supplementary table S11:** Primer efficiency of all the primers used in the study.

**Supplementary table S12:** Minimal inhibitory concentration of Aureobasidin (Aba) for all *Sly-MIR169* promoters.

## References

Balyan S, Rao S, Jha S, Bansal C, Das JR and Mathur S (2020). Characterization of novel regulators for heat stress tolerance in tomato from Indian sub-continent. Plant Biotechnology Journal, doi:10.1111/pbi.13371.

Bates LS, Waldren RP and Teare ID (1973) Rapid determination of free proline for water-stress studies. Plant and soil, 39(1), 205–207.

Bäurle I (2016) Plant heat adaptation: priming in response to heat stress. F1000Research, 5.

Bharti K, Von Koskull-Döring P, Bharti S, Kumar P, Tintschl-Körbitzer A, Treuter E, & Nover L (2004) Tomato heat stress transcription factor HsfB1 represents a novel type of general transcription coactivator with a histone-like motif interacting with the plant CREB binding protein ortholog HAC1. The Plant cell, 16(6):1521–1535.

Bonner J J, Ballou C, Fackenthal D L (1994) Interactions between DNA-bound trimers of the yeast heat shock factor. Molecular Cell Bioogyl,14:501–508.

Buhtz A, Pieritz J, Springer F, Kehr J (2010) Phloem small RNAs, nutrient stress responses, and systemic mobility. BMC Plant Biology, 10(1), 64.

Busch W, Wunderlich M, Schöffl F (2005) Identification of novel heat shock factor-dependent genes and biochemical pathways in Arabidopsis thaliana. The Plant Journal, 41:1–14.

Ceribelli M, Dolfini D, Merico D, Gatta R, Viganò AM, Pavesi G, Mantovani R (2008) The histone-like NF-Y is a bifunctional transcription factor. Molecular and cellular biology, 28:2047–2058.

Chan-Schaminet KY, Baniwal SK, Bublak D, Nover L and Scharf KD (2009) Specific interaction between tomato HsfA1 and HsfA2 creates hetero-oligomeric superactivator complexes for synergistic activation of heat stress gene expression. Journal of Biological Chemistry, 284:20848–20857.

Charng YY, Liu HC, Liu NY, Chi WT, Wang CN, Chang SH, Wang TT (2007) A heat-inducible transcription factor, HsfA2, is required for extension of acquired thermotolerance in Arabidopsis. Plant physiology, 143(1):251–262.

Chow CN, Lee TY, Hung YC, Li GZ, Tseng KC, Liu YH, Kuo PL, Zheng HQ, Chang WC (2019) PlantPAN3. 0: a new and updated resource for reconstructing transcriptional regulatory networks from ChIP-seq experiments in plants. Nucleic acids research, 47(D1): D1155–63.

Collins GG, Nie X, Saltveit ME (1995) Heat shock proteins and chilling sensitivity of mung bean hypocotyls. Journal of Experimental Botany, 46:795–802.

Czarnecka-Verner E, Yuan CX, Scharf KD, Englich G, Gurley WB (2000) Plants contain a novel multi-member class of heat shock factors without transcriptional activator potential. Plant Molecular Biology. 43:459–471.

Daudi A, O’Brien JA (2012) Detection of Hydrogen Peroxide by DAB Staining in Arabidopsis Leaves. Bio Protocol, 2(18),1–4.

Ding Q, Zeng J, He XQ (2016) MiR169 and its target PagHAP2-6 regulated by ABA are involved in poplar cambium dormancy. Journal of plant physiology, 198, 1–9.

Guan Q, Lu X, Zeng H, Zhang Y, Zhu J (2013) Heat stress induction of miR398 triggers a regulatory loop that is critical for thermotolerance in Arabidopsis. The Plant Journal, 74:840–851.

Guo L, Chen S, Liu K, Liu Y, Ni L, Zhang K and Zhang L (2008) Isolation of heat shock factor HsfA1a-binding sites in vivo revealed variations of heat shock elements in Arabidopsis thaliana. Plant and Cell Physiology, 49(9), 1306–1315.

Guo M, Liu JH, Ma X, Luo DX, Gong ZH, Lu MH (2016) The plant heat stress transcription sactors (HSFs): structure, regulation, and function in response to abiotic stresses. Frontiers in Plant Science, 7:114.

Hahn A, Bublak D, Schleiff E, Scharf KD (2011) Crosstalk between Hsp90 and Hsp70 Chaperones and Heat Stress Transcription Factors in Tomato. Plant Cell, 23:741–55.

Hasanuzzaman M, Nahar K, Alam MM, Roychowdhury R, Fujita M (2013) Physiological, biochemical, and molecular mechanisms of heat stress tolerance in plants. International Journal of Molecular Sciences, 14:9643–9684.

Hivrale V, Zheng Y, Puli C, Jagadeeswaran G, Gowdu K, Kakani V, Barakat A, Sunkar R (2015). Characterization of drought- and heat-responsive microRNAs in switchgrass. Plant Science, 242, 10:1016.

Houimli SIM, Denden M and Mouhandes BD (2010) Effects of 24-epibrassinolide on growth, chlorophyll, electrolyte leakage and proline by pepper plants under NaCl-stress. EurAsian Journal of BioSciences, 4(1), 96–104.

Huang YC, Niu CY, Yang CR, Jinn TL (2016) The heat stress factor HSFA6b connects ABA signaling and ABA-mediated heat responses. Plant Physiology, 172, 1182–1199.

Ikeda M, Ohme-Takagi M (2009) A novel group of transcriptional repressors in Arabidopsis. Plant and Cell Physiology 50:970–975.

Jacob P, Hirt H, Bendahmane A (2017) The heat-shock protein/chaperone network and multiple stress resistance. Plant Biotechnology Journal, 15:405–414.

Koch E, Slusarenko A (1990) Arabidopsis is susceptible to infection by a downy mildew fungus. The Plant Cell, 1;2(5):437–45.

Kotak S, Port M, Ganguli A, Bicker F and Von Koskull-Döring P (2004) Characterization of C-terminal domains of Arabidopsis heat stress transcription factors (Hsfs) and identification of a new signature combination of plant class A Hsfs with AHA and NES motifs essential for activator function and intracellular localization. The Plant Journal, 39(1), 98–112.

Kumimoto RW, Zhang Y, Siefers N, Holt III BF (2010) NF–YC3, NF–YC4 and NF–YC9 are required for CONSTANS-mediated, photoperiod-dependent flowering in Arabidopsis thaliana. The Plant Journal, 63(3), 379–391.

Larkindale J, Vierling E (2008) Core genome responses involved in acclimation to high temperature. Plant Physiol, 146:748–761.

Lee H, Yoo SJ, Lee JH, Kim W, Yoo SK, Fitzgerald H, Carrington JC, Ahn JH (2010) Genetic framework for flowering-time regulation by ambient temperature-responsive miRNAs in Arabidopsis. Nucleic acids research, 38: 3081–3093.

Leyva-Gonzalez MA, Ibarra-Laclette E, Cruz-Ramoırez A, Herrera-Estrella L (2012) Functional and transcriptome analysis reveals an acclimatization strategy for abiotic stress tolerance mediated by Arabidopsis NF-YA family members. PLoS ONE, 7: e48138.

Li S, Liu J, Liu, Z, Li X, Wu F, and He Y (2014) HEAT-INDUCED TAS1 TARGET1 Mediates Thermotolerance via HEAT STRESS TRANSCRIPTION FACTOR A1a-Directed Pathways in Arabidopsis. The Plant Cell, 26:1764–1780.

Li WX, Oono Y, Zhu J, He XJ, Wu JM, Iida K, Lu XY, Cui X, Jin H, Zhu JK (2008) The Arabidopsis NFYA5 transcription factor is regulated transcriptionally and post-transcriptionally to promote drought resistance. Plant Cell, 20(8):2238–2251.

Li Y, Fu Y, Ji L, Wu C-A, Zheng C (2010) Characterization and expression analysis of the Arabidopsis mir169 family. Plant Science, 178:271–280.

Li Y, Zhao SL, Li JL, Hu XH, Wang H, Cao XL, Xu YJ, Zhao ZX, Xiao ZY, Yang N, Fan J, Huang F, Wang WM (2017) Osa-miR169 negatively regulates rice immunity against the blast fungus Magnaportheoryzae. Frontiers in Plant Science, 8,2.

Li Z, Zhang L, Wang A, Xu X, Li J (2013) Ectopic overexpression of SlHsfA3, a heat stress transcription factor from tomato, confers increased thermotolerance and salt hypersensitivity in germination in transgenic Arabidopsis. PloS one, 8(1), e54880.

Lin JS, Kuo CC, Yang IC, Tsai WA, Shen YH, Lin CC, Liang YC, Li YC, Kuo YW, King YC, Lai HM, Jeng ST (2018) MicroRNA160 Modulates Plant Development and Heat Shock Protein Gene Expression to Mediate Heat Tolerance in Arabidopsis. Frontiers in Plant Science,9:68.

Liu HC, Liao HT, Charng YY (2011) The role of class A1 heat shock factors (HSFA1s) in response to heat and other stresses in Arabidopsis. Plant Cell and Environment, 34:738–51.

Lobell DB, Schlenker W, Costa-Roberts J (2011) Climate trends and global crop production since 1980. Science, 333:616–620.

Luan M, Xu M, Lu Y, Zhang L, Fan Y, and Wang L (2015) Expression of zma-miR169 miRNAs and their target ZmNF-YA genes in response to abiotic stress in maize leaves. Gene, 555:178–185.

Mittal D, Enoki Y, Lavania D, Singh A, Sakurai H and Grover A (2011) Binding affinities and interactions among different heat shock element types and heat shock factors in rice (Oryza sativa L.). The FEBS journal, 278(17), 3076–3085.

Mittler R, and Blumwald E (2010) Genetic engineering for modern agriculture: challenges and perspectives. Annual review of plant biology, 61:443–462.

Morimoto RI (2002) Dynamic remodeling of transcription complexes by molecular chaperones. Cell 110:281–284.

Mutum RD, Balyan SC, Kansal S, Agarwal P, Kumar S, Kumar M and Raghuvanshi S (2013). Evolution of variety-specific regulatory schema for expression of osa-miR408 in indica rice varieties under drought stress. The FEBS journal, 280(7),1717–1730.

Ni Z, Hu Z, Jiang Q, and Zhang H (2013) GmNFYA3, a target gene of miR169, is a positive regulator of plant tolerance to drought stress. Plant Molecular Biology, 82:113–129.

Niu C, Li H, Jiang L, Yan M, Li C, Geng D, Xie Y, Yan Y, Shen X, Chen P, Dong J, Ma F, Guan Q (2019) Genome-wide identification of drought-responsive microRNAs in two sets of Malus from interspecific hybrid progenies. Horticulture Research, 75:6.

Nover L, Bharti K, Döring P, Mishra SK, Ganguli A, Scharf KD (2001) Arabidopsis and the heat stress transcription factor world: how many heat stress transcription factors do we need? Cell Stress Chaperons, 6:177–189.

Pant BD, Musialak-Lange M, Nuc P, May P, Buhtz A, Kehr J, Walther D, Scheible WR (2009) Identification of nutrient-responsive Arabidopsis and rapeseed microRNAs by comprehensive real-time polymerase chain reaction profiling and small RNA sequencing. Plant physiology, 150:1541–1555.

Paul A, Rao S and Mathur S (2016). The α-crystallin domain containing genes: identification, phylogeny and expression profiling in abiotic stress, phytohormone response and development in tomato (Solanum lycopersicum). Frontiers in plant science, 7, 426.

Rao S, Balyan S, Jha S and Mathur S (2020) Novel insights into expansion and functional diversification of MIR169 family in tomato. Planta, 251:55.

Ravichandran S, Ragupathy R, Edwards T, Michael D, Sylvie C (2019) MicroRNA-guided regulation of heat stress response in wheat. BMC Genomics, 20:488.

Sakuma Y, Maruyama K, Qin F, Osakabe Y, Shinozaki K, Yamaguchi-Shinozaki K (2006) Dual function of an Arabidopsis transcription factor DREB2A in water-stress-responsive and heat-stress-responsive gene expression. Proceedings of the National Academy of Sciences, 103(49):18822–7.

Santoro N, Johansson N and Thiele DJ (1998) Heat shock element architecture is an important determinant in the temperature and transactivation domain requirements for heat shock transcription factor. Molecular and cellular biology, 18(11), 6340–6352.

Sato H, Mizoi J, Tanaka H, Maruyama K, Qin F, Osakabe Y, Morimoto K, Ohori T, Kusakabe K, Nagata M, Shinozaki K (2014) Arabidopsis DPB3-1 a DREB2A interactor, specifically enhances heat stress-induced gene expression by forming a heat stress-specific transcriptional complex with NF-Y subunits. The Plant Cell, 26 4954–4973.

Scharf KD, Berberich T, Ebersberger I, Nover L (2012) The plant heat stress transcription factor (Hsf) family: structure, function and evolution. Biochimica et Biophysica Acta (BBA)-Gene Regulatory Mechanisms,1819:104–119.

Senthil-Kumar M, Mysore K (2014) Tobacco rattle virus–based virus-induced gene silencing in *Nicotiana benthamiana*. Nature Protocols, 9:1549–1562.

Sorin C, Declerck M, Christ A, Blein T, Ma L, Lelandais-Brière C, Njo MF, Beeckman T, Crespi M and Hartmann C (2014) A miR169 isoform regulates specific NF-YA targets and root architecture in Arabidopsis. New Phytologist, 202:1197–1211.

Stief A, Altmann S, Hoffmann K, Pant BD, Scheible W, Bäurle I (2014) Arabidopsis miR156 regulates tolerance to recurring environmental stress through SPL transcription factors. The Plant Cell, 26:1792–1807.

Sun S, Kang XP, Xing XJ, Xu XY, Cheng J, Zheng SW and Xing GM (2015) Agrobacterium-mediated transformation of tomato (Lycopersicon esculentum L. cv. Hezuo 908) with improved efficiency. Biotechnology & Biotechnological Equipment, 29(5), 861–868.

Sunkar R, Zhu JK (2004) Novel and stress-regulated microRNAs and other small RNAs from Arabidopsis. The Plant Cell, 16:2001–2019.

Todesco M, Rubio-Somoza I, Paz-Ares J, Weigel D (2010) A collection of target mimics for comprehensive analysis of microRNA function in Arabidopsis thaliana. PLoS genetics, 6: e1001031.

Topol J, Ruden D M, Parker C S (1985) Sequences required for *in vitro* transcriptional activation of a *Drosophila hsp 70* gene. Cell, 42:527–537.

Vidya S, Ravishankar K, Laxman R (2018) Genome wide analysis of heat responsive microRNAs in banana during acquired thermo tolerance. Journal of Horticultural Sciences, 13:61–71.

Volkov RA, Panchuk II, Mullineaux PM, Schöffl F (2006) Heat stress-induced H2O2 is required for effective expression of heat shock genes in Arabidopsis. Plant Molecular Biology, 61, 733–746.

Von Koskull-Döring P, Scharf KD, and Nover L (2007) The diversity of plant heat stress transcription factors. Trends in Plant Science, 12:452–457.

Warpeha KM, Upadhyay S, Yeh J, Adamiak J, Hawkins SI, Lapik YR, Anderson MB, Kaufman LS (2007) The GCR1, GPA1, PRN1, NF-Y signal chain mediates both blue light and abscisic acid responses in Arabidopsis. Plant physiology, 143(4), 1590–1600.

Wu Z, Liang J, Wang C, Zhao X, Zhong X, Cao X, Li G, He J, Yi M (2018) Overexpression of lily HsfA3 s in Arabidopsis confers increased thermotolerance and salt sensitivity via alterations in proline catabolism. Journal of experimental botany, 69(8), 2005–2021.

Xiao H, Perisic O, Lis J T (1991) Cooperative binding of *Drosophila* heat shock factor to arrays of a conserved 5 bp unit. Cell, 64:585–593.

Yan K, Liu P, Wu CA, Yang G-D, Xu R, Guo Q-H, Huang J-G, Zheng C-C (2012) Stress-induced alternative splicing provides a mechanism for the regulation of microRNA processing in Arabidopsis thaliana. Molecular Cell, 48:521–531.

Yang X, Zhu W, Zhang H, Liu N, Tian S (2016) Heat shock factors in tomatoes: genome-wide identification, phylogenetic analysis and expression profiling under development and heat stress. PeerJournal, 4: e1961.

Yoshida T, Ohama N, Nakajima J, Kidokoro S, Mizoi J, Nakashima K, Maruyama K, Kim JM, Seki M, Todaka D, Osakabe Y (2011) Arabidopsis HsfA1 transcription factors function as the main positive regulators in heat shock-responsive gene expression. Molecular Genetics and Genomics, 286, 321–332

Yoshida T, Sakuma Y, Todaka D, Maruyama K, Qin F, Mizoi, J, Kidokoro S, Fujita Y, Shinozaki K and Yamaguchi-Shinozaki K (2008) Functional analysis of an Arabidopsis heat-shock transcription factor HsfA3 in the transcriptional cascade downstream of the DREB2A stress-regulatory system. Biochemical and biophysical research communications, 368(3), 515–521.

Zang D, Wang J, Zhang X, Liu Z and Wang Y (2019) Arabidopsis heat shock transcription factor HSFA7b positively mediates salt stress tolerance by binding to an E-box-like motif to regulate gene expression. Journal of experimental botany, 70(19), 5355–5374.

Zhang B (2015) MicroRNA: a new target for improving plant tolerance to abiotic stress. Journal of experimental botany, 66 1749–1761.

Zhang X, Zou Z, Gong P, Zhang J, Ziaf K, Li H, Xiao F, Ye Z (2011) Over-expression of microRNA169 confers enhanced drought tolerance to tomato. Biotechnology Letters, 33(2):403–409

Zhao B, Ge L, Liang R, Li W, Ruan K, Lin H and Jin Y (2009) Members of miR-169 family are induced by high salinity and transiently inhibit the NF-YA transcription factor. BMC Molecular Biology, 10(1): e29.

Zhao BT, Liang RQ, Ge LF, Li W, Xiao HS, Lin HX, Ruan KC and Jin YX (2007) Identification of drought-induced microRNAs in rice. Biochemical and biophysical research communications, 354:585–590.

Zhao M, Ding H, Zhu JK, Zhang F, Li WX (2011) Involvement of miR169 in the nitrogen-starvation responses in Arabidopsis. New Phytologist, 190:906–1.

Zhou X, Wang G, Sutoh K, Zhu JK, Zhang W (2008) Identification of cold-inducible microRNAs in plants by transcriptome analysis. Biochimica et Biophysica Acta (BBA)-Gene Regulatory Mechanisms,11:780–8.

