## Supplementary Figures for "A conserved HSF:miR169:NF-YA loop involved in tomato and Arabidopsis heat stress tolerance"

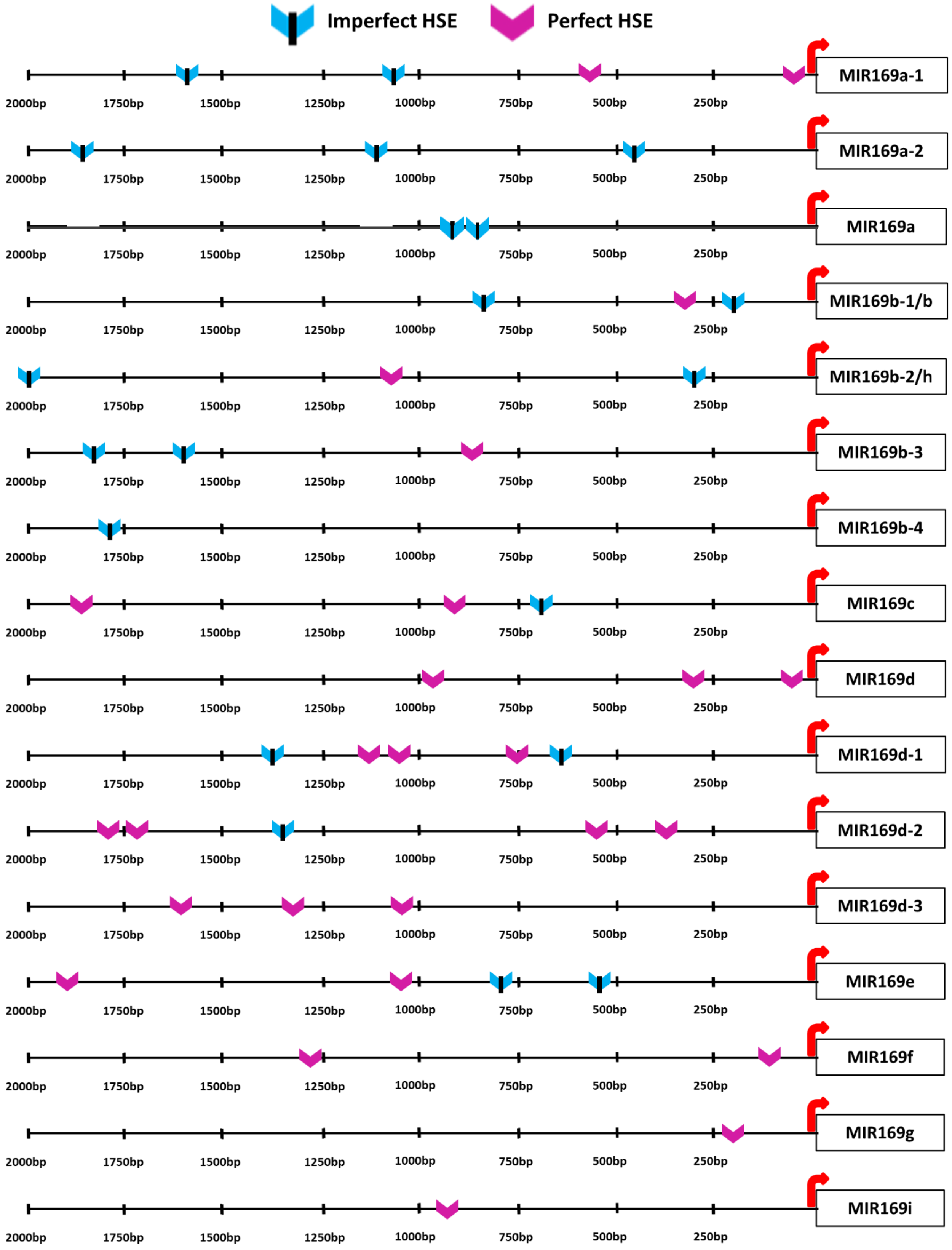


**Supplementary figure S1: Promoter analysis of *MIR169* genes in tomato.** Distribution of perfect and imperfect heat stress elements (HSE) in *MIR169* promoters of tomato. HSEs were identified manually by curating a list from published literature and marked as pink (perfect HSE) and turquoise blue (imperfect HSE) colored shapes on black colored lines representing *MIR169* promoters. The red right-handed arrow represents precursor start site. The HSE sequence variants for perfect and imperfect HSEs is provided in supplementary table S1.

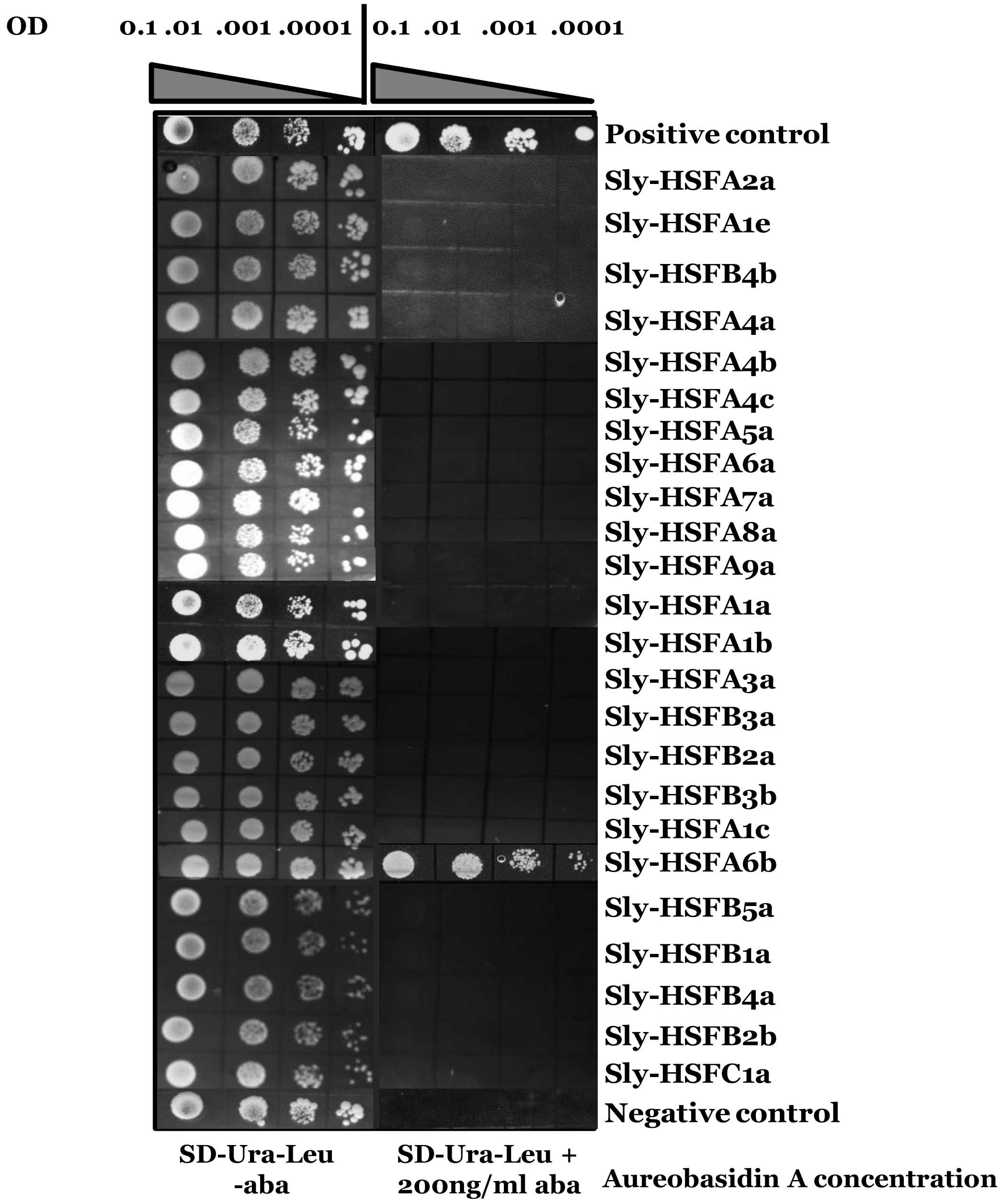


**Supplementary figure S2: Yeast-one-hybrid assay of 24 tomato HSFs on *Sly-MIR169d-1* promoter.** Interaction of only Sly-HSFA6b on *Sly-MIR169d-1* promoter highlights the specificity of the Y1H assay. The co-transformed Y1H gold yeast strain cultures were spotted (O.D. 0.1 to 0.0001) onto plates lacking URA and LEU with and without specific Aureobasidin A concentration and incubated for 3 days. Promoter of *sHSP* has been used as a positive control to show binding of HSFA3a on *sHSP* promoter (Li et al. 2013) and empty vectors as negative controls. The positive binding of HSFA6b on *Sly-MIR169d-1* promoter was assessed at 200 ng/ml Aureobasidin A concentration.

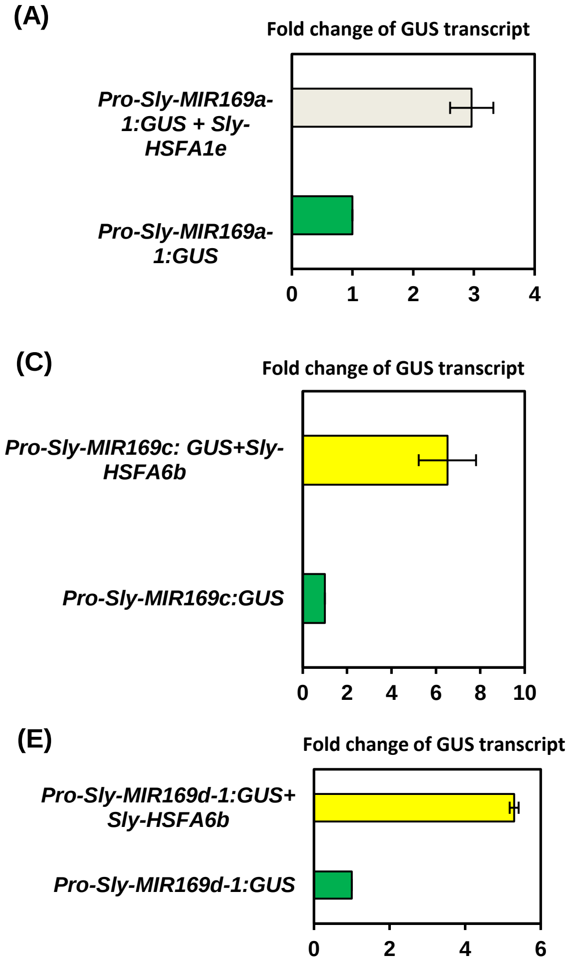


**Supplementary figure S3 : HSF-mediated transcriptional activation of tomato *MIR169* promoters.** *In-planta* transient assays showing HSF-mediated transcriptional activation of *Sly-MIR169:GUS* promoters in *Nicotiana benthamiana* as assessed by qRT-PCR. The fold-change of *GUS* transcripts were calculated using the 2−ΔΔCt method in (A) to (E). (A) Sly-HSFA1e with *pro-Sly-MIR169a-1:GUS*. (B) Sly-HSFA7a and Sly-HSFA1a with *pro-Sly-MIR169b:GUS*. (C) Sly-HSFA6b with *pro-Sly-MIR169c:GUS*. (D) Sly-HSFB1a with *pro-Sly-MIR169d:GUS*. (E) Sly-HSFA6b with *pro-Sly-MIR169d-1:GUS*. The *NPTII* gene co-expressed in the *pro-Sly-MIR169:GUS* construct was taken as the reference gene for normalization. The *GUS* expression from the reporter (*pro-Sly-MIR169:GUS*) infiltrated samples was set to one for normalization (marked as green bars). These experiments were repeated at least six times; the error bars represent standard deviation between the biological replicates.

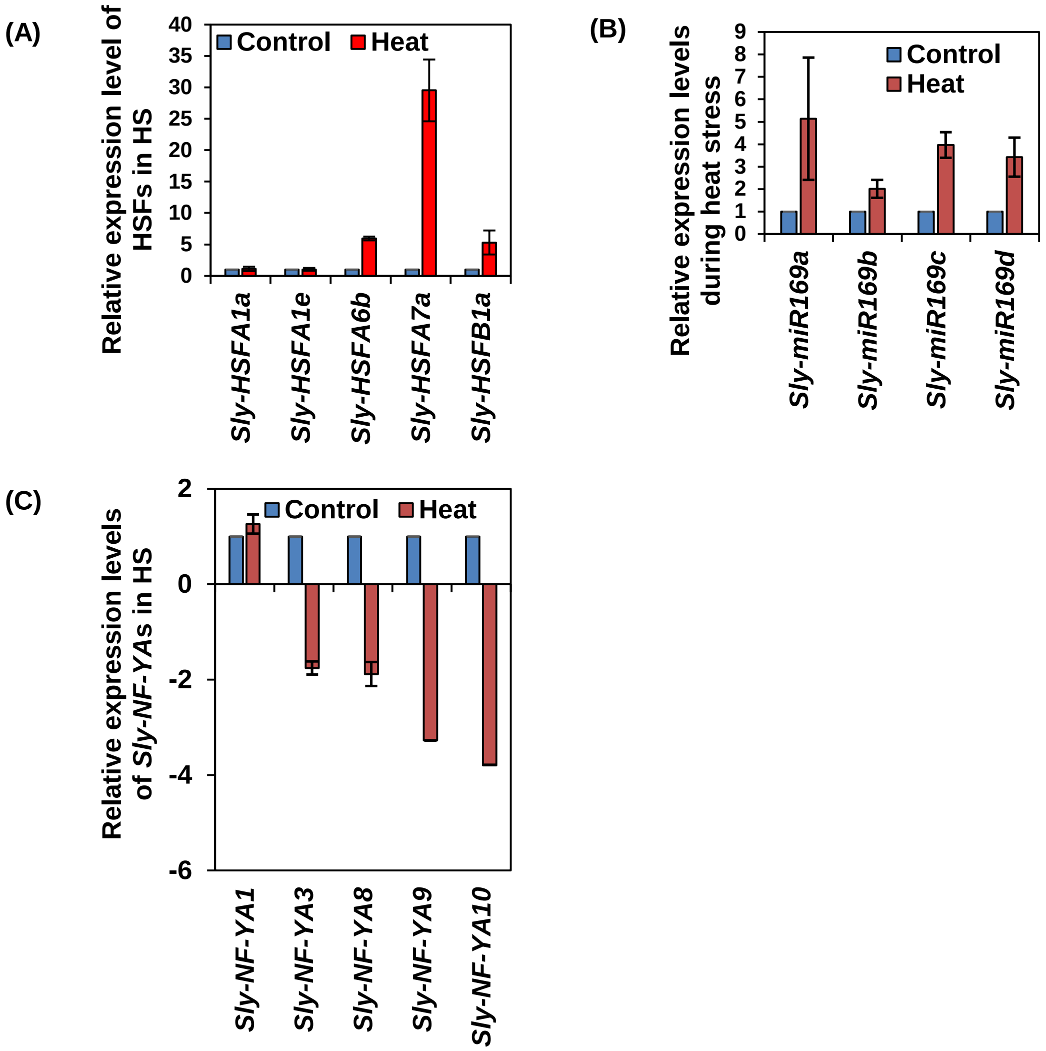


**Supplementary figure S4: Expression profiling of HSFs:miRNAs:NF-YAs during heat stress in tomato leaves.** (A) Expression profiles of *HSF* genes that regulate *Sly-MIR169* transcription in WT tomato leaves during control and heat stress condition by qRT-PCR using the 2−ΔΔCt method. (B) Expression of mature miR169s in response to heat stress as determined by taqman based qRT-PCR in control and heat challenged WT tomato leaves. (C) qRT-PCR based expression profiles of target *NF-YA* transcripts in WT control and heat stressed tomato leaves using the 2−ΔΔCt method. Data represents mean values and standard deviation of biological replicates. *Tubulin* was used as endogenous control in (A) and (C), while *Sly-U6snRNA* was used as normalization control in (B). Error bars in (A), (B) and (C) represent the standard deviation of biological replicates. The fold change normalization was done by setting the control value as one for all qRT-PCR expression analysis.

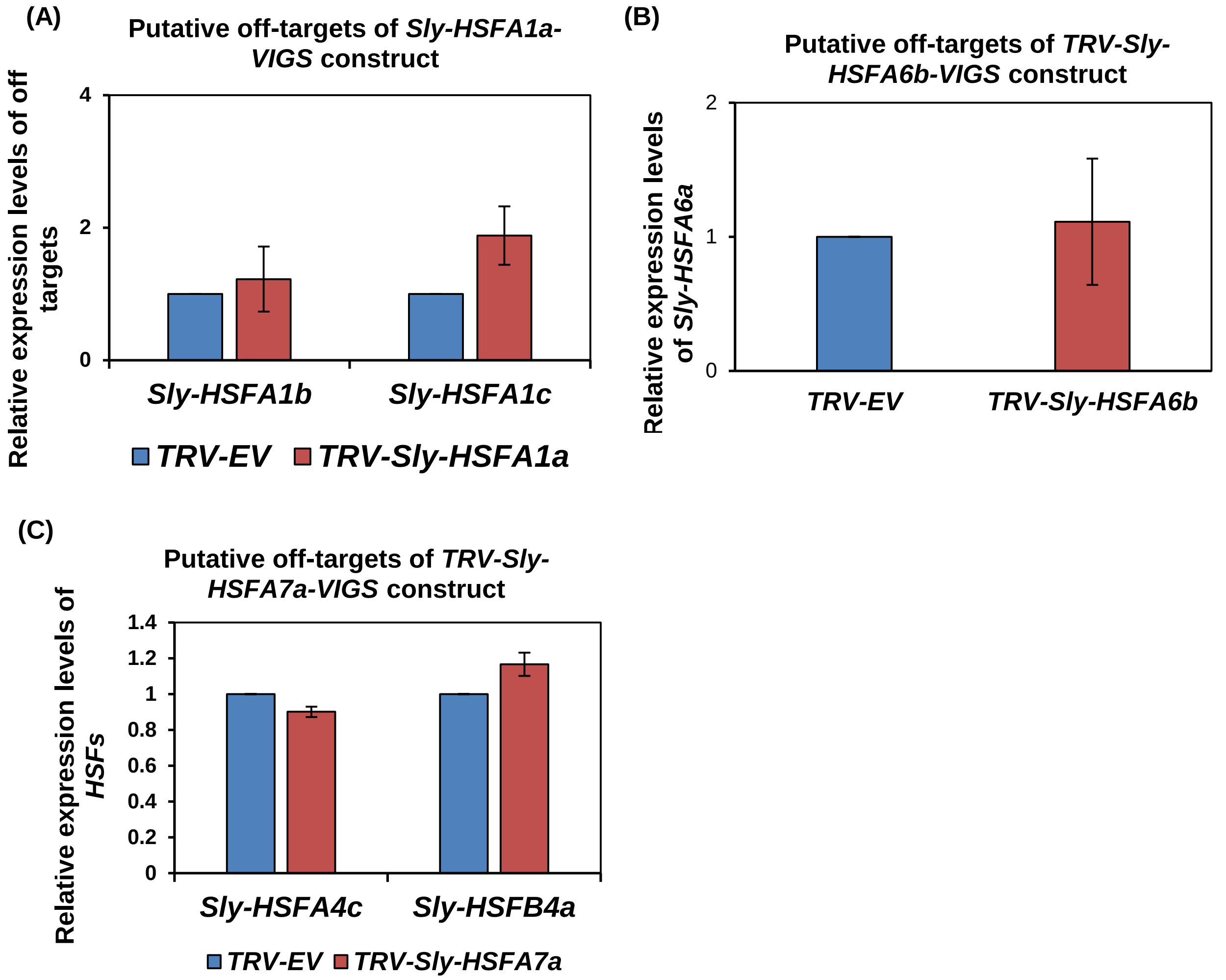


**Supplementary figure S5: Determining the specificity of HSFs silencing in VIGS silenced tomato plants.** (A-C) qRT-PCR based expression profiles of putative off target genes using the 2−ΔΔCt method. (A) Relative expression of Sly-*HSFA1b* and Sly-*HSFA1c* off targets in TRV-*Sly-HSFA1a* silenced plants. (B) Relative expression of Sly-*HSFA6a* off target in *TRV-Sly-HSFA6b* silenced plants. (C) Relative expression of Sly-*HSFA4c* and Sly-*HSFB4a* off targets in *TRV-Sly-HSFA7a* silenced plants. Error bars in (A-C) represent the standard deviation of biological replicates. *ACTIN* was used as endogenous control in A-C. Data normalization and fold change values with *TUBULIN* were presented in Supplementary table S3. The fold change normalization was done by setting the TRV-EV expression as one for all qRT-PCR expression analysis.

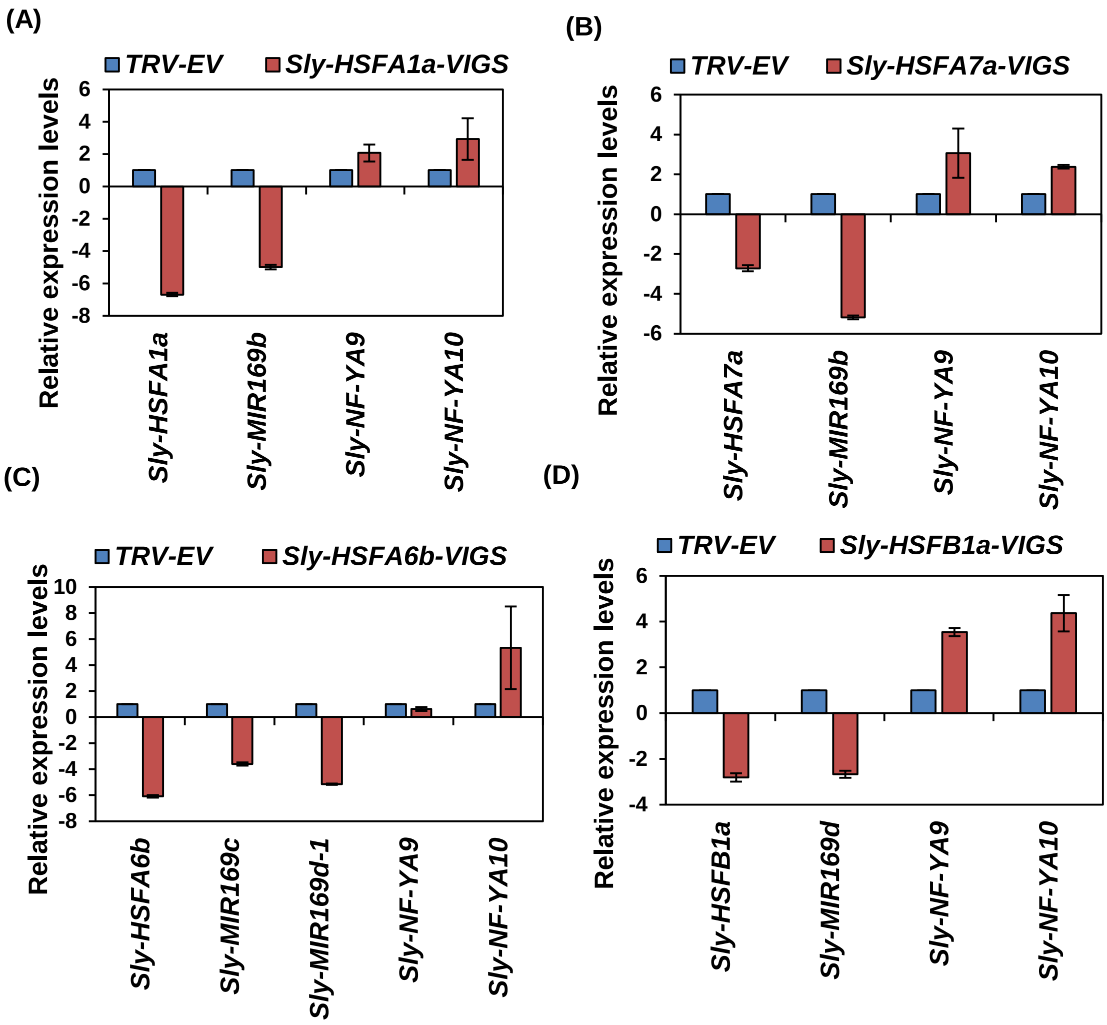


**Supplementary figure S6: In planta validation of heat governed HSFs mediated transcriptional regulation of MIR169s and NF-YAs in tomato leaves.**

The expression analysis of *HSF* genes, *Sly-MIR169* precursors and *NF-YA* transcripts in VIGS plants silenced for *Sly-HSFA1a* (A), *Sly-HSFA7a* (B), *Sly-HSFA6b* (C) and *Sly-HSFB1a* (D). 15-days-old tomato plants were agro-infiltrated with empty vector (TRV-EV) or TRV-Sly-HSF VIGS constructs. The VIGS established plants were subjected to heat stress after 3-weeks-of agro-infiltration, and used for expression studies of different genes. Graphical data represents mean values of expression of three to four biological sets. Error bars show standard deviation. To represent the negative fold-change on y axis, the relative expression values of down-regulated genes were transformed using the formula [-(1 / RQ ≤0.5)]. *Tubulin* was used as endogenous reference control. The fold change normalization was done by setting the control value as one for all qRT-PCR expression analysis.

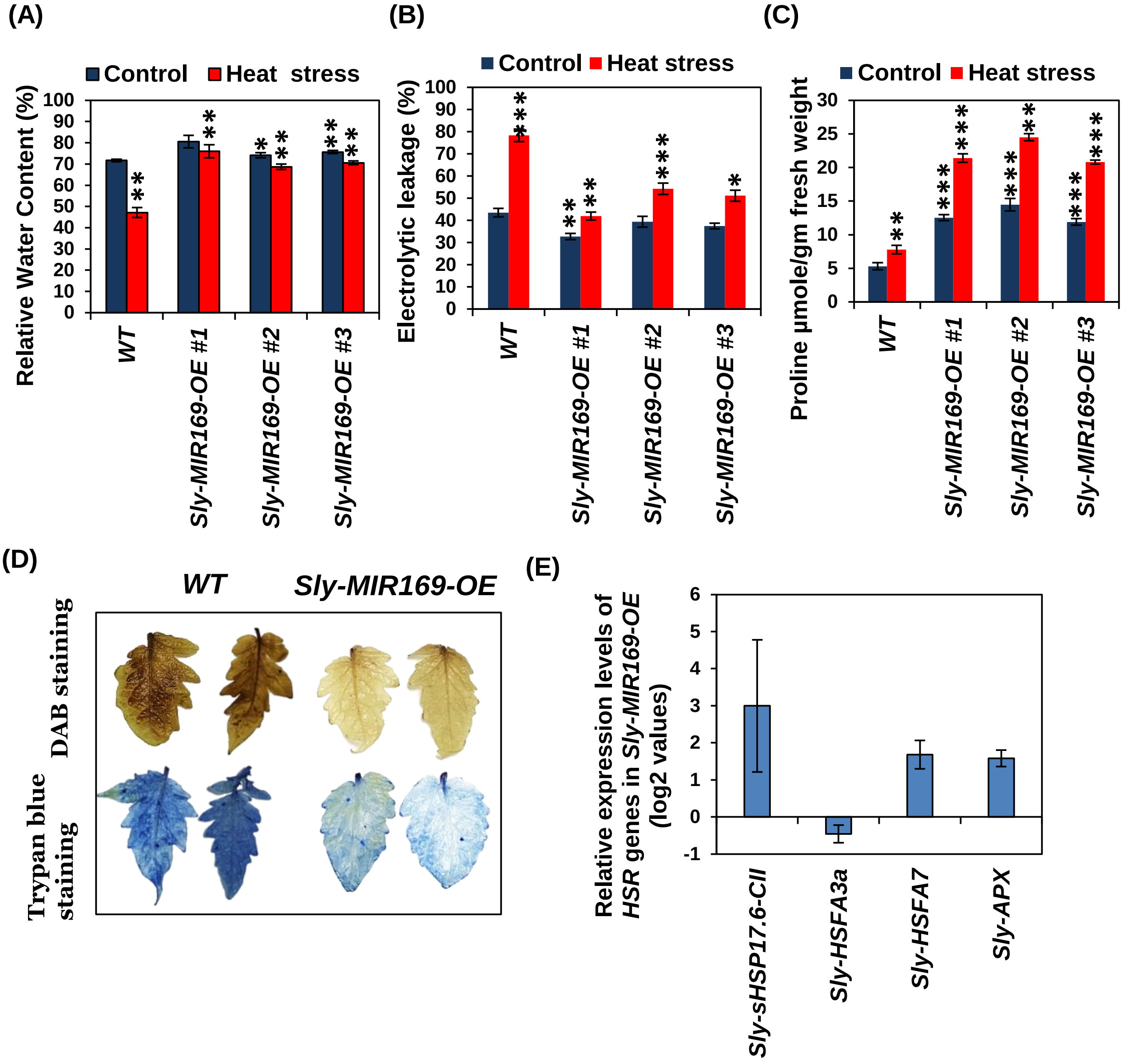


**Supplementary figure S7: Physiological, Biochemical and Molecular characterization of *Sly-MIR169-OE* lines.** (A) Estimation of relative water content in WT and *Sly-MIR169-OE* (three independent lines #1,2,3) transgenic plants during non-stressed and heat stressed conditions. (B) Quantification of electrolyte leakage in WT and *Sly-MIR169-OE* (three independent lines #1,2,3) transgenic plants during non-stressed and heat stressed conditions. (C) Estimation of proline content in WT and *Sly-MIR169-OE* (three independent lines #1,2,3) transgenic plants during non-stressed and heat stressed conditions. (D) DAB and trypan blue staining of WT and *Sly-MIR169-OE* transgenic plants during heat stressed conditions. (E) qRT-PCR based expression profiles of HSR genes in WT and *Sly-MIR169-OE* transgenic plants. Error bars represent the standard deviation of biological replicates in (A, B, C and E). The experiments were repeated at least three times with similar results, and data from one representative experiment are shown in (D). Error bars show standard deviation. *$p<0.05$, **$p<0.01$ and ***$p<0.001$ vs. wild type, by two-tailed Student's t-test.

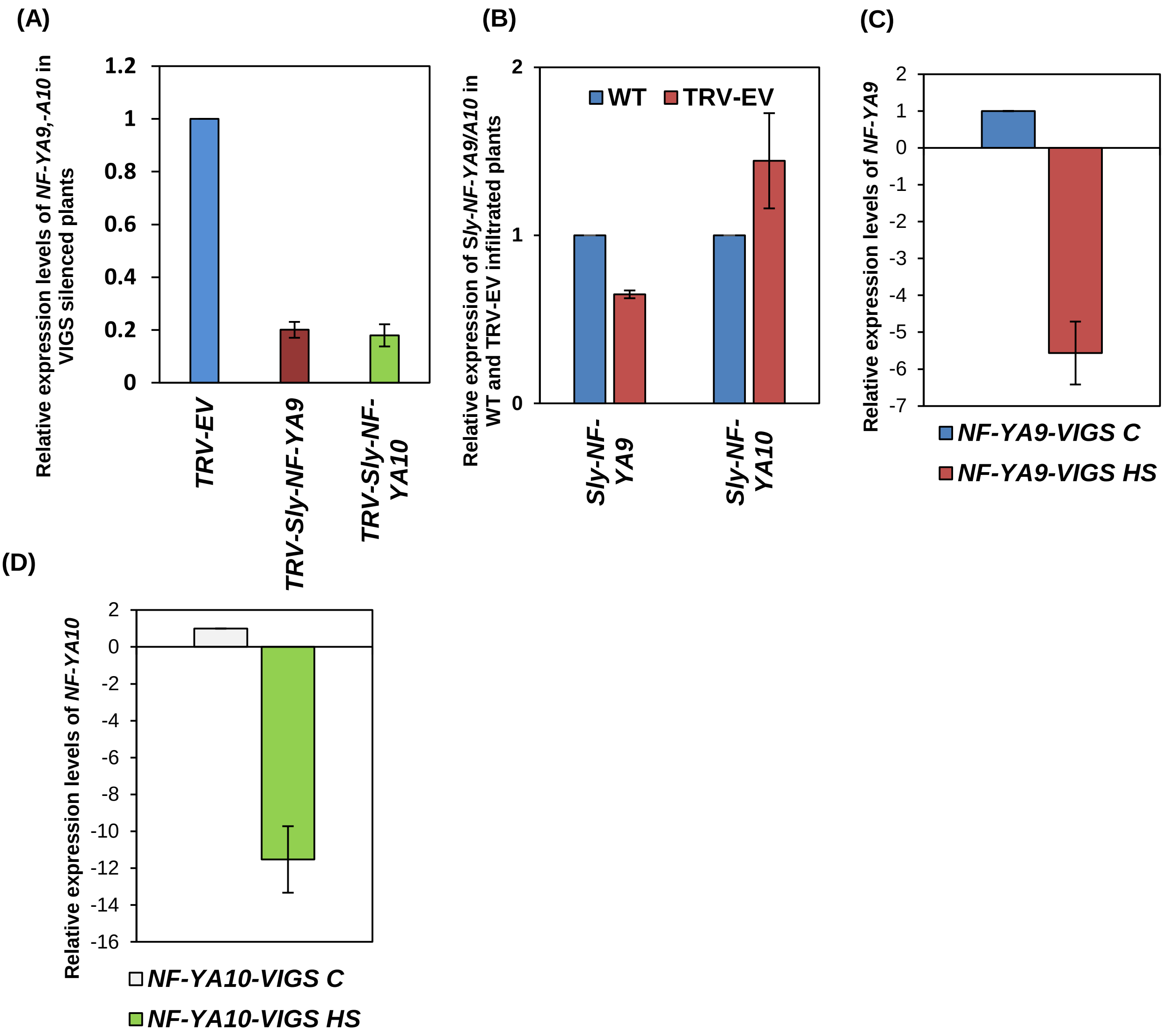


**Supplementary figure S8: Virus induced silencing of Sly-*NF-YA9* and Sly-*NF-YA10*.** (A) qRT-PCR analysis of Sly-*NF-YA9* and Sly-*NF-YA10* gene in TRV-EV, *TRV-Sly-NF-YA9* and *TRV-Sly-NF-YA10* plants in control (non-stressed) conditions confirming their silencing. The fold change normalization was done by setting the TRV-EV expression as one for all qRT-PCR expression analysis. (B) Expression of Sly-*NF-YA9* and Sly-*NF-YA10* in WT tomato and TRV-EV infiltrated plants in control (non-stressed) conditions. The fold change normalization was done by setting the WT control value as one for all qRT-PCR expression analysis (C-D) Expression profiles of *Sly-NF-YA9* (C) and *Sly-NFY-A10* (D) in *TRV-Sly-NF-YA9/A10* VIGS silenced plants in heat stress (HS) in comparison to control non-stressed conditions. The fold change normalization was done by setting the control value as one for all qRT-PCR expression analysis. The expression levels of genes were calculated using the 2−ΔΔCt method and presented using fold-change values. *Actin* was used for normalising expression in (A-D). Fold change values for (A-D) with *TUBULIN* normalisation were presented in Supplementary table S3. EV: vector control. Error bars represent the standard deviation of biological replicates.

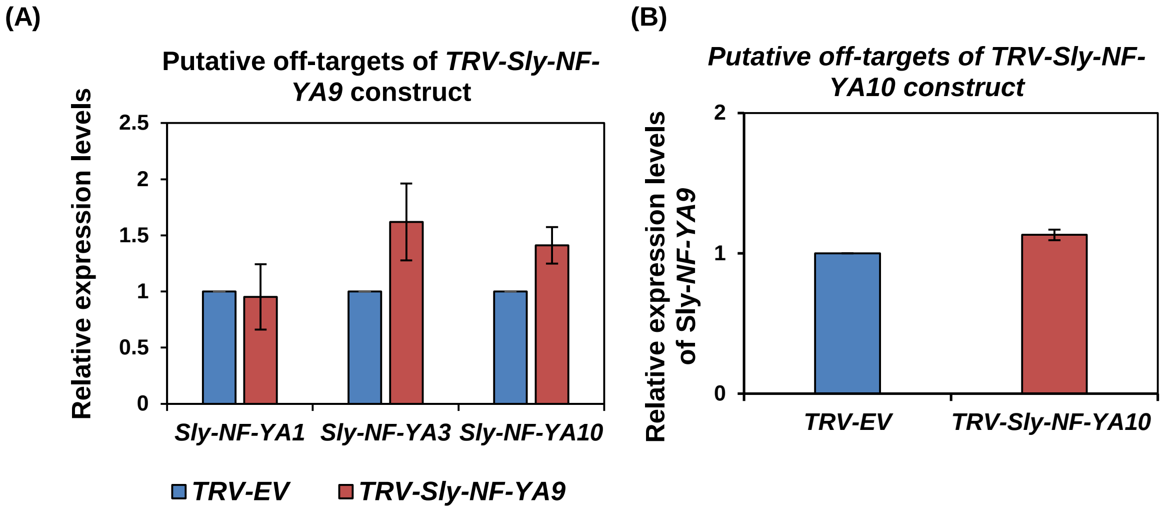


**Supplementary figure S9: Determining the specificity of NF-YA9/A10 silencing in VIGS silenced tomato plants.** (A-B) qRT-PCR based expression profiles of putative off target genes using the $2^{-\Delta\Delta Ct}$ method. (A) Relative expression of Sly-*NF-YA1, Sly-NF-YA3* and Sly-*NF-YA10* off targets in TRV-*Sly-NF-YA9* silenced plants. (B) Relative expression of *Sly-NF-YA9 (*no off targets were predicted for TRV-Sly-NF-YA10, we used *Sly-NF-YA9* to show specificity of silencing) as in *TRV-Sly-NF-YA10-VIGS* silenced plants. Error bars in (A-B) represent the standard deviation of biological replicates. *ACTIN* was used as endogenous control in A-B. Fold change values for (A-B) with *TUBULIN* normalisation were presented in Supplementary table S3. The fold change normalization was done by setting the TRV-EV expression as one for all qRT-PCR expression analysis.

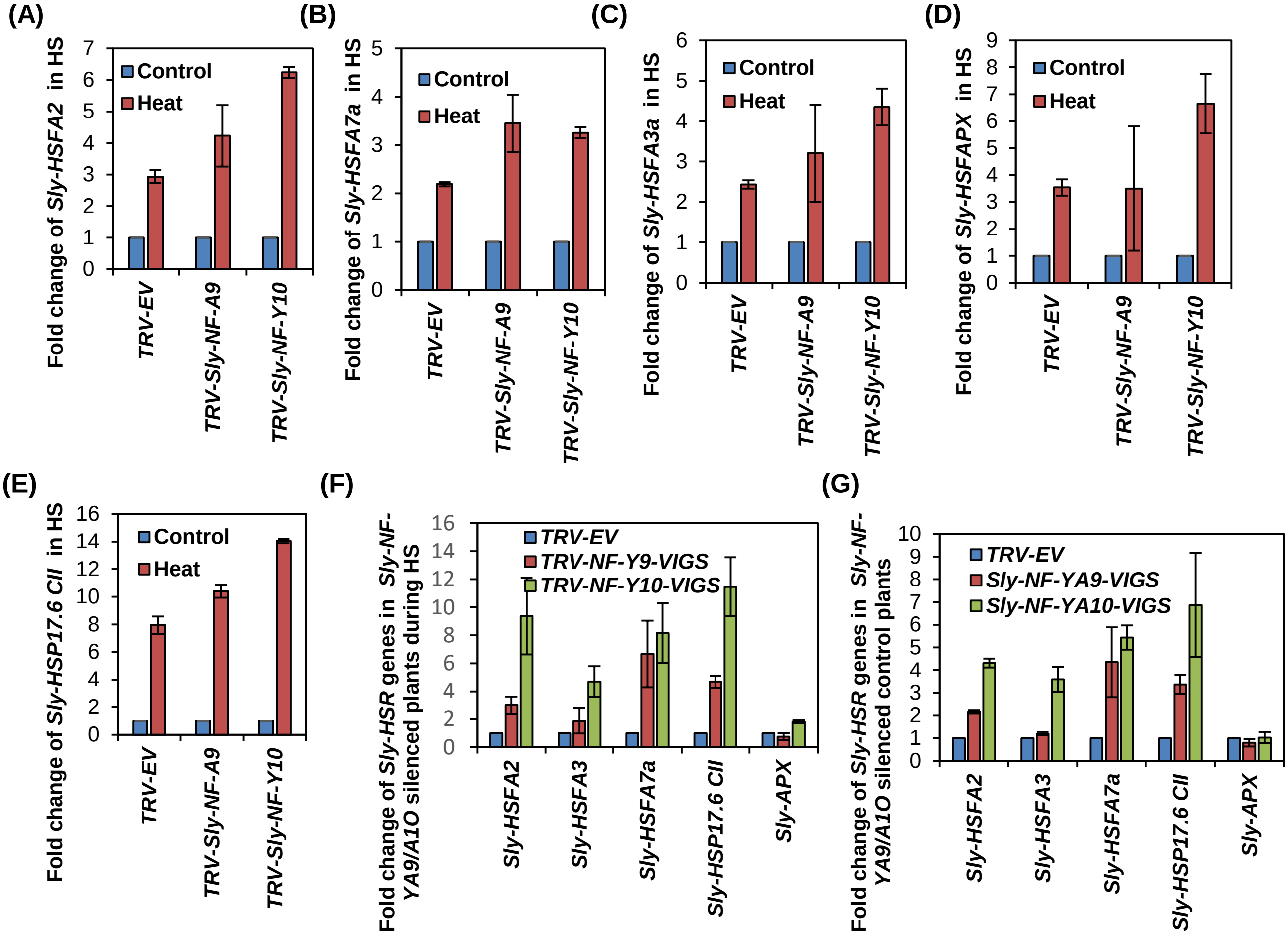


**Supplementary figure S10: Expression analysis of heat stress responsive genes in control and heat treated TRV-Sly-*NF-YA9* and TRV-Sly-*NF-YA10* silenced tomato plants.**

(A-E) Expression profiles of HSR genes in control and heat treated TRV-EV, *TRV-Sly-NF-YA9* and *TRV-Sly-NF-YA10* silenced plants. qRT-PCR based fold change in expression in control and heat stressed plants of *Sly-HSFA2* (A); *Sly-HSFA7a* (B); *Sly-HSFA3a* (C); *Sly-APX* (D) and *Sly-HSP17.6 CII*. (E). Relative expression of HSR genes in *TRV-Sly-NF-YA9* and *TRV-Sly-NF-YA10* silenced plants in control conditions as compared to *TRV-EV* plants. (G) Expression profiles of HSR genes in control and heat treated *TRV-Sly-NF-YA9* and *TRV-Sly-NF-YA10* silenced plants. The expression levels of genes were calculated using the $2^{-\Delta\Delta Ct}$ method and presented using fold-change values. *Tubulin* was used as reference gene for A-E and *Actin* was used as endogenous control for F-G. Fold change values for (F-G) with *TUBULIN* normalisation were presented in Supplementary table S3 The fold change normalization was done by setting the expression in control non-stressed as one for all qRT-PCR expression analysis in (A-E). The fold change normalization was done by setting the TRV-EV heat stressed expression as one for all qRT-PCR expression analysis in (F). The fold change normalization was done by setting the expression of TRV-EV in control non-stressed as one for all qRT-PCR expression analysis in (G). Error bars represent the standard deviation of independent biological replicates.

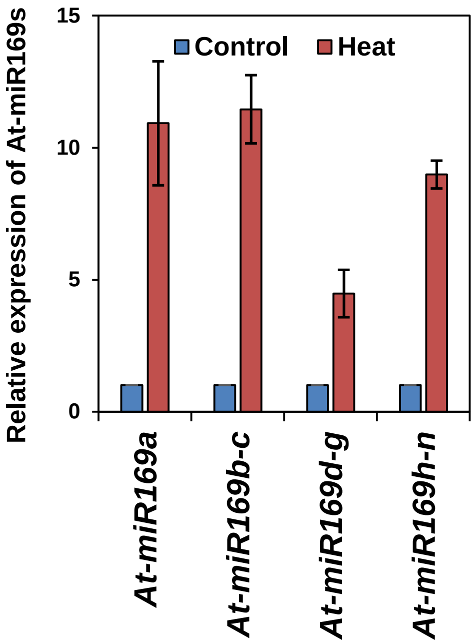


**Supplementary figure S11: Expression profiling of At-miR169s in heat stress.** Expression of mature At-miR169s in response to heat stress as determined by Taqman-based qRT-PCR. *U6snRNA* was used as the endogenous normalization control. The average values of multiple biological replicates are plotted as bars, error bars depict standard deviation between three replicates. The fold change normalization was done by setting the control value as one for all qRT-PCR expression analysis.

.

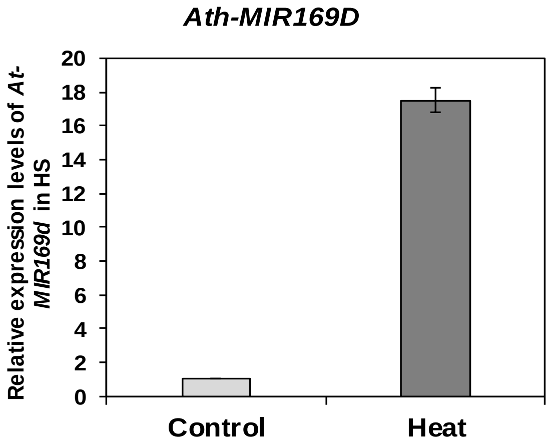


**Supplementary figure S12: Expression analysis of At-MIR169d in heat stress in Arabidopsis**. Expression levels of *At-MIR169d* in wild-type plants subjected to 0 (control) and 2 h heat stress at 42 °C. Data are shown as means ± SE of three biological replicates and normalized to *At-EF-α* reference gene using (2−ΔΔCT) method. The fold change normalization was done by setting the control value as one for the qRT-PCR expression analysis.

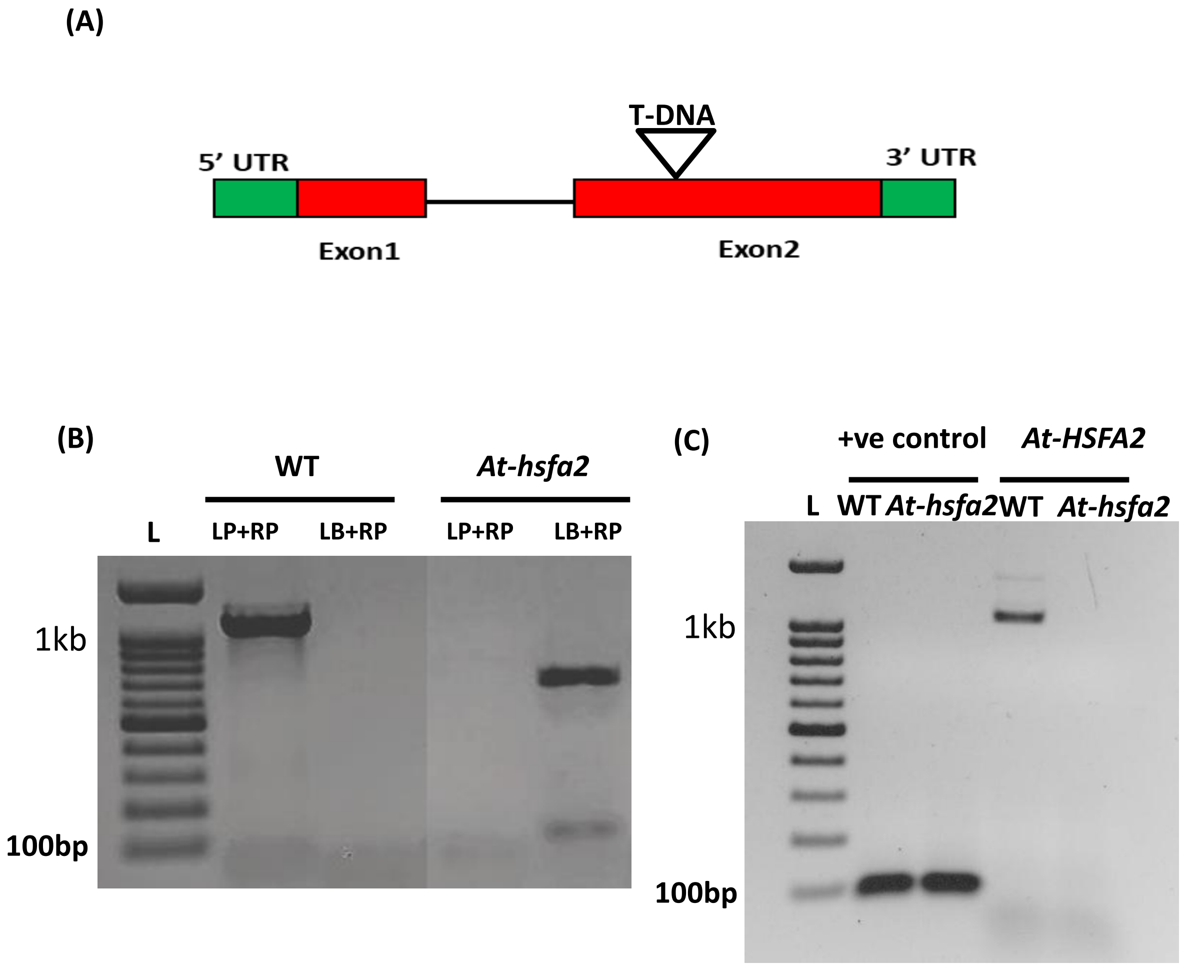


**Supplementary figure S13: Schematic structure of *At*-HSFA2 gene carrying the T-DNA insertion and lack of *At*-HSFA2 expression in the *At-hsfA2* mutant (Salk_008978).** (A) The two exons of *At*-HSFA2 genomic DNA are shown according to the gene features annotated at TAIR database. T-DNA insertion site is shown as inverted triangle in the second exon of *At*-HSFA2. The exon sequences are depicted by red and the UTRs are represented by green color. (B) Genotyping PCR of *At-hsfa2* mutants using genomic DNA, presence of border specific amplification in *At-hsfa2* confirms their homozygosity, while presence of only gene specific amplification in control Columbia plants confirms their wild type nature. (C) Expression of *At-HSFA2* was determined by RT-PCR, using full length cDNA from WT and mutant *At-hsfa2* plants. Absence of *At-HSFA2* specific band in mutant plants confirms the knockout nature of *At-hsfa2* homozygous mutants. RNA was isolated from detached mature leaves of the wild-type (wt) or *At-hsfa2* plants. Expression of *actin* gene is shown as a loading control.

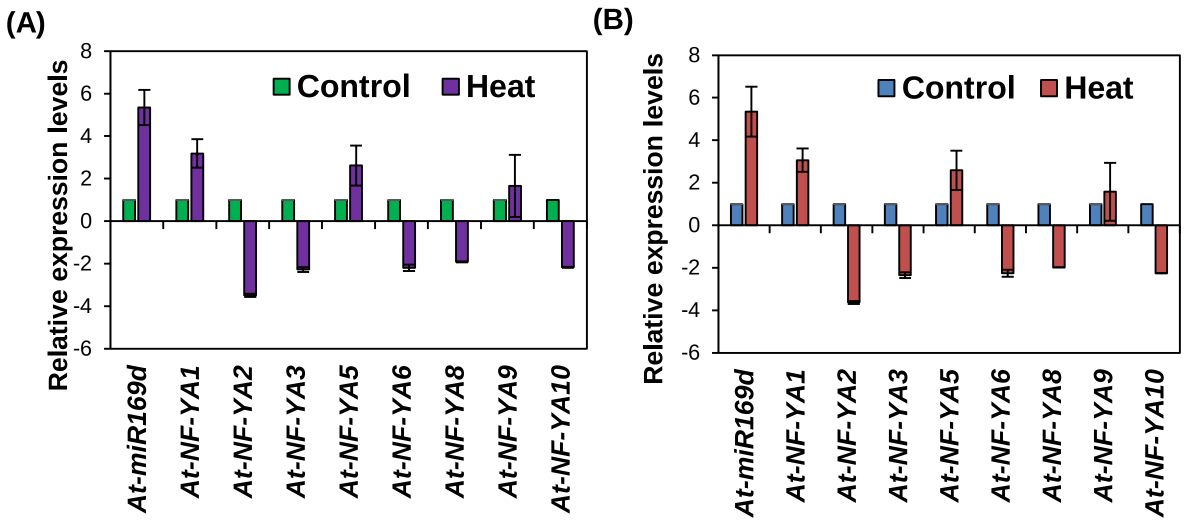


**Supplementary figure S14: Expression pattern of miRNA169d:NF-YA in Arabidopsis.** (A) Expression levels of mature miR169d and eight miR169 target genes in wild-type plants subjected to 0 (control) and 2 h heat stress at 42°C using qRT-PCR. Data are shown as means ± SD of biological replicates and normalized to *actin* and *5SrRNA* reference genes, for mRNA and miRNA respectively, using ($2^{-\Delta\Delta CT}$) method for A and with *At-EF-α* reference gene and *U6snRNA* for mRNA and miRNA respectively, using ($2^{-\Delta\Delta CT}$) method for B. The fold change normalization was done by setting the control value as one for all qRT-PCR expression analysis.

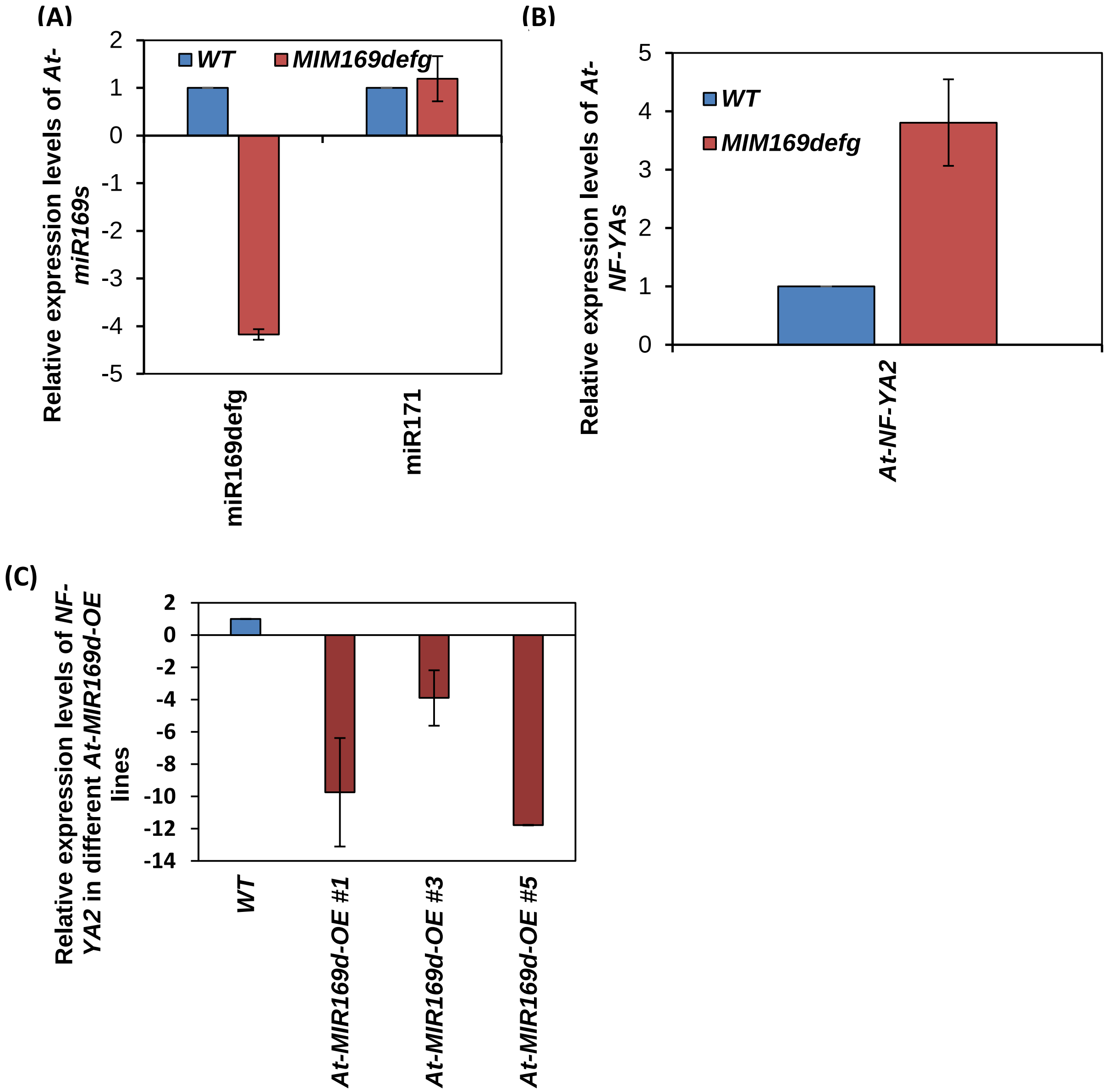


**Supplementary figure S15: Establishing miRNA169:target modules operating in Arabidopsis**. (A) Taqmann based qRT- qPCR analysis of the accumulation of mature miR169defg and miR171bc (used as negative control) in Arabidopsis MIM169defg lines. Average values of expression for different miR169s in three independent lines of MIM169defg transgenic plants were plotted as bars and error bars deoicts the standard deviation between the replicates. (B) qRT-PCR analysis of the accumulation of the *At-NF-YA2* transcripts in MIM169defg Arabidopsis lines. Data are presented as average fold induction relative to control where induction is the ratio of transcript levels in two independent MIM169defg transgenic lines to those in wild-type plants (WT). Error bars, ± SD of three repeats. (C) qRT-PCR analysis of the reduction of *At-NF-YA2* transcripts in three independent miR169d over expressing (OE) lines. Data are presented as average fold in two independent *MIM169defg* lines relative to WT control, that was set as one.

(A)

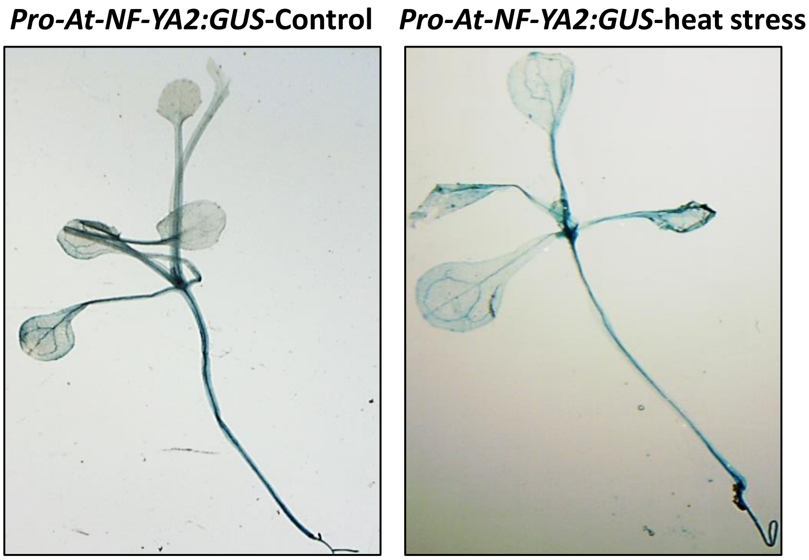


(B)

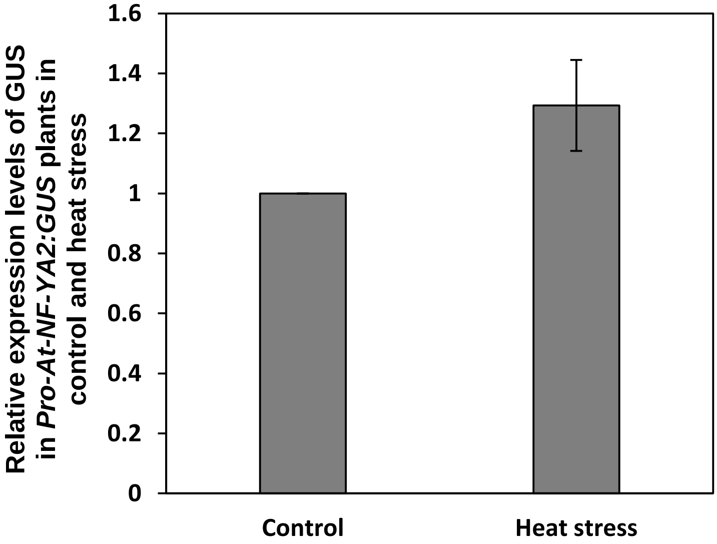


**Supplementary figure S16: Heat stress mediated transcriptional regulation of *At-NF-YA2*.** (A) Histochemical GUS expression of Pro-*At-NF-YA2:GUS* transgenic Arabidopsis plants during control and heat stress. (B) qRT-PCR based determination of *GUS* transcripts of Pro-*At-NF-YA2:GUS* transgenic Arabidopsis plants during control and heat stress. These experiments were repeated 3 times with 20 seedling per replicate. Error bars depict standard deviation between three replicates. The fold change normalization was done by setting the control value as one for qRT-PCR expression analysis.

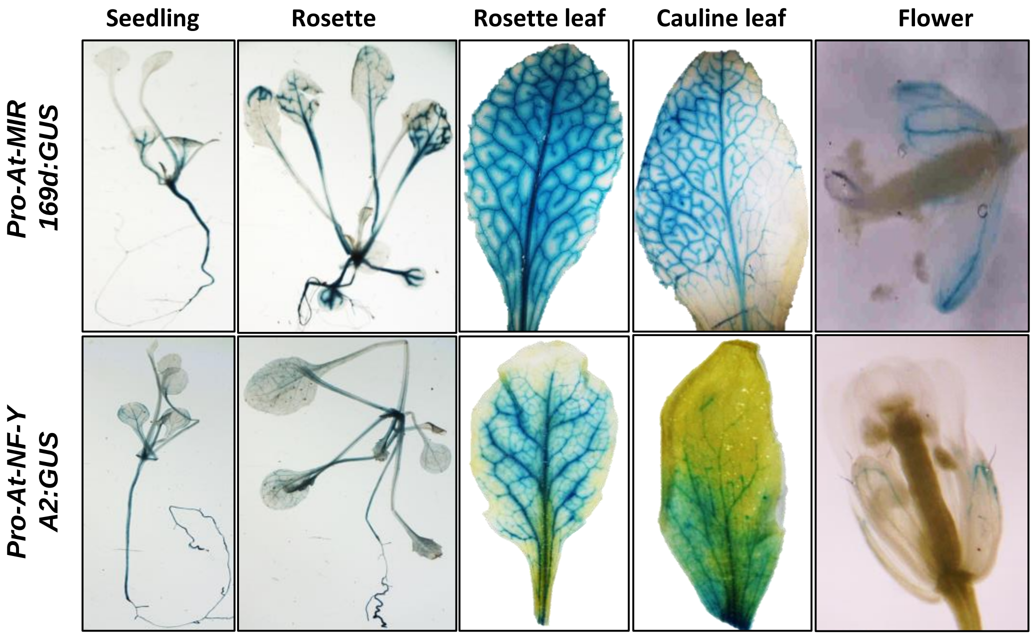


**Supplementary figure S17: Delineating the existence and localization of miR169d:At-NF-YA2 functional module in Arabidopsis.** Histochemical GUS expression patterns of *Pro-At-MIR169d:GUS and Pro-At-NF-YA2:GUS* promoter reporter transgenic Arabidopsis plants in: seedling (2 weeks old), complete rosette (4 weeks), mature rosette leaf, cauline leaf and flower

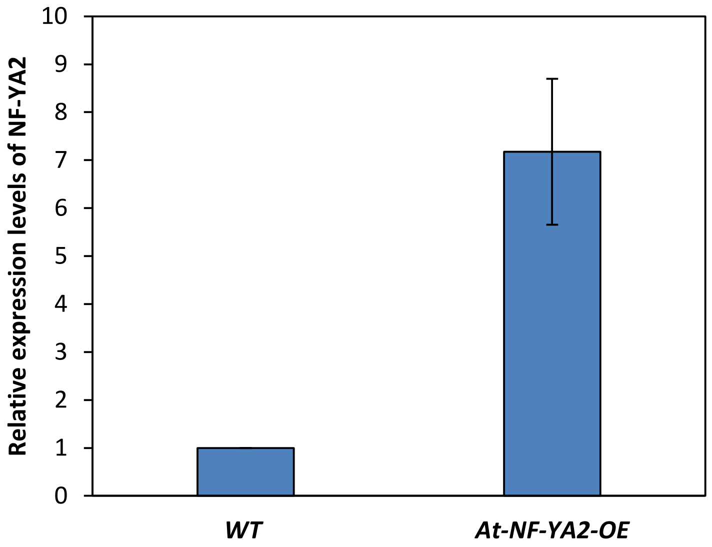


**Supplementary figure S18: Expression levels of *At-NF-YA2* in *At-NF-YA2* overexpressing transgenic plants.** Relative expression of *At-NF-YA2* in transgenic Arabidopsis plants overexpressing *At-NF-YA2* under constitutive CaMV35S promoter. Bar represents the average data of three biological replicates and error bar represents standard deviation of the replicates.

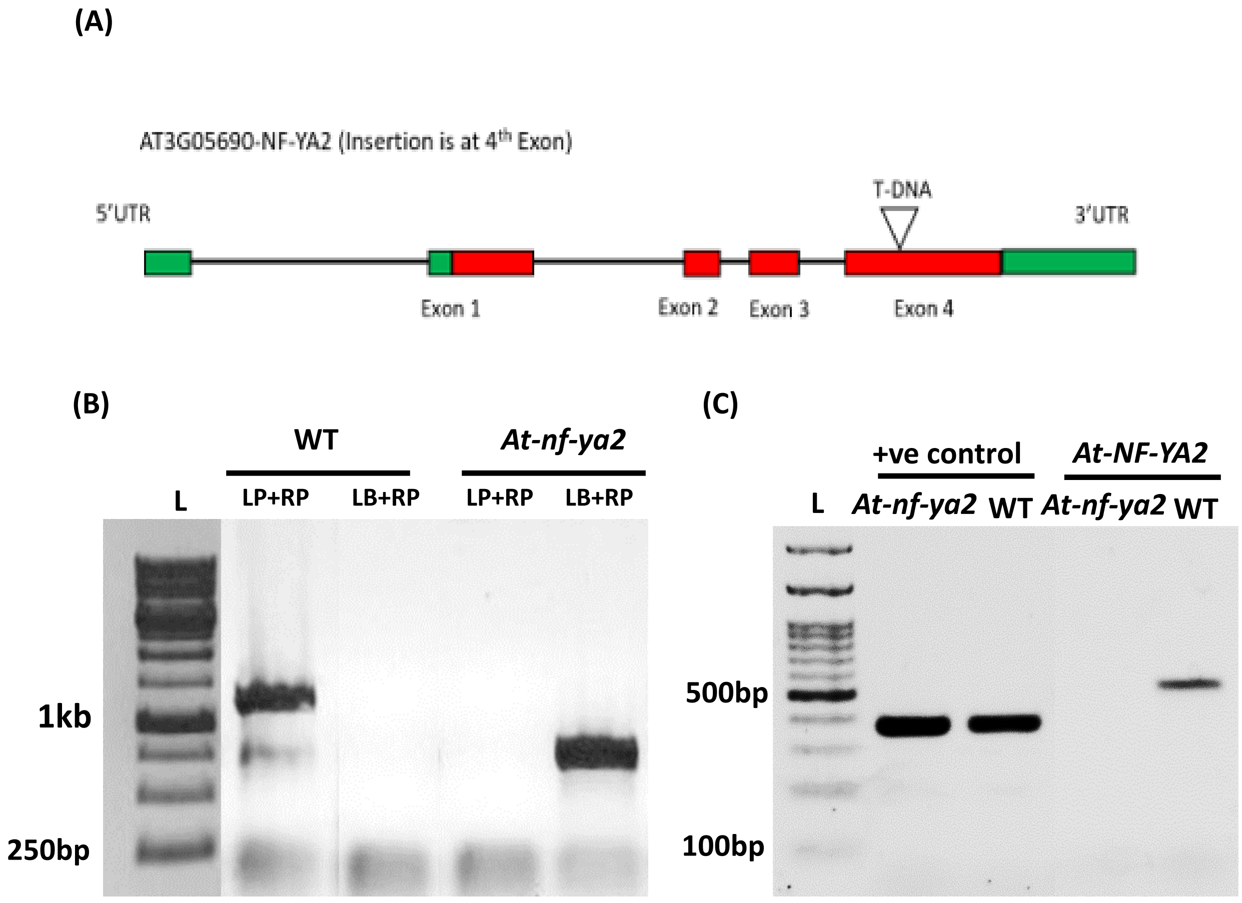


**Supplementary figure S19: Schematic structure of *At-NF-YA2* gene carrying the T-DNA insertion and lack of *At-NF-YA2* expression in the *At-nf-ya2* mutant (SALK_021228).** (A) The four exons of *At-NF-YA2* genomic DNA are shown according to the gene features annotated at TAIR database. T-DNA insertion site is shown by inverted triangle in the fourth exon of *At-NF-A2*. The exon sequences are depicted by red and the UTR are represented by green color. (B) Genotyping PCR of *At-nf-ya2* mutants, presence of border specific amplification in *At-nf-ya2* plants genomic DNA confirms their homozygosity, while presence of only gene specific amplification in control Columbia plants confirms their wild type nature. (C) Expression of *At-NF-YA2* was determined by RT-PCR, using full length cDNA from WT and mutant *At-nf-ya2* plants. Absence of *At-NF-YA2* specific band in mutant plants confirms the knockout nature of *At-nf-ya2* homozygous mutants. RNA was isolated from detached mature leaves of the wild-type (WT) or *At-nf-ya2* plants. Expression of actin is shown as a loading control.

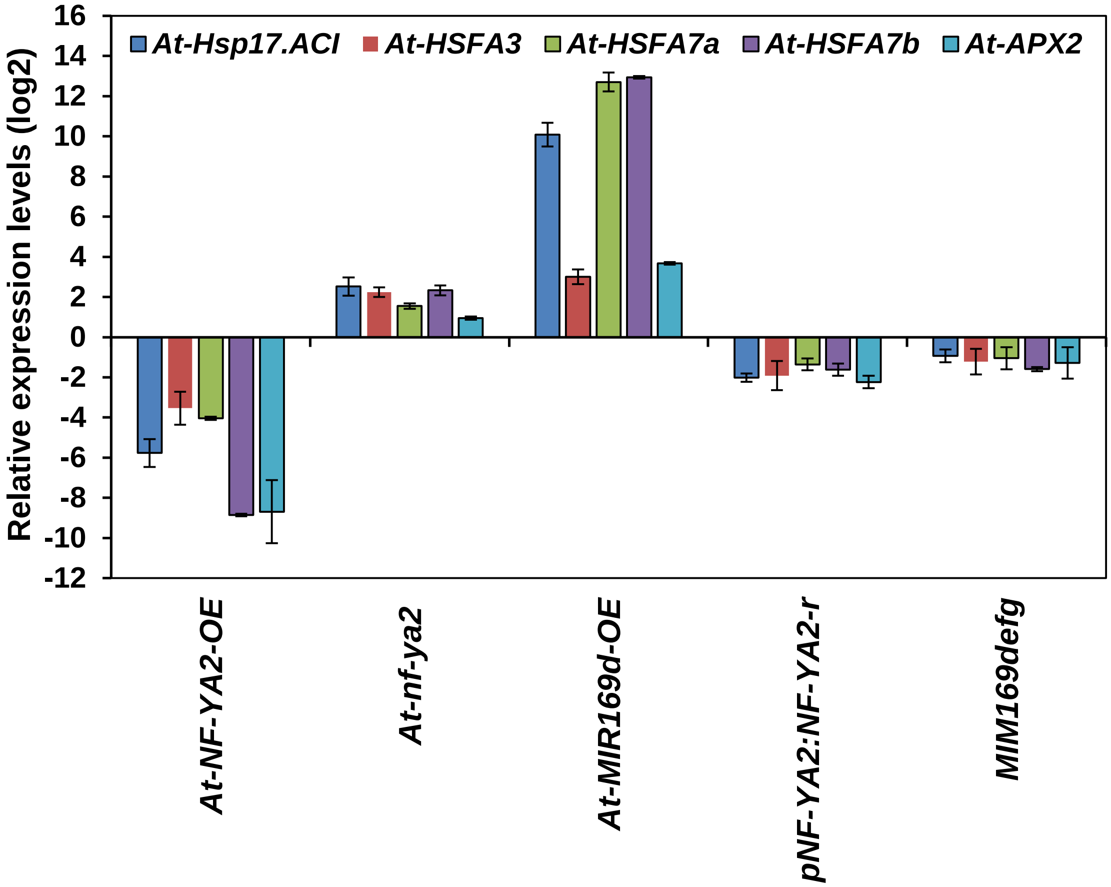


**Supplementary figure S20: Expression profiling of HSR genes in different transgenic lines of Arabidopsis.** Expression patterns of five heat stress-responsive genes in WT, *At-MIR169d-OE*, *At-NF-YA2-OE*, *At-nf-ya2, pNF-YA2:NF-YA2*-r and MIM169defg plants. Data are shown as means ± SD of biological replicates and normalized to *At-EF-α* reference gene using ($2^{-\Delta\Delta CT}$) method. The fold change normalization was done by setting the control value as one for qRT-PCR expression analysis.

.

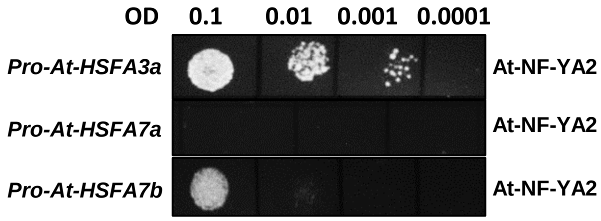


**Supplementary figure S21: Yeast-one-hybrid assay based transcriptional regulation of *At-HSFA3a, At-HSFA7a* and *At-HSFA7b* promoters by At-NF-YA2**. Positive interaction of At-NF-YA2 with the promoters of *At-HSFA3a* and *At-HSFA7b* were obtained. Serial dilutions (O.D. 0.1 to 0.0001) of yeast cultures were spotted onto plates lacking LEU with specific Aureobasidin A concentration and incubated for 3 days.

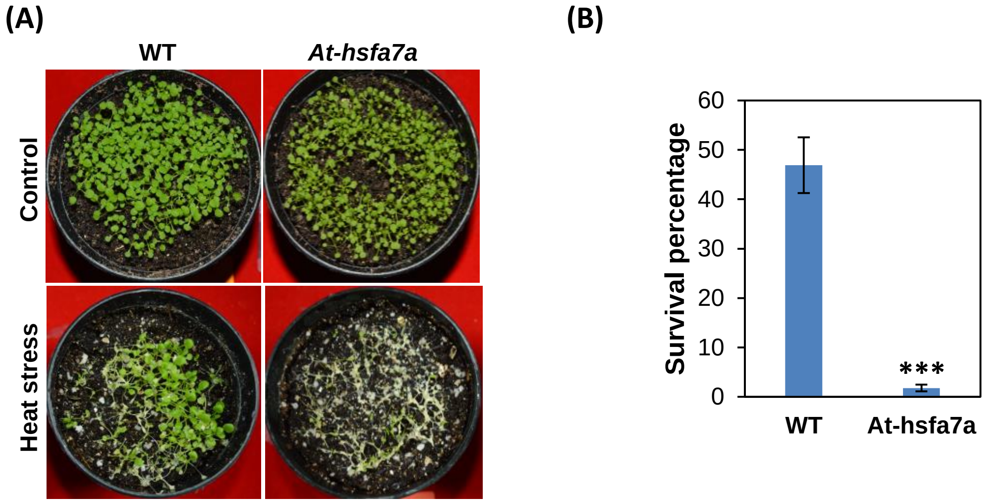


**Supplementary figure S22: Thermotolerance assay of WT and *At-hsfa7a* plants**. (A) Two-weeks-old soil-grown WT and *At-hsfa7a* mutant plants were subjected to 0 (control) or 2h heat stress at 42°C, and damage was recorded 6 days later. (B) Estimation of survival (percentage) of heat stress treated WT and *At-hsfa7a* plants. Plants were assayed for heat stress tolerance after 6 days of HS. These experiments were repeated at least three times with similar results, and average data of all experiments are shown. Error bars, ± SD of three repeats. ***p<0.001 vs. wild type, by two-tailed Student's t-test.

**(A)**

```
>Solyc01g087240.3.1         Sly-NF-YA9
Length=303

 Score = 105 bits (263),  Expect = 1e-26, Method: Compositional matrix adjust.
 Identities = 75/199 (38%), Positives = 106/199 (53%), Gaps = 33/199 (17%)

At-NF-YA2   18   WWTAFG--SQPLAPESLAGDSDSFAGVKVGSVGETGQRVDKQSNSATHLAFSLGDVKSPR  75
                 WW  +G  + PLA E+      S  G +           DK++ +    + S G+ +  +
Sly-NF-YA9  29   WWHGYGDNTMPLASENAVAQEKSEGGNQ-----------DKETKALAMESGSDGNNEQYK  77

Query  76   LVPKPHGATFS--MQSPCLELGFSQPPIYTKYPYGEQQYYGVVSAYGS----------QS  123
               K    T +  M     EL       +    YPY + QY G+++ YG+           +
Sbjct  78   QHLKHFAPTTAAIMAEQQKEL-TGHSAMLASYPYPDMQYGGMMT-YGAPVHPHLFEIHHA  135

Query  124  RVMLPLNMETEDSTIYVNSKQYHGIIRRRQSRAKAAAVLDQKKLSSRCRKPYMHHSRHLH  183
            R+ LPL+ME E   +YVN+KQYHGI+RRRQ RAKA   L++K +  + RKPY+H SRH H
Sbjct  136  RMPLPLDMEEE--PVYVNAKQYHGILRRRQIRAKAE--LERKAI--KARKPYLHESRHQH  189

Query  184  ALRRPRGSGGRFLNTKSQN  202
            A+RR RG+GGRFLNTK  N
Sbjct  190  AMRRARGTGGRFLNTKKLN  208
```

**(B)**

```
>Solyc01g006930.3.1         Sly-NF-YA10
Length=311

 Score = 145 bits (367),  Expect = 9e-42, Method: Compositional matrix adjust.
 Identities = 117/323 (36%), Positives = 168/323 (52%), Gaps = 70/323 (22%)

At-NF-YA2    10   LFSAPQTSWWT-AFGSQPLAPESLAGDSDSFAGVKVGSVGETGQRVDKQSNSATHLAFSL  68
                  L SAP   WW+  F SQ +A       ++ F  +K  SV +       +   +AT    S
Sly-NF-YA10  22   LCSAP---WWSNGFMSQSVA------YAEPFGQLKSASVEQ------QPKGNATEFTISS  66

Query  69   GDVKSP---RLVPKPHGATFSMQSPCL------ELGFSQPPIYTKYPYGEQQYYGVVSAY  119
            GD KS    + +P    A+ S+++  +      ELGF Q  IY K+PYGEQ   G+ SAY
Sbjct  67   GDCKSSANGQKLPNIQAAS-SVRAANMDYRGHFELGFGQSLIYAKHPYGEQ-CIGLFSAY  124

Query  120  GSQ--SRVMLPLNMETEDSTIYVNSKQYHGIIRRRQSRAKAAAVLDQKKLSSRCRKPYMH  177
              Q   R+MLPLN+ +++  I+VN+KQYHGI+RRR++RAK     + +K + + RKPY+H
Sbjct  125  APQLSGRIMLPLNLASDEGPIFVNAKQYHGILRRRKTRAK-----EMEKKALKPRKPYLH  179

Query  178  HSRHLHALRRPRGSGGRFLNTKSQN--LENSGTNAKKGDGSMQIQSQPKPQQSNSQNSEV  235
             SRHLHALRRPRG GGRFLNT++ N  ++   TN    G +Q         + SQNSEV
Sbjct  180  LSRHLHALRRPRGCGGRFLNTRNMNGTMKAGKTNNMFKTGDVQ-----NFYPTGSQNSEV  234

Query  236  VHPENGTMNLSNGLNVSGSEV------------TSMNYF----LSSPVHSL-------GG  272
            +  +  + NLS+    SGS              +++ F    L  PV ++       G
Sbjct  235  LQSD--SSNLSSPKETSGSRFFDSSGVANMYSSDNLDPFLFQNLRPPVQAIPDMMNTGHG  292

Query  273  MVMPSKWIAAAAAMDNGCCNFKT  295
            + +  KW+  A    + CCN K
Sbjct  293  IFVSGKWVCTA----DSCCNLKV  311
```

**Supplementary figure S23: At-NF-YA2, Arabidopsis orthologue of tomato Sly-NF-YA9 and Sly-NF-YA10.** (A-B) Protein sequence alignment of tomato Sly-NF-YA9 (A) and Sly-NF-YA10 with At-NF-YA2. Protein sequences were aligned with Arabidopsis proteome at TAIR database by using Blastp tool.

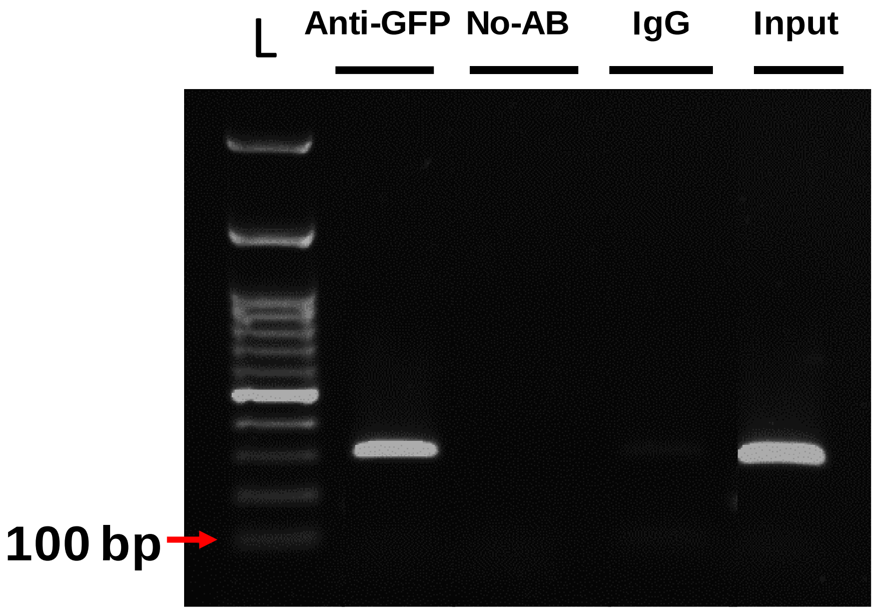


**Supplementary figure S24:** Binding of *Sly-HSFA6b* to Sly-MIR169d-1 promoter. Absence of any amplification in no-Ab (no antibody) and slight amplification in IgG control confirms the specificity of immunoprecipitation. Presence of strong amplification on input DNA confirms the good amplification efficiency of the primer used for analysis. Presence of amplification in Anti-GFP lane confirms the enrichment of Sly-MIR169d-1 promoter in *Sly-HSFA6b:GFP* CHIP. L denotes the 100 bp ladder.

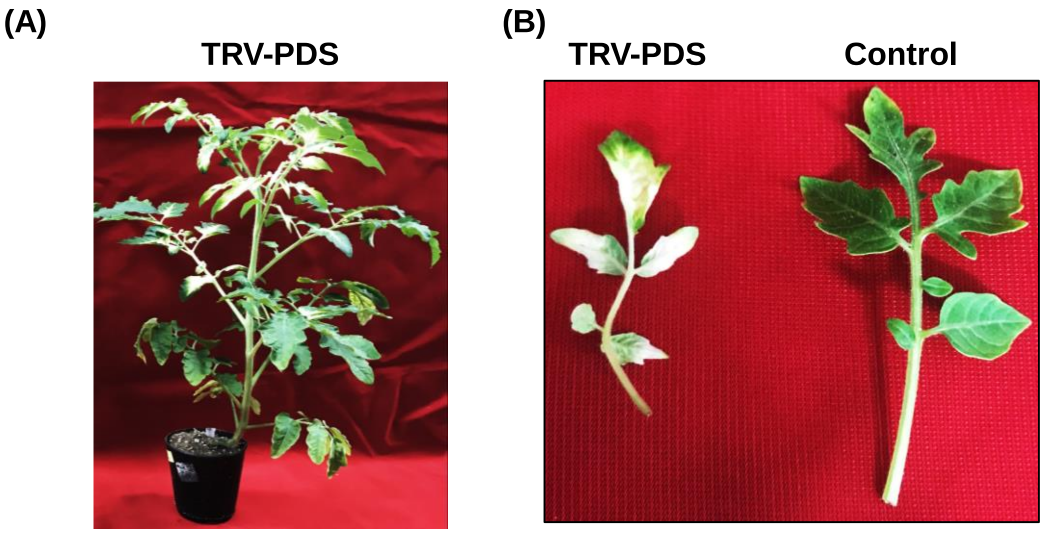


**Supplementary figure 25: Virus induced silencing of tomato *Phytoene desaturase* (*PDS*) gene.** *PDS* silencing was used as a control for assessing establishment of gene silencing by VIGS. (A) Silencing of the PDS control gene causing photobleaching in tomato plants 3 weeks after agroinfection. (B) Enlarged individual leaves of TRV-PDS and control WT leaves.
